# Pathogen profiling of Australian rabbits by metatranscriptomic sequencing

**DOI:** 10.1101/2022.02.15.480619

**Authors:** Maria Jenckel, Robyn Hall, Tanja Strive

## Abstract

Australia is known for its long history of using biocontrol agents, like myxoma virus and rabbit haemorrhagic disease virus (RHDV), to manage wild European rabbit populations. Interestingly, while undertaking RHDV surveillance of rabbits that were found dead we observed that approximately 40% of samples were negative for RHDV. To investigate whether other infectious agents are responsible for killing rabbits in Australia we subjected a subset of these RHDV-negative liver samples to metatranscriptomic sequencing. In addition, we investigated whether the host transcriptome data could provide additional differentiation between likely infectious versus non-infectious causes of death. We identified transcripts from several *Clostridia* species, *Pasteurella multocida, Pseudomonas*, and *Eimeria stiedae* in liver samples of several rabbits that had died suddenly, all of which are known to infect rabbits and are capable of causing fulminant disease. In addition, we identified *Hepatitis E virus* and *Cyniclomyces* yeast in some samples, both of which are not usually associated with severe disease. In one third of the sequenced liver samples, no infectious agent could be identified. While metatranscriptomic sequencing cannot provide definitive evidence of causation, additional host transcriptome analysis provided further insights to distinguish between pathogenic microbes and commensals or environmental contaminants. Interestingly, three samples where no pathogen could be identified showed evidence of upregulated host immune responses, while immune response pathways were not upregulated when *E. stiedae, Pseudomonas*, or yeast were detected. In summary, although no new putative rabbit pathogens were identified, this study provides a robust workflow for future investigations into rabbit mortality events.

**Importance:** We have observed that approximately 40% of rabbit liver samples submitted for RHDV testing (from rabbits that had died suddenly without obvious cause) are RHDV-negative. Interestingly, a similar finding was reported in pet rabbits in the United Kingdom. This raises the intriguing question of what else is killing rabbits, both in Australia and internationally? Using a metatranscriptomic sequencing approach, we found that *Clostridiaceae, Pasteurella multocida*, and *Eimeria* are frequently detected in cases of sudden rabbit death in Australia. While we did not identify any potential new pathogens that could be explored in the context of wild rabbit management, we have validated an approach to explore future mortality events of lagomorphs that may identify candidate novel biocontrols. Furthermore, our findings reaffirm the recommendation to follow good hygiene practices when handling rabbits, since domestic rabbits harboured several pathogens of potential public health significance, including *Escherichia, Pasteurella multocida*, and Hepatitis E virus.

## Introduction

Australia has a long history of managing overabundant wild European rabbit (*Oryctolagus cuniculus*) populations with biocontrol agents, including rabbit haemorrhagic disease virus (RHDV) and myxoma virus (MYXV) (1). Both viruses have a high case fatality rate and are transmitted mechanically by insect vectors, and in the case of RHDV also through direct contact and fomites (1). RHDV typically presents as sudden death and is only definitively diagnosed through histopathology or molecular testing, while MYXV can frequently be diagnosed based on characteristic clinical signs. However, a recent study reported a novel MYXV disease phenotype (in domestic rabbits with no genetic resistance to MYXV) caused by highly virulent field strains, which also presented as peracute death (2). It is frequently assumed that most cases of sudden death in wild and domestic rabbits in Australia, particularly where multiple deaths occur within a short period, are due to these viruses. Notably, we have observed that approximately 40% of rabbit liver samples submitted for RHDV testing, i.e., from rabbits that had died suddenly without obvious cause, are RHDV-negative (3). Interestingly, a similar finding was reported in pet rabbits in the United Kingdom (UK) (4). This raises the intriguing question of what else is killing rabbits, both in Australia and internationally?

This question is important for several reasons. Firstly, in their native home range on the Iberian Peninsula, rabbits are a keystone species (5). Since the emergence of RHDV2 in 2010 (6, 7) wild rabbit populations in Spain have continued to decline, leading to their reclassification as “endangered” by the International Union for Conservation of Nature in 2018 (8). Secondly, rabbits are a popular pet species, especially for young children. This close human-animal interface poses a potential public health risk for the transmission of zoonotic diseases from rabbits to their owners, particularly when good hygiene practices are not followed. Rabbits are known reservoir hosts of several zoonotic pathogens, including enterohaemorrhagic *Escherichia coli*, *Cryptosporidium*, *Pasteurella multocida*, *Encephalitozoon cuniculi*, and *Hepatitis E virus* (HEV) (9–13). Thirdly, with the increasing accessibility of exploratory sequencing methods (i.e., ‘metagenomics’ and ‘metatranscriptomics’), laboratories can now apply these methods to specific disease syndromes and/or mortality events to detect putative associations with known or emerging pathogens (14–16). Finally, in the Australian context, these pathogen discovery approaches may reveal candidate future biocontrol agents, or potential ecological interactions between microbes (either synergistic or antagonistic), that may enhance future rabbit management approaches.

There are several known causes of sudden death in rabbits. Non-infectious differential diagnoses include degenerative (heart disease, renal disease), developmental (congenital defects), inflammatory (e.g., pancreatitis), neoplastic, nutritional, traumatic, toxic, physical (e.g., liver lobe torsion, intussusception, aspiration pneumonia, heat stroke), and vascular (pulmonary embolism, haemorrhagic syndromes) pathologies (4, 17). Examples of known infectious causes of acute fatalities in rabbits include pasteurellosis, staphylococcosis, hepatic coccidiosis, enterotoxaemia/epizootic rabbit enteropathy (ERE), colibacillosis, Tyzzer’s disease, pseudotuberculosis, tularaemia, myxomatosis, and rabbit haemorrhagic disease (4, 11–13, 18). But are there potentially overlooked pathogens? In this study, we profiled the metatranscriptome of liver samples collected from RHDV-negative rabbits found dead in Australia to determine what putative pathogens may be killing these rabbits.

## Results

### Clostridiaceae, Pasteurella, and Eimeria are common colonists of rabbits in Australia

To identify microbes associated with sudden death of rabbits in Australia, we conducted metatranscriptomic sequencing on 60 RHDV-negative rabbit liver samples collected from Victoria (VIC; n = 38), Tasmania (TAS; n = 8), New South Wales/Australian Capital Territory (NSW/ACT; n = 11), South Australia (SA; n = 2), and Western Australia (WA; n = 1) between 2016 and 2020 (Supplementary table 1). Samples were obtained from a mix of breeds of domestic pet, show, and meat rabbits that ranged from 4.5 weeks to 9 years of age, as well as from two wild rabbits; 23 samples were from does and 33 samples were from bucks (for the remaining 4 samples the sex was not specified). Rabbits were reported to have a wide range of clinical signs prior to death, although many were simply found dead (Supplementary table 1). Notably, six rabbits from Victoria that died between 2017 and 2018, including three from a single shelter facility, were reported with frank haemabdomen. On further investigation, these six rabbits had no access to anticoagulants, there were no clear dietary associations between the cases, and at least four were housed indoors. This prompted us to look more closely at cases from Victorian rabbits, and haemorrhagic signs prior to death were also reported in a further five cases.

The 60 sequencing libraries ranged in size from 6,356,968 to 24,147,560 paired-end reads, of which 8.0–93.6% 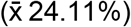 did not map to the phylum *Chordata* (i.e., the vertebrate host). Reads were assembled into contigs, which were used for taxonomic assignment. The transcripts per million (TPM) method was used to normalise the data and to calculate the relative abundance of taxa, where reads were used in place of transcripts. Taxonomic assignment at the kingdom level revealed three clear groupings of samples—bacteria-dominant (n = 9), eukaryota-dominant (n = 1), and unassigned-dominant (n = 50) (Figure 1A). For comparison, metatranscriptomic sequencing of 24 known RHDV+HEV-positive liver samples almost always showed an extremely high proportion of viral reads (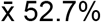; Figure 1B) (19). Overwhelmingly, most unknown samples grouped as unassigned (i.e., most reads were classified as unassigned).

**Figure 1:**
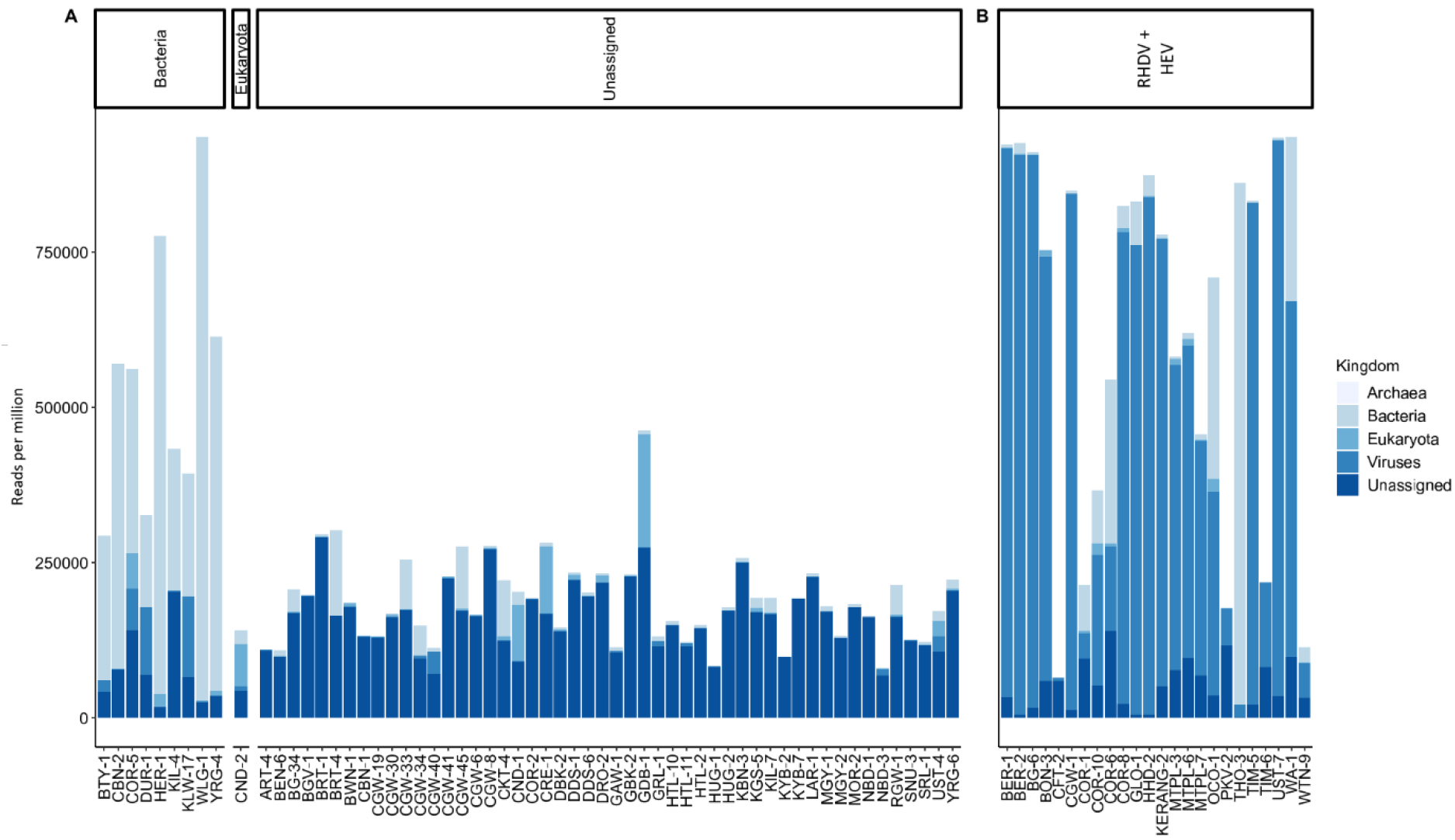
Classification of contigs by kingdom reveals distinct clustering of samples. Metatranscriptomic sequencing was conducted on liver samples from rabbits that had died suddenly and that were negative for *Rabbit haemorrhagic disease virus* (RHDV) (A). Reads were assembled into contigs, classified to the kingdom level, and normalised using the transcript per million method by mapping individual reads to contigs. This revealed three distinct clusters of samples: those with a high proportion of 1) bacterial reads, 2) eukaryotic reads (excluding phylum *Chordata*), or 3) unassigned reads. For comparison, the same analyses were performed on 24 known RHDV+HEV-positive liver samples.

The dominant microorganisms detected included *Hepatitis E virus* (n = 10), phylum *Firmicutes* (particularly family *Clostridiaceae*; n = 15), *Pasteurella* species (phylum *Proteobacteria;* n = 7), *Eimeria* species (n = 4), *Cyniclomyces* yeast (n = 2), and *Pseudomonas* species (phylum *Proteobacteria;* n = 2) (Figure 2 A,C,E,G). Mixed infections were common. Indeed, despite these being liver samples, there were no ‘pure’ infections, where only a single microbe was identified. Even in known RHDV+HEV-positive samples, *Eimeria, Cyniclomyces* yeast, *Firmicutes*, *Proteobacteria*, and various other bacterial genera were frequently detected (Figure 2 B,D,F,H). Importantly, sample collection occurred at variable times after death and was not performed in sterile conditions, so environmental contamination and translocation from the gastrointestinal tract is highly likely.

**Figure 2:**
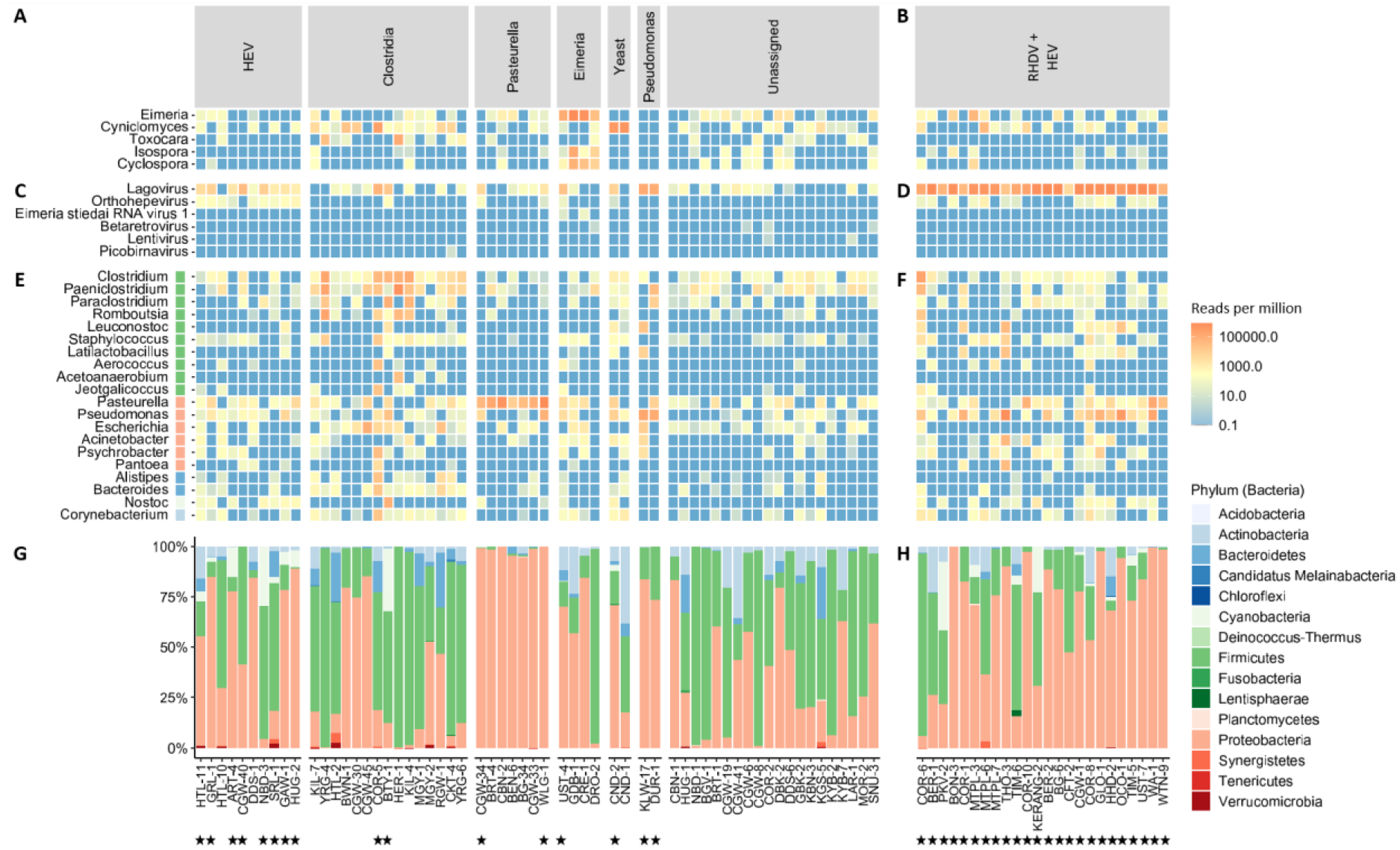
Classification of contigs from metatranscriptomic sequencing of rabbit liver samples to the genus (A-F) and phylum (G-H) level. Samples (along the x-axis) were grouped based on their most abundant microbial reads as: *Hepatitis E virus* (HEV), *Clostridia, Pasteurella, Eimeria*, yeast, *Pseudomonas*, unassigned, or *Rabbit haemorrhagic disease virus* (RHDV). Heatmaps based on reads per million were generated for eukaryotic reads (A,B), viral reads (C,D), and bacterial reads (E,F), with individual genera listed on each line (y-axis). Boxes adjacent to the bacterial genera (E,F) are coloured based on their respective phylum classification. Stacked bar plots for each sample show the proportion of bacterial reads by phylum (G,H). Stars indicate samples that were included in sequencing run 2, while samples without stars were sequenced on run 1.

In addition to *Eimeria* and *Cyniclomyces* yeast, other eukaryotic reads detected included those of the roundworm *Toxocara* and the coccidian parasites *Isospora* and *Cyclospora* (Figure 2A). The latter reads correlated strongly with the presence and abundance of *Eimeria* reads, suggesting that perhaps conserved coccidia regions were misclassified between these three genera. RHDV reads were detected in most samples (Figure 2C), however, this most likely reflects cross-contamination of the flow cell during sequencing, since RHDV-positive and -negative samples were combined in the same sequencing run and the abundance of RHDV reads in positive samples is extremely high (Figure 1B, Figure 2D). Indeed, samples from run 2, which comprised 24 RHDV-positive samples, revealed a higher level of RHDV reads than samples from run 1, which included two RHDV positive samples (that were not part of this study). Other viruses identified included HEV, *Eimeria stiedai RNA virus 1* (in two samples, both of which were also positive for *Eimeria*), retroviruses (likely reflecting rabbit endogenous retroviruses), and a rabbit picobirnavirus in one sample (Figure 2C). As well as the dominant bacterial genera discussed above, other putative bacterial pathogens were detected sporadically, such as *Escherichia*, *Staphylococcus*, *Corynebacterium*, and *Bacteroides*, but typically at low abundance and/or secondary to other dominant microbes. Furthermore, many likely commensal and/or environmental bacterial genera were identified frequently and typically at low abundances.

For multiple samples collected from the same ‘outbreak’ event, there was not always a strong correlation between the dominant microorganism detected (Supplementary table 1). For example, *C. cuniculi* was detected in CBN-2 but CBN-1 was classified as unassigned. Similarly, *Clostridiacae* were detected in COR-5 but not in COR-2. However, for samples CND-1 and −2, *Cyniclomyces* yeast were detected in both cases. *C. cuniculi* and HEV were detected in both HTL-10 and HTL-11. *Clostridiacae* were detected in both MGY-1 and MGY-2, although *C. cuniculi* specifically was only detected in MGY-2. While no infectious agents were identified for KYB-2 and KYB-7, both samples were classified as unassigned. For Victorian samples with a haemorrhagic presentation, most samples were classified as unassigned, with *Clostridiacae* and HEV each being identified in two of 11 cases.

To verify detections of HEV, *Clostridiacae*, *Pasteurella*, and *Eimeria*, and to confirm the RHDV status of the samples, we conducted confirmatory RT-PCR and RT-qPCR testing (Figure 3). The ‘Clostridium generic’ RT-PCR showed poor sensitivity for the presence of the rabbit-specific *C. cuniculi*, so all samples were tested with both the ‘Clostridium generic’ and *‘Clostridium cuniculi’* RT-PCRs. Additionally, to identify clostridial pathogens to the species level, several specific RT-PCRs for *Paeniclostridium sordellii* (previously *C. sordellii*), *C. perfringens*, *C. spiroforme*, and *C. piliforme* were tested on samples that were positive on the ‘Clostridium generic’ RT-PCR. *C. cuniculi* was the most common clostridial species identified (n = 19), followed by *P. sordellii* (n = 8), and two detections each of *C. perfringens* and *C. spiroforme. C. piliforme* was not confirmed in any samples. Mixed clostridial infections were common (n = 8) (Figure 3).

**Figure 3:**
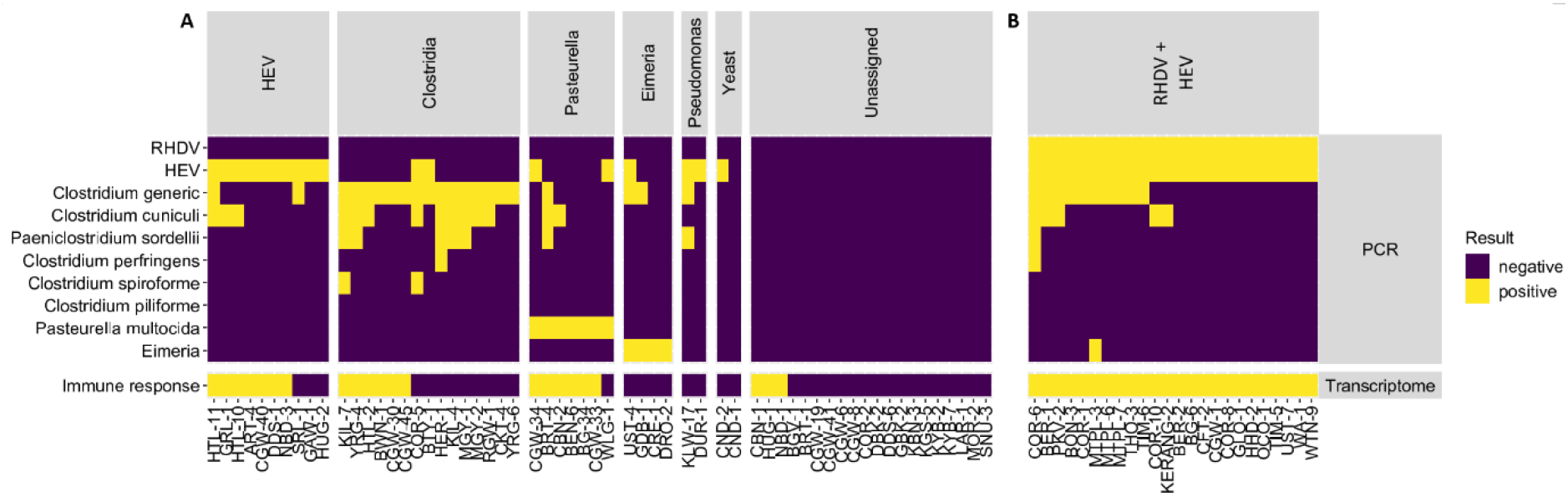
Confirmatory RT-PCR and RT-qPCR testing and host transcriptome analyses. RHDV-negative liver RNAs were screened by specific RT-PCRs or RT-qPCRs targeting RHDV, HEV, a conserved region of the *Clostridiacae* class (‘Clostridium generic’), *C. cuniculi, Pasteurella multocida*, and rabbit *Eimeria* species (A). *Clostridiaceae*-positive samples were further screened for four additional clostridial species: *P. sordellii*, *C. perfringens*, *C. spiroforme*, and *C. piliforme*. Samples are faceted by the dominant microorganism detected through metatranscriptomic sequencing. Yellow squares indicate a positive result while purple squares indicate that the target was not detected. Samples were classified for ‘immune response’ based on host transcriptome analysis. Samples were considered positive if a gene ontology term related to the immune response or defense mechanisms (Supplementary table 2) yielded a positive enrichment score relative to known healthy controls. For comparison, the same analyses were performed on 24 known RHDV+HEV-positive liver samples (B).

We further explored epidemiological factors associated with *Clostridiacae, P. multocida*, and *Eimeria* infections in Australian rabbits. Due to the sampling strategy employed, there was a strong bias towards areas with larger domestic rabbit populations, namely VIC, TAS, and NSW/ACT (Figure 4A). Interestingly, all four samples where *Eimeria* was detected had a history of multiple contemporaneous deaths and all were from young animals (Figure 4B, Supplementary table 1). In contrast, *Pasteurella* detections were mostly in adult animals, although importantly the sample size was small (Figure 4C, Supplementary table 1). One detection was in a wild rabbit. Clostridial infections showed no clear association with age or geography (Figure 4D). Strikingly, the relative abundance of *Pasteurella* reads was extraordinarily high in positive samples (up to almost 800,000 reads per million), compared to more moderate relative abundances (up to ~200,000 reads per million) observed for *Clostridiacae* and *Eimeria* (Figure 4).

**Figure 4:**
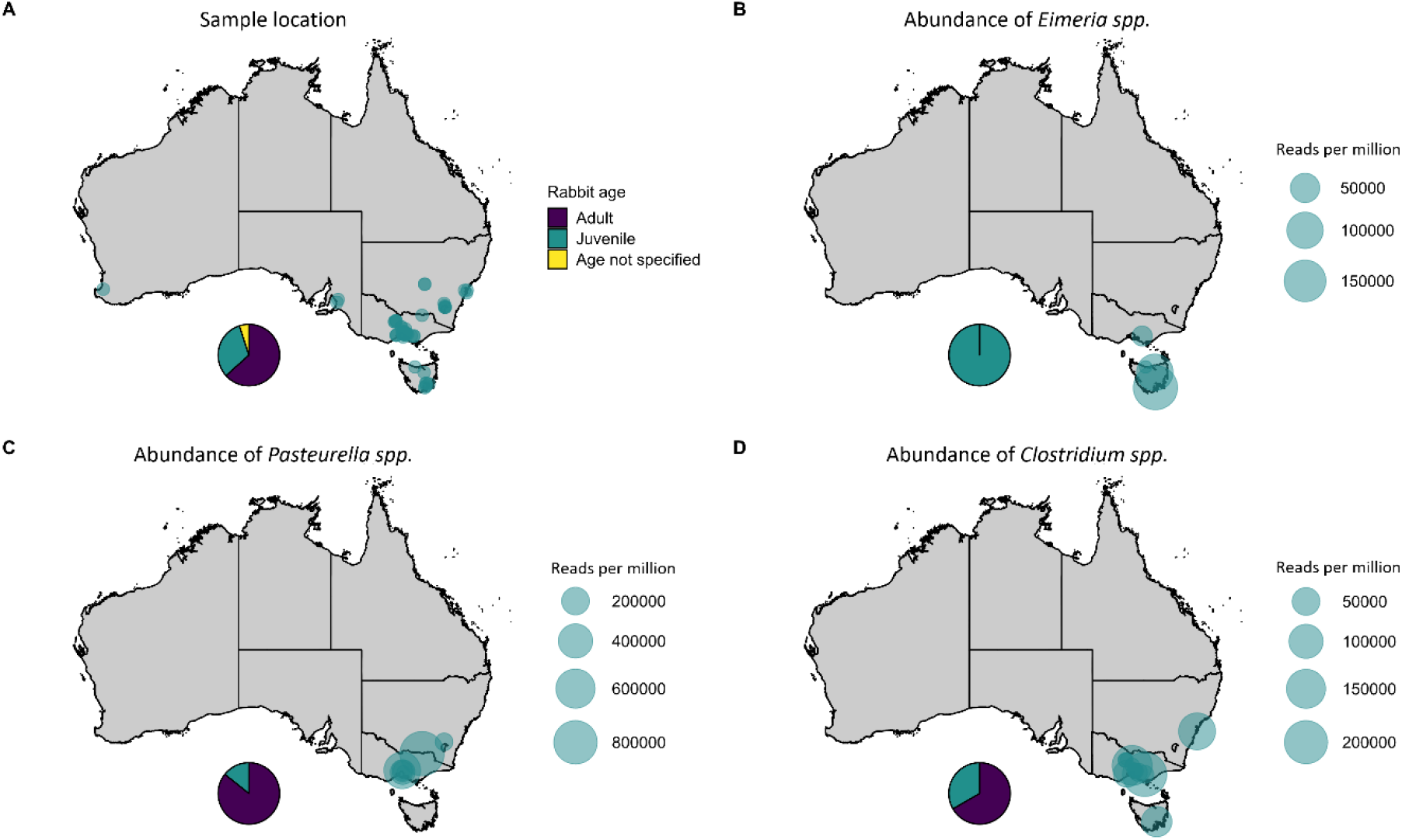
Epidemiological factors associated with *Clostridiacae, Pasteurella*, and *Eimeria* infections in Australian rabbits. Metatranscriptomic sequencing was conducted on liver samples from rabbits that had died suddenly and that were negative for *Rabbit haemorrhagic disease virus* (RHDV). Maps show the sampling location (A). Pie charts represent the distribution of samples by age (juvenile vs adult). The relative abundance (reads per million) of *Eimeria* (B), *Pasteurella* (C), and *Clostridiaceae* (D) in positive samples is represented by the size of each circle. WA – Western Australia, NT – Northern Territory, SA – South Australia, QLD – Queensland, NSW – New South Wales, ACT – Australian Capital Territory, VIC – Victoria, TAS – Tasmania.

### Host transcriptome

While metatranscriptomic analyses can identify the presence of microbial reads and high abundance may be suggestive of fulminant infection, these analyses cannot reliably be used to infer pathogenicity/cause of death at an individual level. Therefore, we interrogated the ‘residual’ host transcriptome for gene ontology (GO) terms in the ‘biological process’ domain for processes related to immune responses and/or defense mechanisms (hereafter described as ‘immune response’). Host transcriptome data from three healthy laboratory rabbits generated in a previous study were used as ‘known non-infectious cause of death’ controls (20). GO terms that we considered as indicative of an immune response, based on a positive enrichment score relative to healthy control laboratory rabbits, are detailed in Supplementary table 2. The 24 known RHDV+HEV-positive liver samples (described above) served as ‘known infectious cause of death’ samples.

As expected, all RHDV+HEV-positive samples showed evidence of an upregulated immune response in the host transcriptional profile relative to the healthy controls, and principal component analysis (PCA) showed clear segregation of these two groups (Supplementary figure 1). Immune responses were detected in most (6/7) *Pasteurella*-positive samples, most (7/10) HEV-positive samples, some (6/15) *Clostridiaceae*-positive samples, but only 3 of 20 samples classified as unassigned by metatranscriptomic analysis (Figure 3, Supplementary figure 2). No positive enrichment of immune responses was detected in *Eimeria*-positive samples, *Pseudomonas*-positive samples, or samples where *Cyniclomyces* yeast was detected. There was considerable overlap in the host transcriptome PCA between unknown cause of death samples (Supplementary figure 1), indicating poor resolution in host responses to different pathogens at the global transcriptome level.

## Discussion

Through our long-term Australian lagovirus surveillance program, we were surprised to observe that approximately 40% of rabbit liver samples collected from rabbits that had died suddenly were negative for RHDV (3), with a recent UK study presenting similar findings (4). To identify whether other infectious agents may be responsible for these sudden deaths, particularly in outbreak situations where multiple rabbit deaths were reported, we undertook metatranscriptomic sequencing of 60 RHDV-negative rabbit liver samples. While reads from several putative bacterial and eukaryotic pathogens were identified at high relative abundance, including several *Clostridiaceae* species, *Pasteurella*, *Eimeria*, and *Pseudomonas*, most liver samples were classified as ‘unassigned’, where no hit could be identified in the NCBI nucleotide database. This suggests that most cases of sudden death in (RHDV-negative) rabbits may be due to non-infectious causes. Alternatively, liver samples may not have been suitable for diagnosis of these cases, or the pathogen may not have been present at high abundance at the time of sampling. Interestingly, three of these ‘unassigned’ cases showed positive enrichment for immune responses on host transcriptome analysis, raising the possibility of a systemic inflammatory response secondary to either non-infectious pathology or a previous infection that was no longer detectable. Indeed, one of these cases (HUG-1) did report pyrexia in the clinical history.

Only liver samples were available because samples were submitted originally for diagnosis of hepatotropic lagovirus infections. Because of the sampling strategy employed, samples were heavily biased towards domestic rabbits, with most samples submitted from VIC, TAS, and NSW/ACT. Samples were collected at variable times post-mortem, without regard for sterility, with variably complete clinical histories, and sample transport and storage were not ideal for metatranscriptomic analyses (i.e., samples were not snap-frozen at −80 °C), which may have adversely impacted our findings. Importantly, the RHDV+HEV-positive samples were derived from the same sampling program, making them ideal controls since they were subject to the same limitations, and upregulation of immune responses were still detectable in these samples. For metatranscriptomic analyses to be revealing, the pathogen must be transcriptionally active in the sampled tissue at the time of sampling. Since the liver is generally considered to be a sterile site, this organ is infrequently targeted for exploratory metatranscriptomic analyses (21). However, because of the highly vascular nature of the liver, it could be expected that any systemic infection would also be detected in this tissue. Indeed, it may be easier to differentiate pathogens from the healthy commensal microbiome using liver samples compared to, for example, gastrointestinal tract samples. While metatranscriptomic analyses can only provide evidence of association, not causation, several criteria support an agent being potentially pathogenic. For example, in the context of this study, finding an agent at high abundance, consistently across several similar cases, temporally associated with sudden death, particularly if known to be pathogenic in other species, and with corresponding transcriptomic evidence of upregulation of immune responses, would additionally support a putatively causal relationship.

The incorporation of host transcriptome analysis into a metatranscriptomic survey offers a novel and innovative approach to the diagnosis of infectious disease via metatranscriptomics, utilising the host mRNA data that is usually discarded in such analyses. While subject to several limitations in the implementation of this study, such as variable post-mortem degradation of samples, we clearly observed positive enrichment of immune responses and defense pathways in our known RHDV+HEV-positive controls, which had been subjected to similarly variable sampling and handling regimes. Notably, most of the unassigned cases generally did not show evidence of upregulation of host immune responses. In summary, whilst the detection of upregulated immune genes is still not proof for causation (such as in the case of HEV), inclusion of these data in combination with other findings such as high abundance of a single dominant microorganism can lend additional support to the hypothesis of infection as a contributing factor to death.

While HEV was identified in 18 samples, we do not suspect this to be the primary cause of death in these cases, despite many cases also showing evidence of an immune response in their transcriptome profiles. HEV is present globally in wild and domestic rabbit populations at relatively high seroprevalence (3–60%) (22), yet was only identified for the first time in rabbits in 2009 through a serosurvey of farmed rabbits (23). This suggests that it is not a major cause of morbidity or mortality, at least in healthy animals. Experimental infection studies have shown that while rabbits can develop both acute and chronic hepatitis following HEV infection, infection is often subclinical and sudden death has not been observed, except in pregnant rabbits (24–27). A recent study found a seroprevalence of 9% in healthy shot wild rabbits in Australia (19), providing further support that HEV was likely an incidental finding.

We identified several clostridial species in rabbit liver samples in this study, including *C. cuniculi*, *Paeniclostridium sordellii, C. spiroforme*, and *C. perfringens*, first through metatranscriptomic sequencing and subsequently verified by RT-PCR. Toxigenic *Clostridium* species, particularly *C. spiroforme* but also infrequently *C. perfringens* and *C. difficile*, are known to cause enterotoxaemia in rabbits, a major cause of acute diarrhoea leading to severe dehydration and death in 24–48 hours (12). A similar syndrome, epizootic rabbit enteropathy (ERE), has recently been associated with *C. cuniculi* overgrowth (28, 29). Both syndromes frequently occur in farmed rabbits at weaning with very high (30–95%) mortality (12, 28). The disease is multifactorial, with stress, dietary changes, or antibiotic use triggering gastrointestinal dysbiosis leading to subsequent proliferation of *Clostridium* species, sometimes with secondary opportunistic overgrowth of coliforms (12). Coinfections with other pathogens (such as enteropathogenic *E. coli*, *C. piliforme*, rotaviruses, and *Eimeria*) are common, with one study identifying coinfections in 86% of rabbits with enterotoxaemia (29, 30). Neither *C. spiroforme* nor *C. cuniculi* are typically observed in the microbiome of healthy rabbits (12, 28). Given the major disruption to the gut epithelium in both enterotoxaemia and ERE and the high abundance of *Clostridium* species during fulminant disease, it would not be surprising to observe bacterial translocation into the bloodstream with subsequent detection in the liver. However, there was evidence of a host immune response in only 40% of samples in this study, although poor sample quality could have adversely affected this analysis. While *P. sordelii* is not classically associated with enterotoxaemia in rabbits, it has been associated with various enteric and histotoxic infections in a wide variety of species (31). However, its role in disease is controversial, as it is a common environmental bacterium found in soil (31). Another important clostridial pathogen of rabbits is *C. piliforme*, the causative agent of Tyzzer’s disease, characterised by diarrhoea, dehydration, multifocal hepatic necrosis, and death in 1–2 days (12). While *C. piliforme* contigs were identified here, the relative abundance was low compared to other clostridial species and detections could not be verified by PCR (32). It is possible that the contigs mapping to *C. piliforme* spanned conserved clostridial genomic regions and were misclassified from other species. The lack of specific RT-PCR detections suggests that none of these rabbits succumbed to Tyzzer’s disease.

*Pasteurella multocida* is considered to be the most common bacterial pathogen of laboratory rabbits (12) and indeed, we identified *P. multocida* in 7 of 60 samples. There are multiple clinical manifestations of pasteurellosis, including rhinitis, pneumonia, genital tract infections, otitis media, and septicaemia. *P. multocida* is also a common commensal in the rabbit nasopharynx; for example, one study showed that 31% of healthy rabbits were infected asymptomatically (33). Septicaemia typically occurs from haematogenous spread following localised disease and is rapidly fatal. In these cases, *P. multocida* can be recovered from parenchymal organs (12). Therefore, it is highly probable that our detections of this organism were clinically significant, especially given the very high abundances observed in *Pasteurella*-positive samples and the corresponding transcriptional upregulation of host immune responses in 6 of the 7 positive samples. Other *Pasteurella* species known to infect rabbits include *P. pneumotropica* and *P. aerogenes*, although neither have been associated with systemic disease (12).

*Eimeria* are apicomplexan parasites that cause coccidiosis. Eleven *Eimeria* species infect rabbits, resulting in hepatic coccidiosis (*E. stiedae*) or intestinal coccidiosis (the remaining 10 species) (13). All rabbit *Eimeria* species can be carried subclinically, typically by adult animals, which serve as the infection source for young animals. Disease is enhanced by stressors such as overcrowding, poor hygiene, poor nutrition, transportation, and weaning. Hepatic coccidiosis is characterised by severe liver disease, resulting in anorexia, ascites, icterus, and death, particularly in young animals 2–3 months of age (13). Intestinal coccidiosis manifests as diarrhoea, the severity of which depends on the pathogenicity of the infecting species. The presence of *E. stiedae* was confirmed via RT-PCR with *E. stiedae* specific primers (34, 35). Of note was the detection of *Eimeria stiedae virus RNA 1* in two *Eimeria*-positive samples, further confirming the presence of *Eimeria stiedae* in those rabbits. This is a double-stranded RNA virus belonging to the family *Totiviridae* (36) known to specifically infect *E. stiedae* (37). All *Eimeria* infections identified here were in young animals and multiple deaths were reported in each case; diarrhoea was not a feature of these cases. None of the positive samples showed a positive enrichment score for host immune responses on transcriptome analysis, which is perhaps not unexpected given that death is due to secondary liver failure. While *Cyclospora* and *Isospora* were also identified in our samples, both of which are also coccidian parasites, these species are not known to infect rabbits and the presence and abundance correlated strongly with detections of *Eimeria*. Therefore, we suggest that these were probably *Eimeria* contigs spanning conserved genomic regions that were misclassified as *Cyclospora* or *Isospora*.

Reads matching *Cyniclomyces* yeast were detected at high relative abundance in two samples, although the clinical significance of this finding remains unclear. The most well-characterised species of this genus is *C. guttulatus* (formerly *Saccharomycopsis guttulata*), a normal commensal of the gastrointestinal tract of rabbits and rodents (38). Although it has been detected in association with various clinical presentations, such as oculonasal discharge and systemic abscesses, bloat, enteritis, and coccidiosis, most researchers agree that this is likely to be an opportunistic pathogen or a secondary overgrowth following a prior insult (38). We did not find an association with the presence of *Eimeria* in this study. Following experimental infections with *C. guttulatus*, healthy rabbits remain asymptomatic (38, 39). The detection of this yeast in liver samples may suggest translocation post-mortem or contamination during sample collection. The lack of enrichment for immune responses on host transcriptome analysis provides further support that these yeasts were not primary pathogens in these samples.

*Pseudomonas* reads were identified at high abundance in two rabbit liver samples. *P. aeruginosa* is a common environmental bacterium and is well-known to cause opportunistic, often severe, infections in a range of species, including humans. In rabbits, infections are typically associated with dermatitis but there are also reports of abscessation, septicaemia, pneumonia, and diarrhoea (12). Interestingly, we found that *Pseudomonas* reads were also abundant in many RHDV+HEV-positive samples, although confirmatory RT-PCR analyses were negative. The widespread distribution of this organism in the environment suggests that these detections were likely environmental contaminants rather than co-infections, particularly since there was no evidence of transcriptional upregulation of host immune responses or defense pathways in *Pseudomonas*-positive samples.

Other potentially interesting findings in this study included *Streptococcus pneumoniae* in CRE-1 (*Eimeria*-positive), *Streptococcus sanguinis* in YRG-6 (*Clostridiaceae*-positive), a picobirnavirus in CKT-4 (*Clostridiaceae*-positive) and *Toxocara*, retroviruses, *Staphylococcus*, *Escherichia*, and *Corynebacterium* in several samples with varying relative abundance. Infections with *Streptococcus* species have infrequently been reported in rabbits, but have rarely been associated with septicaemia, abscesses, and osteomyelitis (12). Genetically divergent picobirnaviruses have been identified previously from rabbit faeces and caecal contents but are now thought to represent bacterium-associated viruses rather than vertebrate pathogens (11, 21). Rabbits can act as aberrant hosts for *Toxocara canis*, the dog roundworm, and other ascarids. Migration of parasite larvae through the tissues (visceral larva migrans) after the eggs hatch in the intestine can cause various clinical signs, including neurological signs, but would not be expected to cause sudden death (13). No pathogenic retroviruses have been reported in rabbits and the betaretrovirus and lentivirus contigs detected here likely derive from endogenous retroviruses (40). Staphylococcosis caused by *S. aureus* is a common infection of rabbits, with clinical signs similar to those observed with pasteurellosis and occurring either sporadically in individual rabbits or as an epizootic (12). *Staphylococci* species are common commensals of skin and mucous membranes and different species and strains vary in virulence, making definitive association with disease challenging in the context of metatranscriptomics. Colibacillosis is a diarrhoeal disease of either neonates or weanling rabbits, sometimes with high mortality. *Escherichia coli* is a normal component of the gastrointestinal flora and, while it can be a primary pathogen, it also proliferates in cases of enteritis caused by other pathogens (12). Indeed, in this study *Escherichia* was observed to be present at high abundance in many samples primarily classified as *Clostridiaceae-positive. Corynebacterium bovis* has been associated with systemic disease and testicular abscessation in rabbits both clinically and experimentally (12). However, other *Corynebacterium* species are probably also a normal part of the microbiota, as they are in other species (41).

Despite samples being selected because they were RHDV-negative on sensitive and specific RT-qPCR and RT-PCR assays (42), RHDV was identified in most samples, albeit at lower abundance than seen in known RHDV+HEV-positive samples. This most likely reflects cross-contamination of the flowcell during sequencing, since RHDV-positive and -negative samples were combined in the same sequencing run and the abundance of RHDV reads in positive samples is extremely high. The average viral RNA load in the liver of infected animals is 3 × 10^8^ capsid copies per mg of tissue, which equates to 1.2 × 10^8^ capsid copies per μl of RNA (42). Indeed, the highest relative abundances of RHDV was observed in samples from sequencing run 2, which also included 24 RHDV+HEV positive samples. Both inter-run and intra-run contamination are known concerns with Illumina platforms. For example, several studies have reported that up to 10% of reads from a sample can be incorrectly assigned when multiplexing, particularly with ExAmp chemistry such as that used for the NovaSeq (43–45). For this reason, a non-redundant dual-indexing strategy would have been preferable in hindsight. However, we also cannot rule out low-level cross-contamination during sequencing library preparation or RNA extraction since extraction controls were not sequenced.

Finally, while our sample size was relatively small, several notable pathogens were not identified in this study. For example, *Salmonella enterica*, while uncommon in rabbits, can cause epizootics with high morbidity and mortality and can potentially be transmitted to humans (12). *Listeria monocytogenes* is also an infrequent cause of sudden death in rabbits but is significant from a public health perspective. *Francisella tularensis*, the causative agent of the zoonotic disease tularaemia, is endemic in wild rabbits and hares in Eurasia and North America and can cause sudden death in these species (12). Recently, four locally acquired human cases of tularaemia have been reported in Australia, linked to contact with infected possums, however an animal reservoir of *F. tularensis* has not yet been identified locally (46, 47). While our study focussed mainly on domestic rabbits, we did not detect any *Francisella* contigs in these samples. Surprisingly, we also did not identify MYXV in this study. Recently, a highly lethal immune collapse syndrome was demonstrated in domestic rabbits infected with MYXV isolates from the 1990s (2). Given the active circulation of MYXV in wild rabbit populations in Australia, we had expected to find MYXV as a cause of death in RHDV-negative domestic rabbits. *Leporid herpesvirus 4* is a recently emerged alphaherpesvirus that was isolated from a mass mortality event in Alaska in 2008 and from a single pet rabbit in Canada in 2010 (48, 49). It has not been reported elsewhere and was also not detected in the rabbits analysed in our study. Other viruses known to be associated with sudden death in rabbits include rabbit enteric coronavirus. Finally, no fungal contigs were identified in these samples, although rabbits appear to be remarkably resistant to systemic mycoses (17).

In summary, while sudden death in domestic rabbits in Australia can mostly be attributed to RHDV, our study found that *Clostridiaceae, Pasteurella multocida*, and *Eimeria* are also frequently detected in cases of sudden rabbit death. Importantly however, most non-RHDV cases of sudden death in Australian rabbits were not able to be attributed to a known pathogen and no novel putative rabbit pathogens were identified. Furthermore, our findings reaffirm the recommendation to follow good hygiene practices when handling rabbits, since domestic rabbits were found to harbour several pathogens of potential public health significance, including *Escherichia, Pasteurella multocida*, and HEV. While this study did not reveal any potential new pathogens that could be explored in the context of wild rabbit management, we have validated an approach to explore future mortality events of lagomorphs either in Australia or internationally that may identify candidate novel biocontrols. Similarly, we demonstrate that the use of host transcriptome data can lend additional support to diagnosing an infectious cause of death or conversely, suggesting absence of infection.

## Material and Methods

### Sample selection

Samples were selected from a rabbit tissue bank established for lagovirus surveillance (50). No animal ethics approvals are required for sampling rabbits that are found dead in Australia. Samples from NSW and ACT were grouped together, since the ACT is a small (~2400 km^2^) enclave within NSW. Since RHDV is hepatotropic, liver was generally the only sample available. Samples were collected post-mortem (at various times post-death) by pet owners and veterinarians and were stored in an RNA preservative solution at −20 °C. RHDV-negative samples were selected initially (n = 45) based on a detailed clinical history, with a preference for cases where sudden deaths had occurred in multiple rabbits over a short time period (42). Because of these selection criteria, most cases were from domestic rabbits. Subsequently, 34 known HEV-positive domestic rabbit liver samples (from the same sample collection), 24 of which were also RHDV-positive, were sequenced for another study (19) and the data were re-analysed here.

### Metatranscriptomic sequencing

Total RNA was extracted from 20–30 mg of liver tissue with the Maxwell SimplyRNA Tissue Kit (Promega) on a Maxwell RSC 16 instrument (Promega) after homogenisation with glass beads, as described previously (19). Libraries were prepared for metatranscriptomic sequencing using the NEB-Next Ultra II RNA Library Prep Kit for Illumina (New England Biolabs) with the addition of an rRNA depletion step (NEBNext rRNA Depletion Kit (Human/Mouse/Rat), New England Biolabs). Sequencing was performed on an Illumina NovaSeq6000 instrument (SP300 cycle flow cell) at the Biomolecular Resource Facility (BRF), The John Curtin School of Medical Research, Australian National University. Raw reads were deposited in the NCBI Sequence Read Archive under Biosample accession numbers SAMN24852673 - SAMN24852758, BioProject accession number PRJNA796430.

### Data analysis

Raw data were pre-processed using FastQC (v0.11.08), Trimmomatic (v0.38) (51) and FLASh (v1.2.11) (52), as described previously (19). Cleaned reads were mapped against the rabbit reference genome (GCA_000003625.1 OryCun2.0) using Bowtie2 (v2.2.9) (53) to filter out host reads. The remaining reads were assembled into contigs using Trinity (v2.12.0) (54) and contigs were blasted against the NCBI nt database (BLAST+ v2.12.0; default parameters). Results with a query coverage of less than 50% were discarded and TaxonKit (v0.8.0) (55) was used to assign taxonomic lineages to each remaining BLAST hit. All reads used for assembly were then mapped against the assembled contigs to calculate the coverage per contig and the relative abundance of each taxon in TPM (56) (i.e., the proportion of one million randomly selected reads that match the taxon of interest) using SAMtools (v1.12) (57) and R (v4.1.0) (58). Bacterial phyla with an abundance of less than 100 across all samples were excluded for the bar plot. Heats maps were generated using the R package ampvis2 (v2.7.11) (59).

### Host transcriptome analysis

Raw reads were processed as described above. Additionally, previous transcriptomic data from three healthy laboratory rabbits generated in another study (20) were used as ‘known non-infectious cause of death’ controls. Reads were mapped against the rabbit reference genome (GCA_000003625.1 OryCun2.0) using TopHat (v2.1.1) (60). Reads per exon were counted using HTSeq (v0.13.5) (61). Exons were matched to Entrez-IDs and exons without an Entrez-ID were discarded, as no further GO information could be gathered. The DESeq2 package (v1.32.0) (62) in R (v4.1.0) (58) was used to calculate the log2fold changes and *p*-values for all genes compared to the ‘known non-infectious cause of death’ control samples. Genes with an adjusted *p*-value <0.05 were used for a GO Gene Set Enrichment Analysis in the “biological processes” category using the GOSemSim (v2.18.1) (63) and clusterProfiler (v4.0.5) (64). GO-terms with a *p*-value <0.05 were considered significant.

### Confirmatory RT-PCR

Specific RT-PCRs using the SuperScript III One-Step RT-PCR System with Platinum Taq DNA Polymerase (Invitrogen) were run to verify the presence of *Clostridiaceae* species (65), *Clostridium cuniculi* (28), *Paeniclostridium sordellii* (65), *Clostridium perfringens* (65), *Clostridium spiroforme* (66), *Clostridium piliforme* (32), *Pasteurella multocida* (67), and *Eimeria stiedae* (34, 35). Briefly, each 25 μl reaction contained 12.5 μl of reaction mix (2x), 9.5 μl of nuclease-free water, 1 μl of 10 μM primer mix, 1 μl of enzyme mix and 1 μl of total RNA. *Eimeria* and *Pasteurella* PCR reactions were run under the same cycling conditions: 45 °C for 15 min, 94 °C for 2 min, followed by 35 cycles of 94 °C for 15 sec, 55 °C for 30 sec and 68 °C for 90 sec and a final elongation at 68 °C for 120 sec. The *Clostridium* PCR reactions all used 68 °C for 120 sec for elongation, except for *C. spiroforme* and *C. piliforme* where the elongation time was reduced to 30 sec. PCR products were visualized on a 1% agarose gel for a band of appropriate size. The presence of *Hepatitis E virus* and RHDV were verified via RT-qPCR as previously described (19, 42).

## Acknowledgements

We wish to thank all sample submitters, in particular Belinda Oppenheimer for her extensive contributions and helpful advice. Rabbit tissue samples were obtained through a project funded by the Centre for Invasive Species Solutions (P01-B-002). We thank Pat Blackall (The University of Queensland) for providing helpful advice and a positive control for the *P. multocida* PCR. We wish to thank Ina Smith, Alexander Gofton and John Roberts for critical revision of the manuscript.

## Funding

This project was co-funded by Meat and Livestock Australia (P.PSH.1059) and CSIRO. Rabbit tissue samples were obtained through a project funded by the Centre for Invasive Species Solutions (P01-B-002).

## Notes

### Competing Interest Statement

The authors have declared no competing interest.

