## Supplementary figure 2 for "Pathogen profiling of Australian rabbits by metatranscriptomic sequencing"

Supplement figure 2A

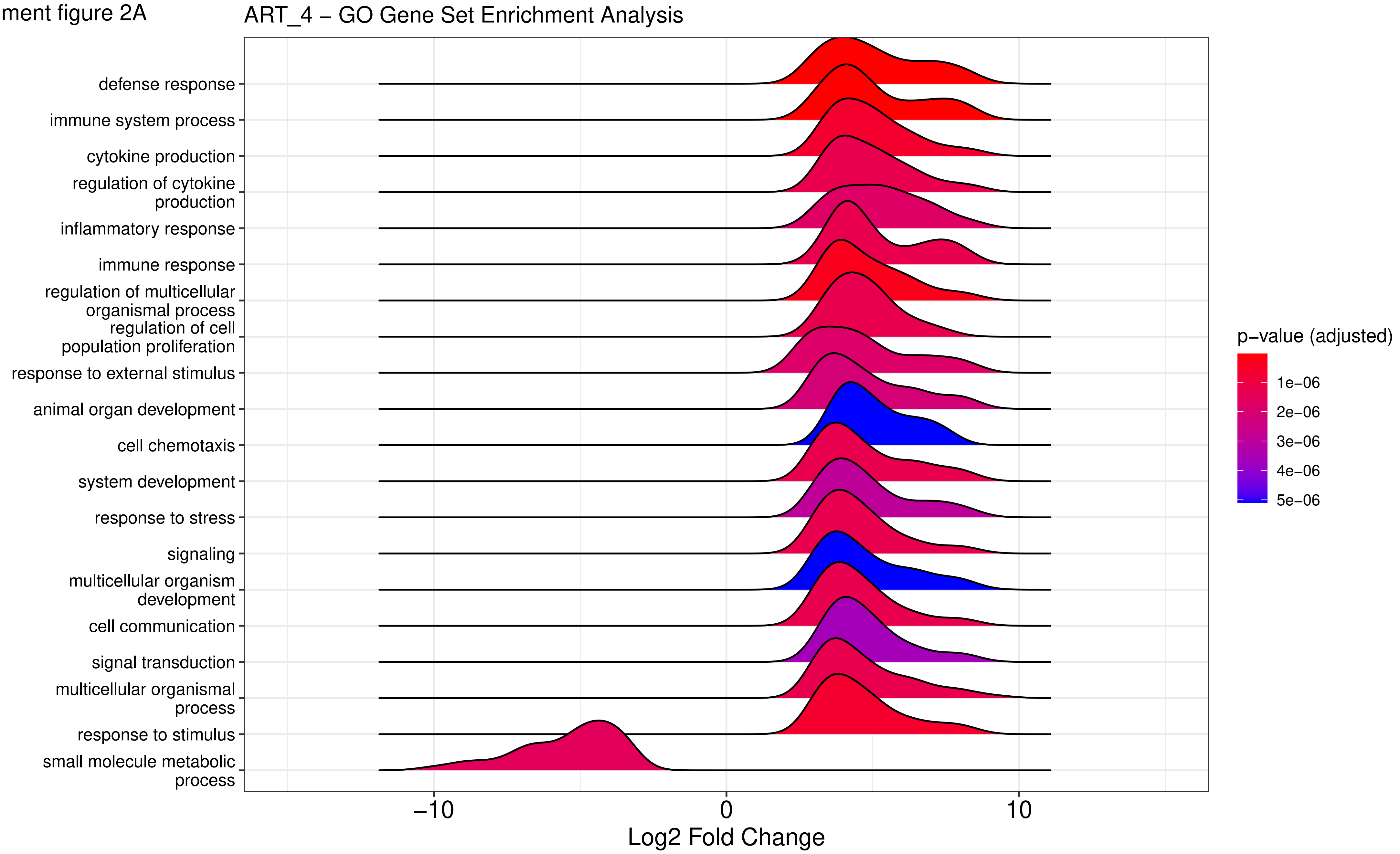

Supplement figure 2B

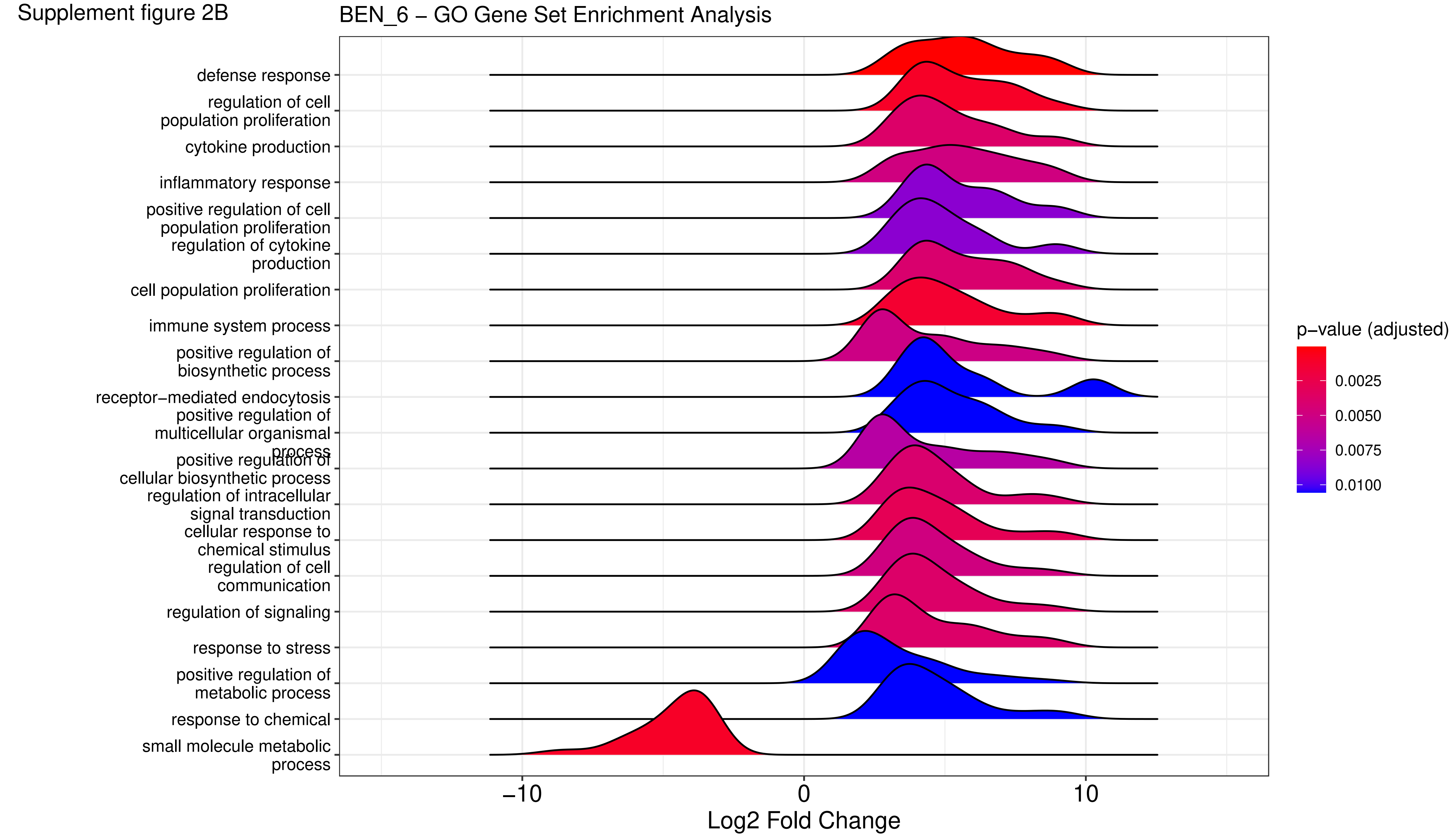

Supplement figure 2C

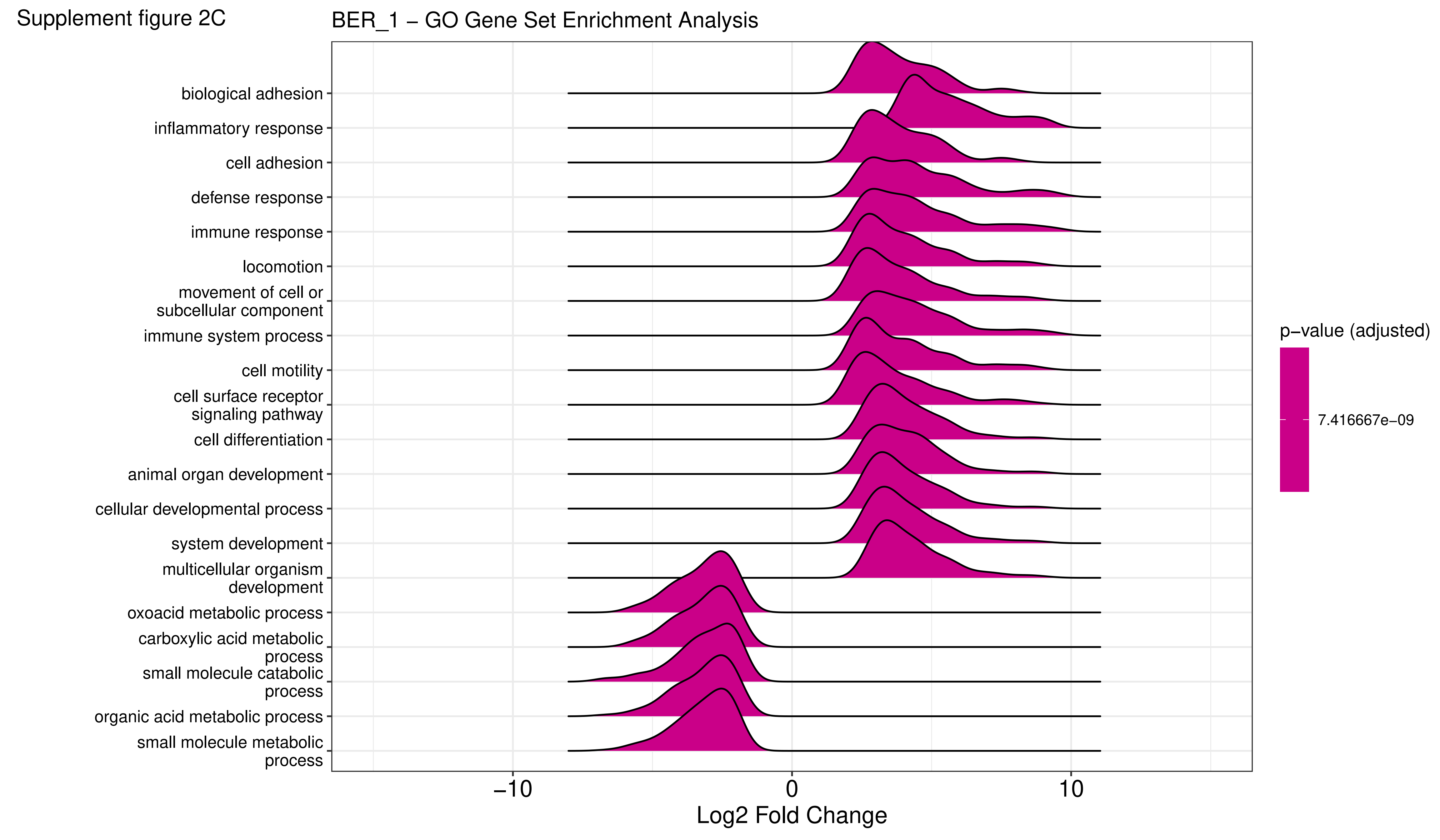

Supplement figure 2D

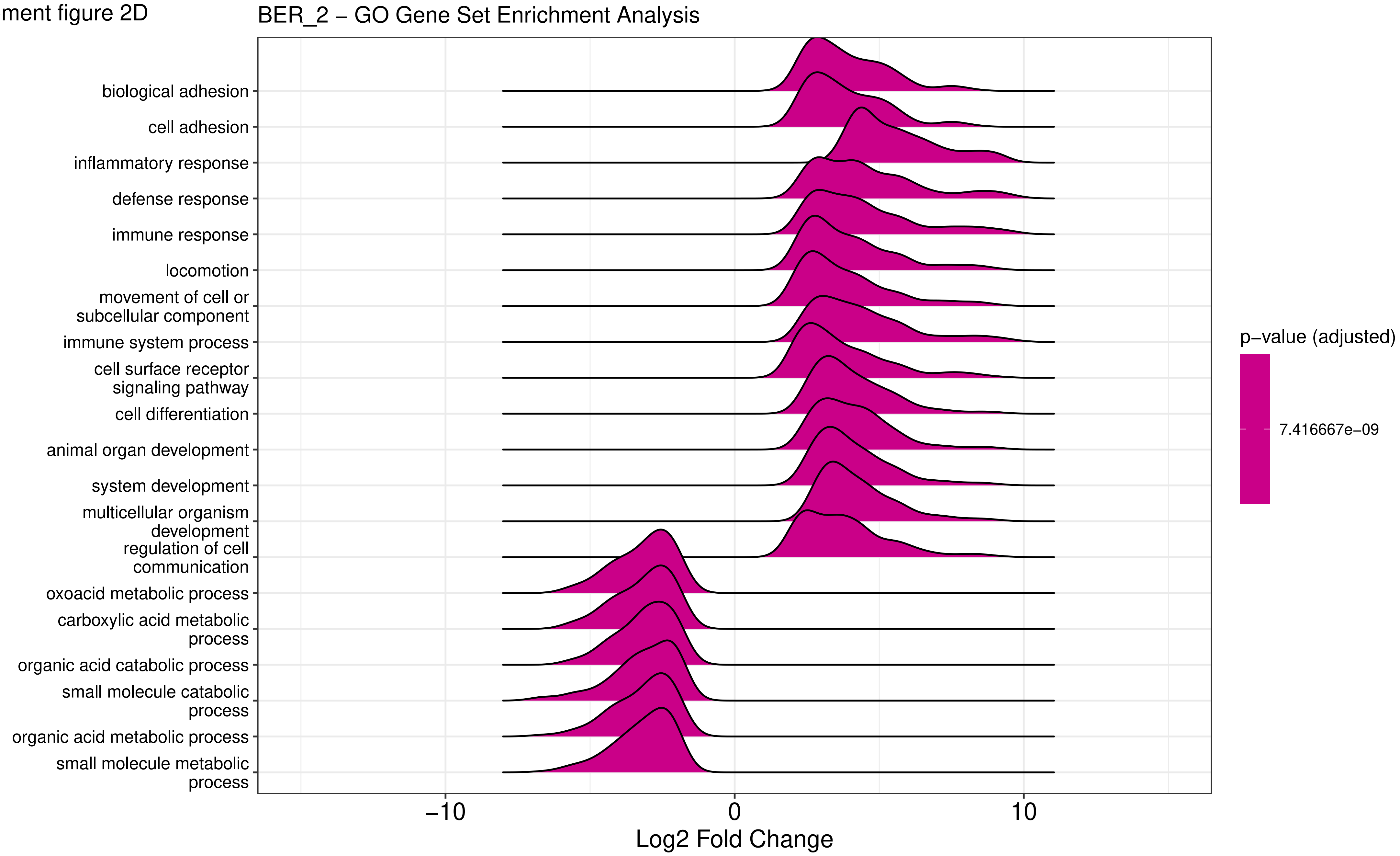

Supplement figure 2E

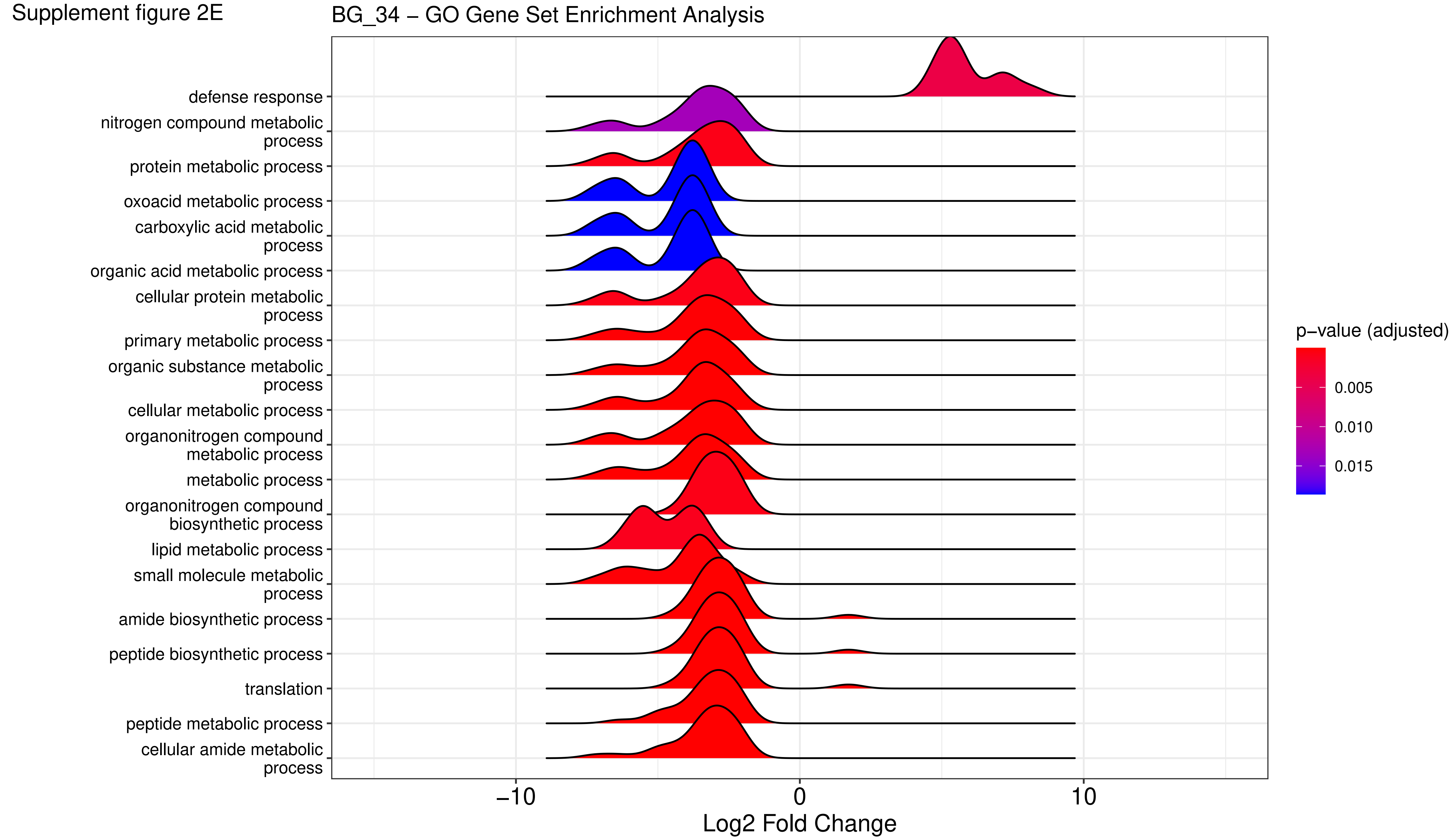

Supplement figure 2F

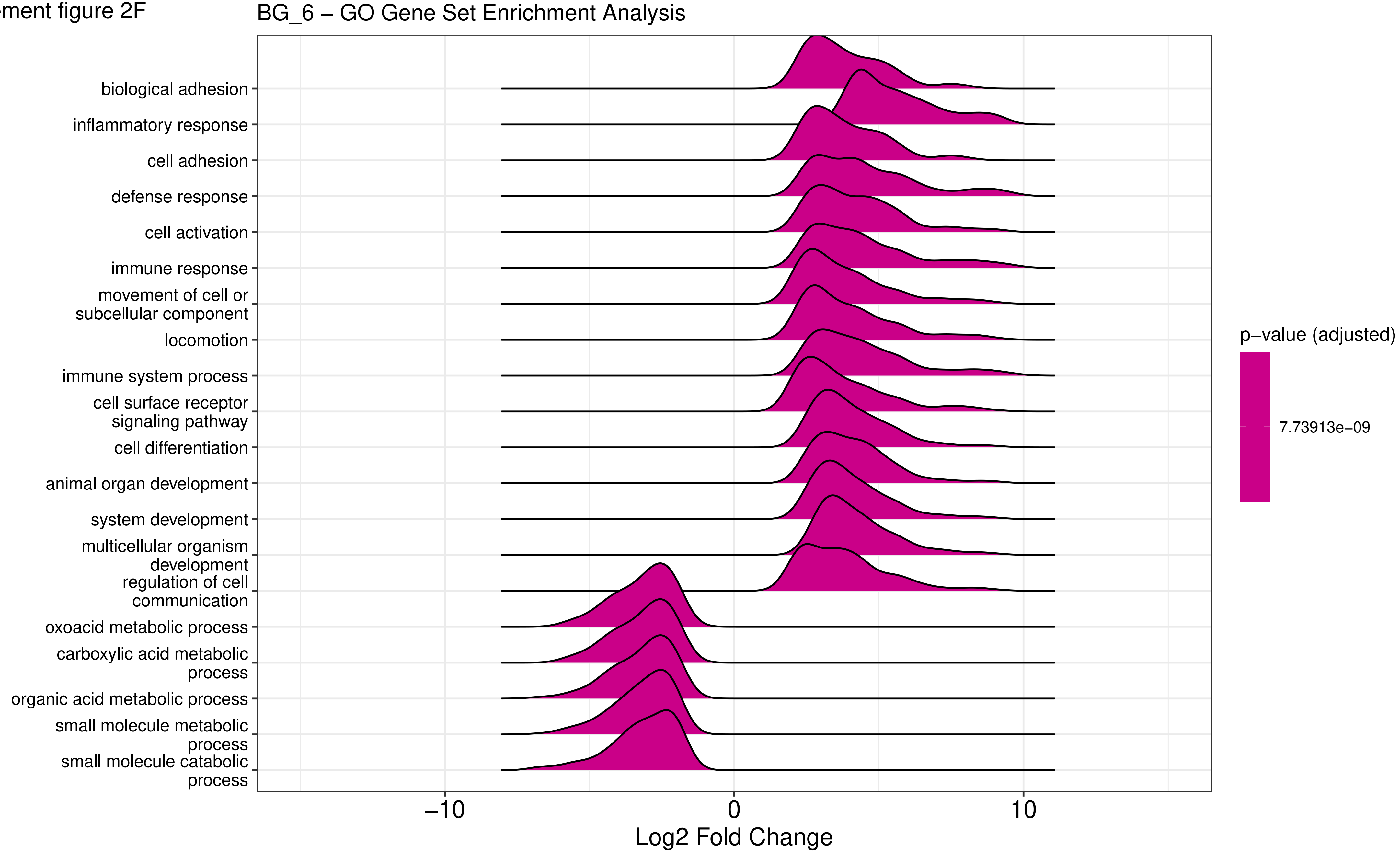

Supplement figure 2G

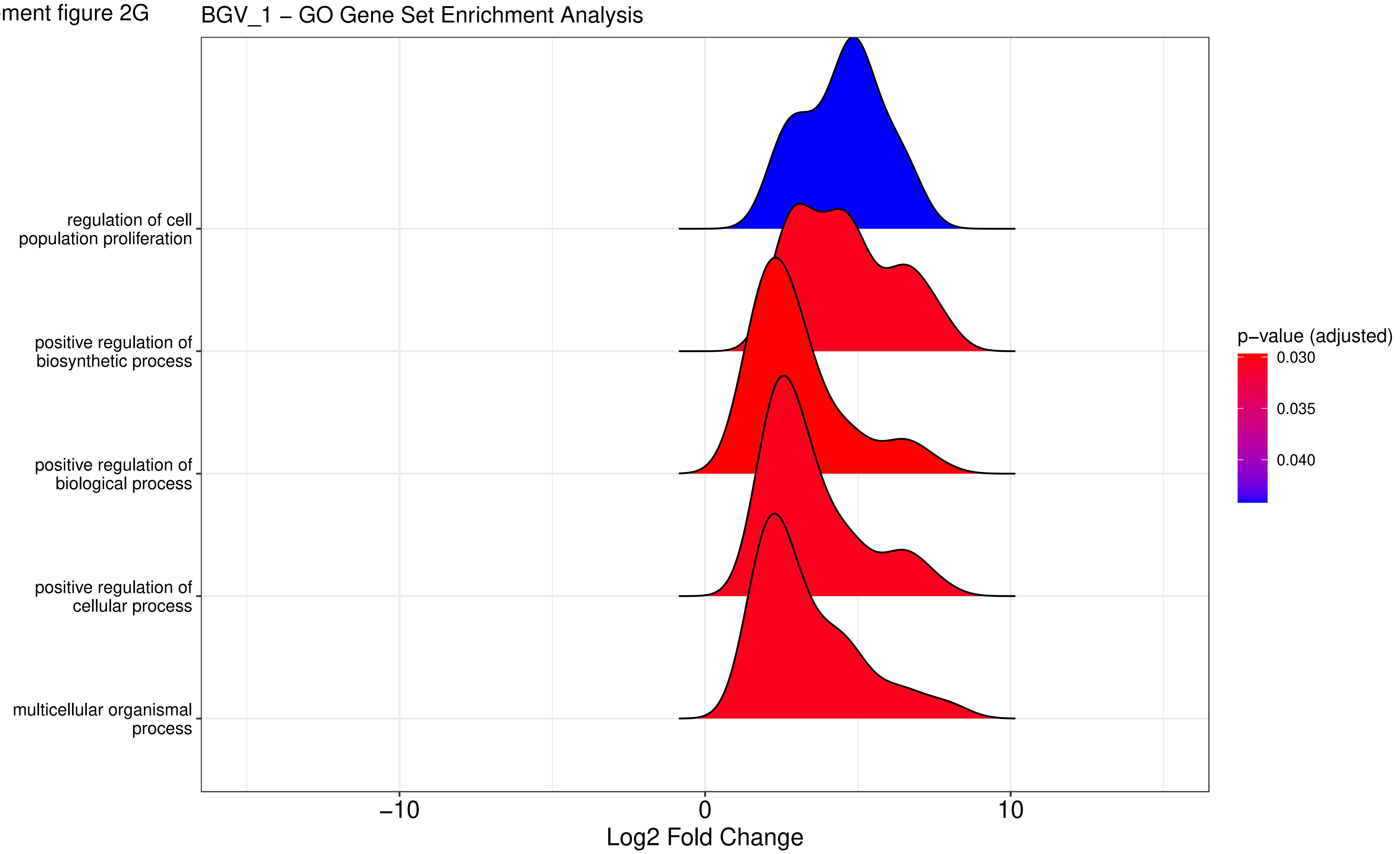

Supplement figure 2H

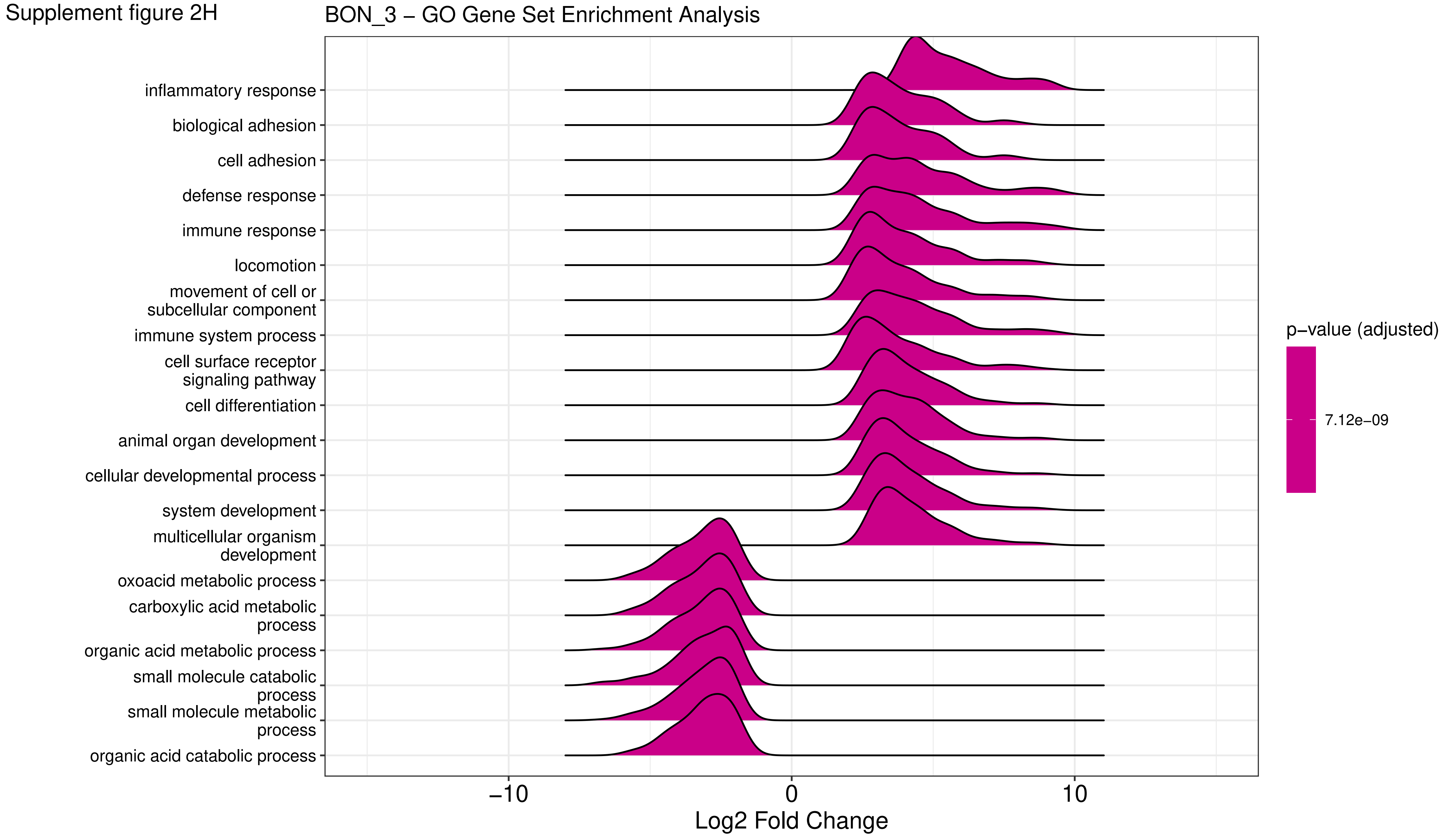

Supplement figure 2I

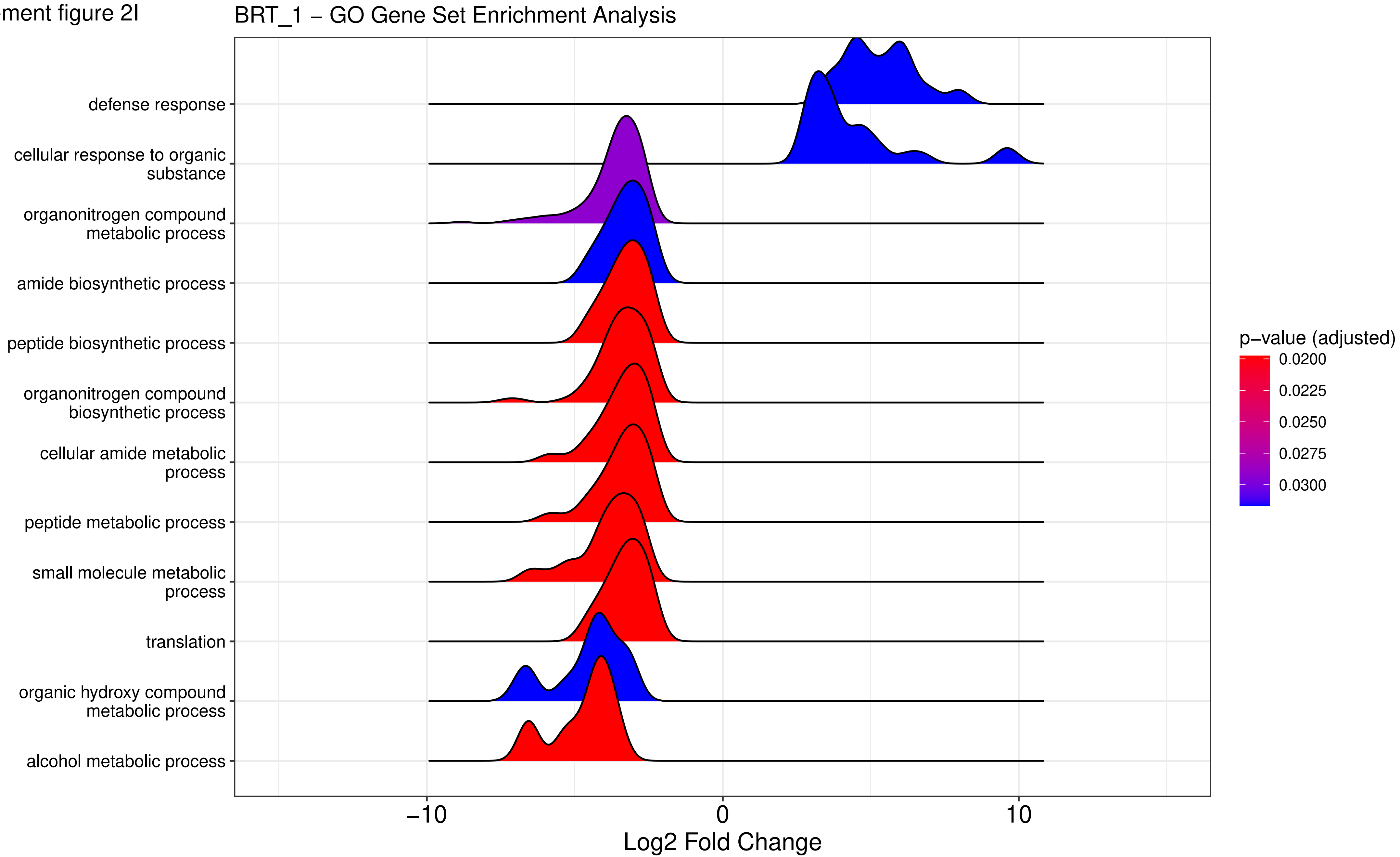

Supplement figure 2J

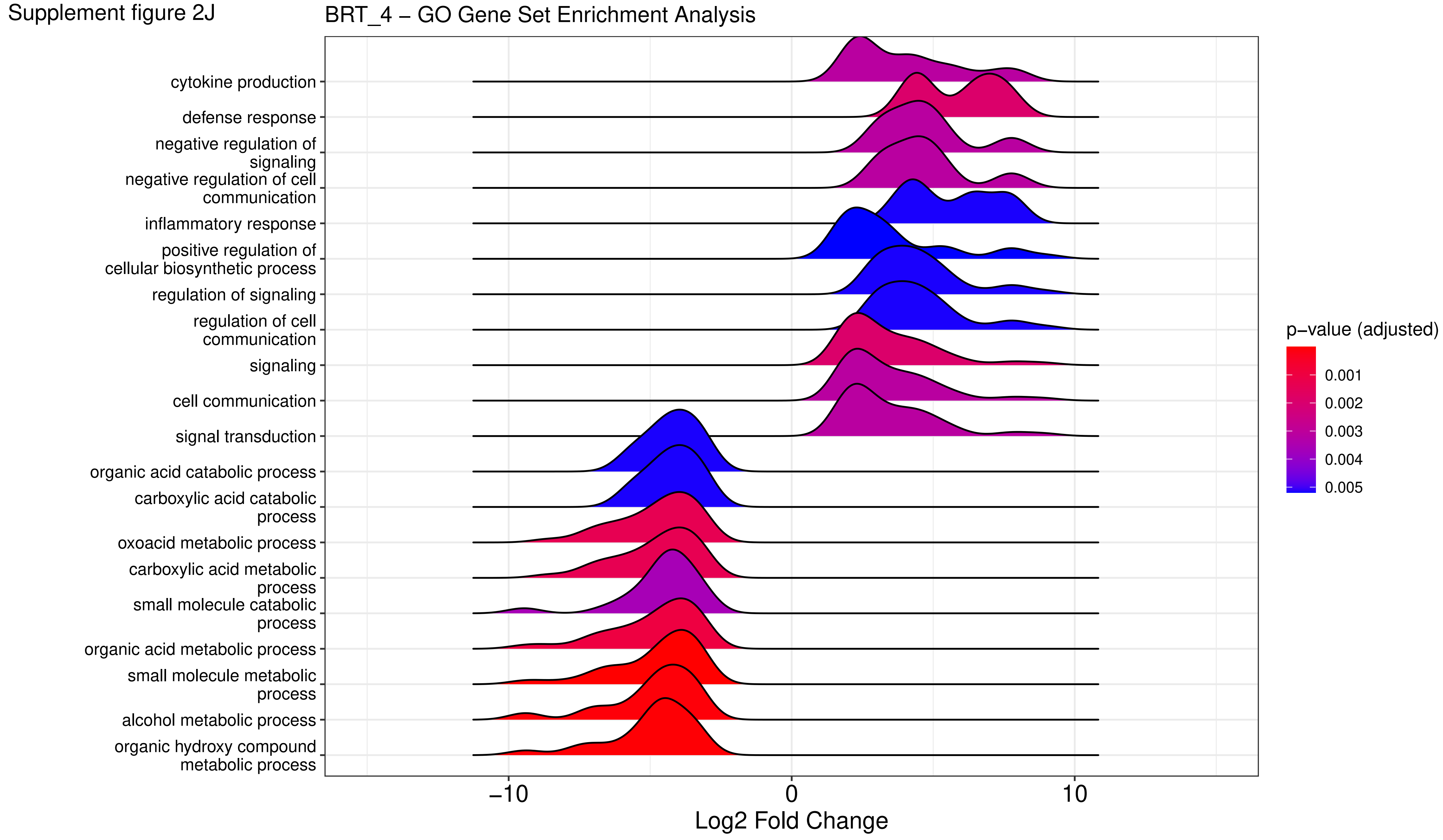

Supplement figure 2K

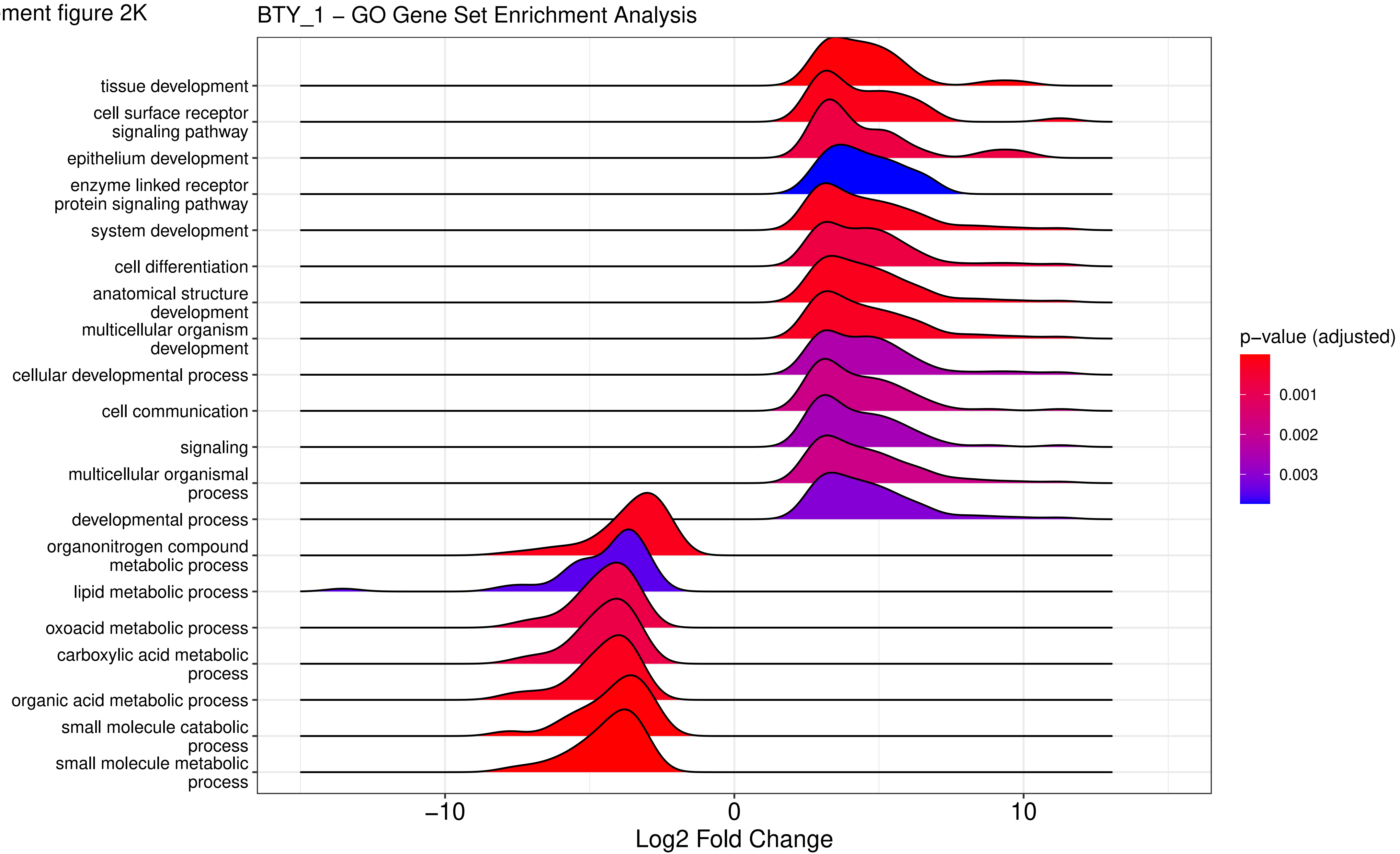

Supplement figure 2L

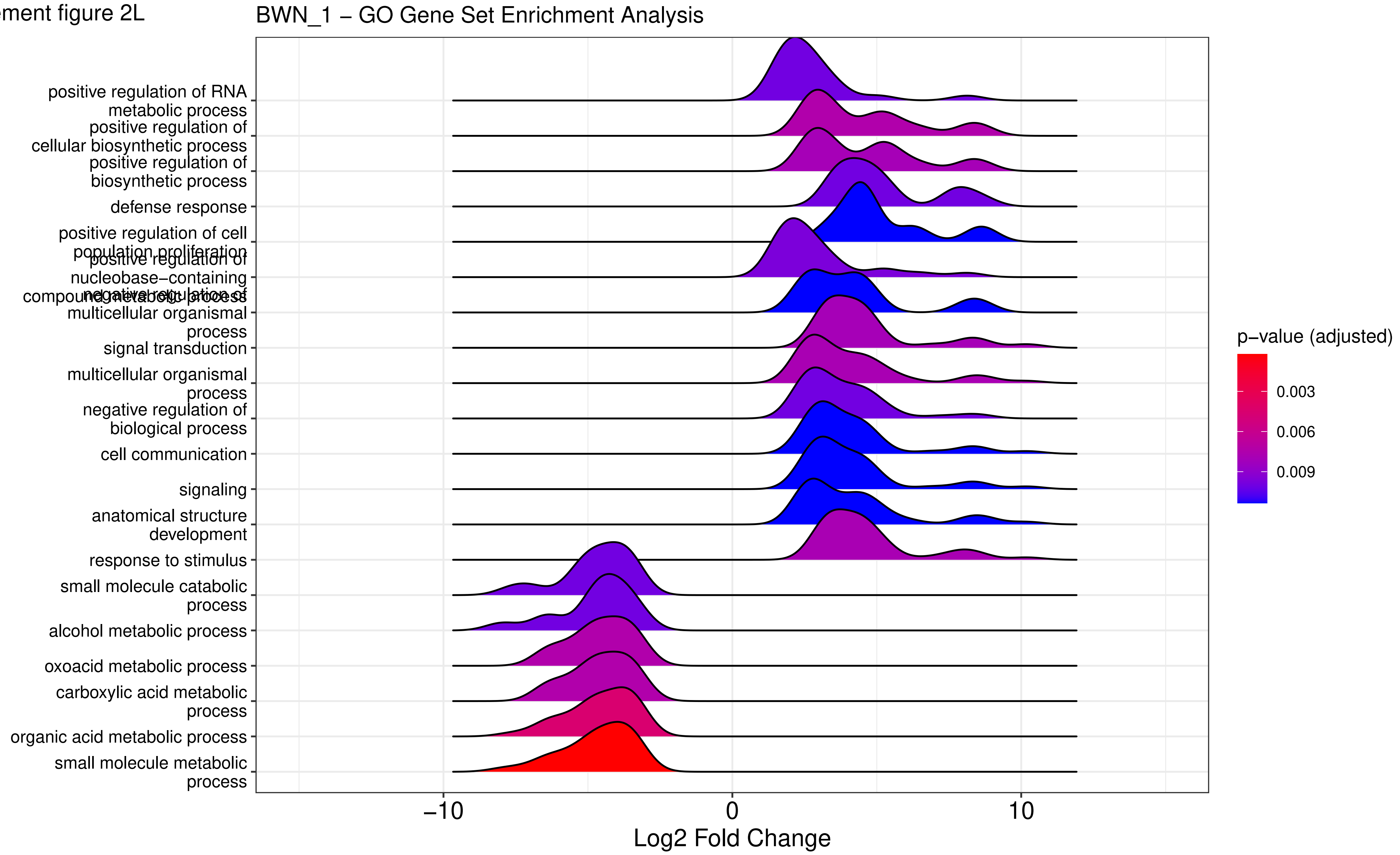

Supplement figure 2M

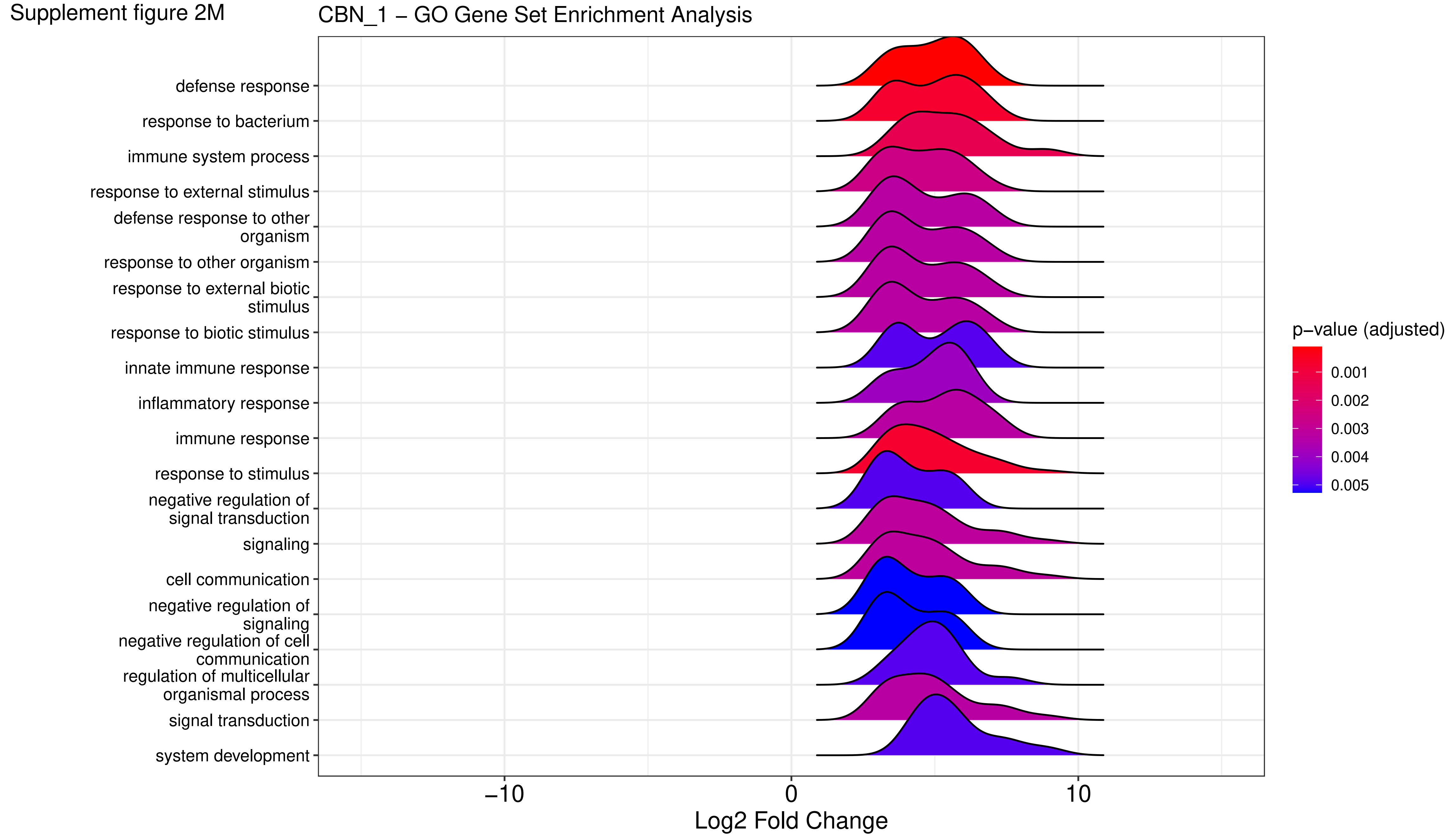

Supplement figure 2N

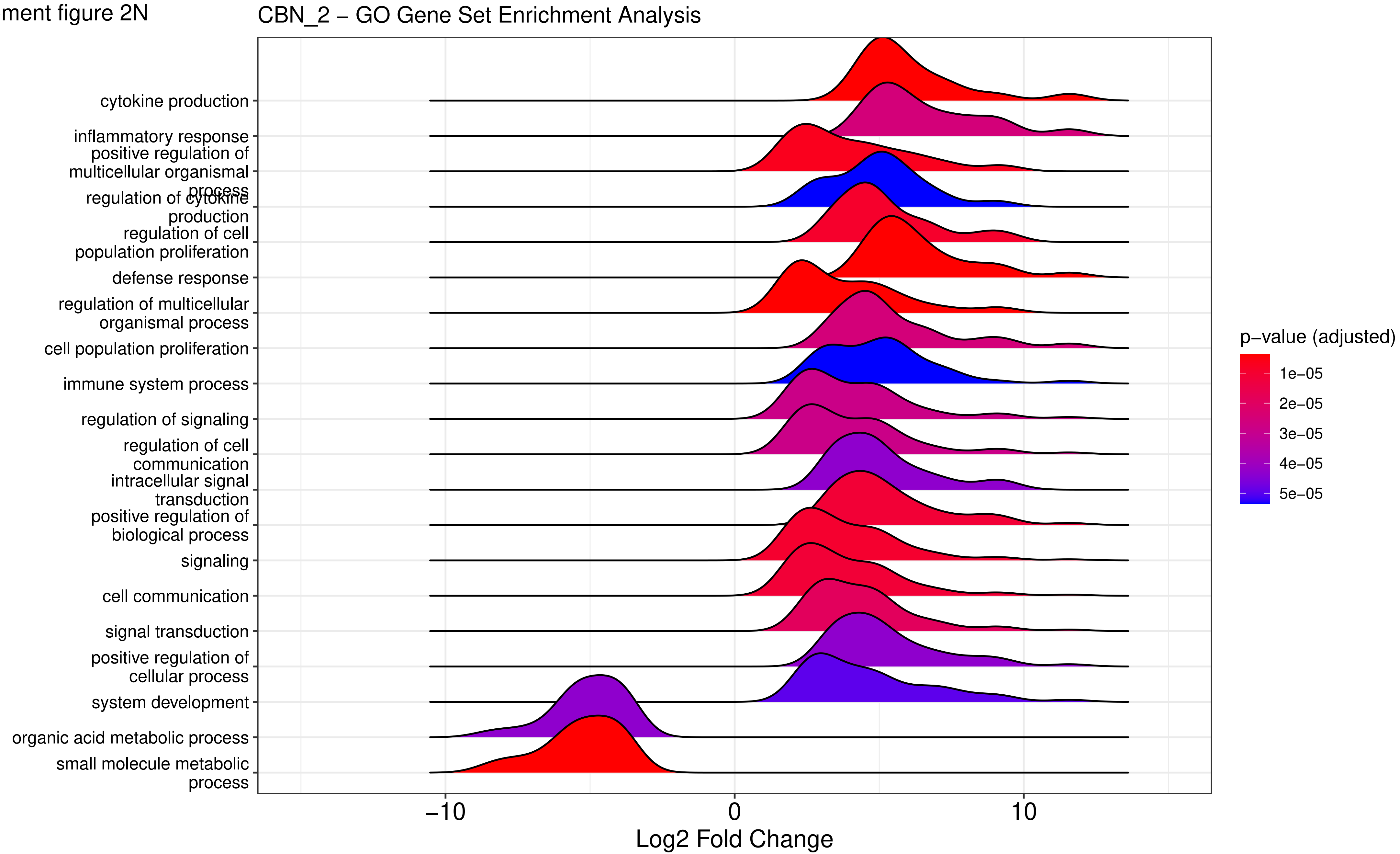

Supplement figure 2O

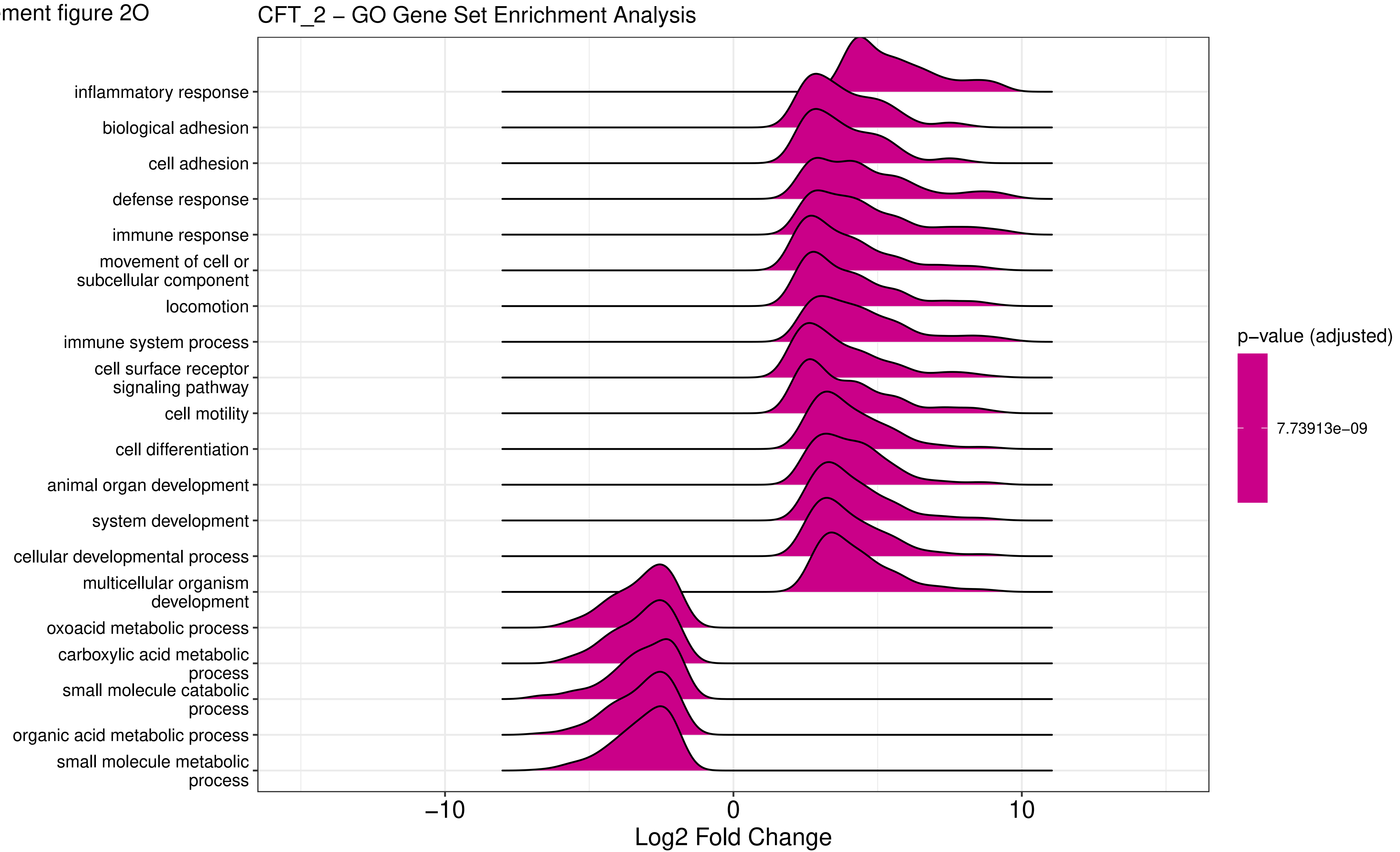

Supplement figure 2P

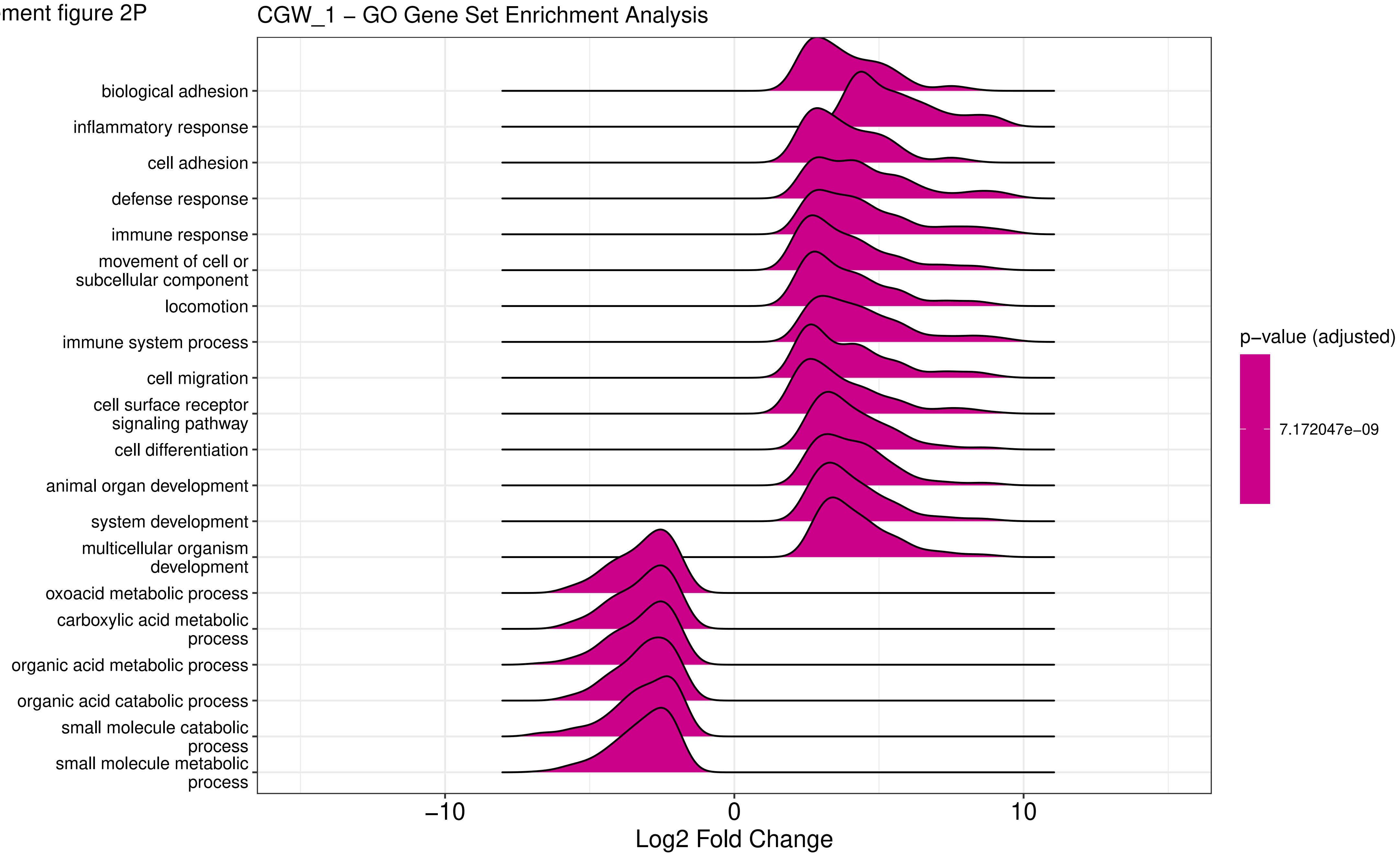

Supplement figure 2Q

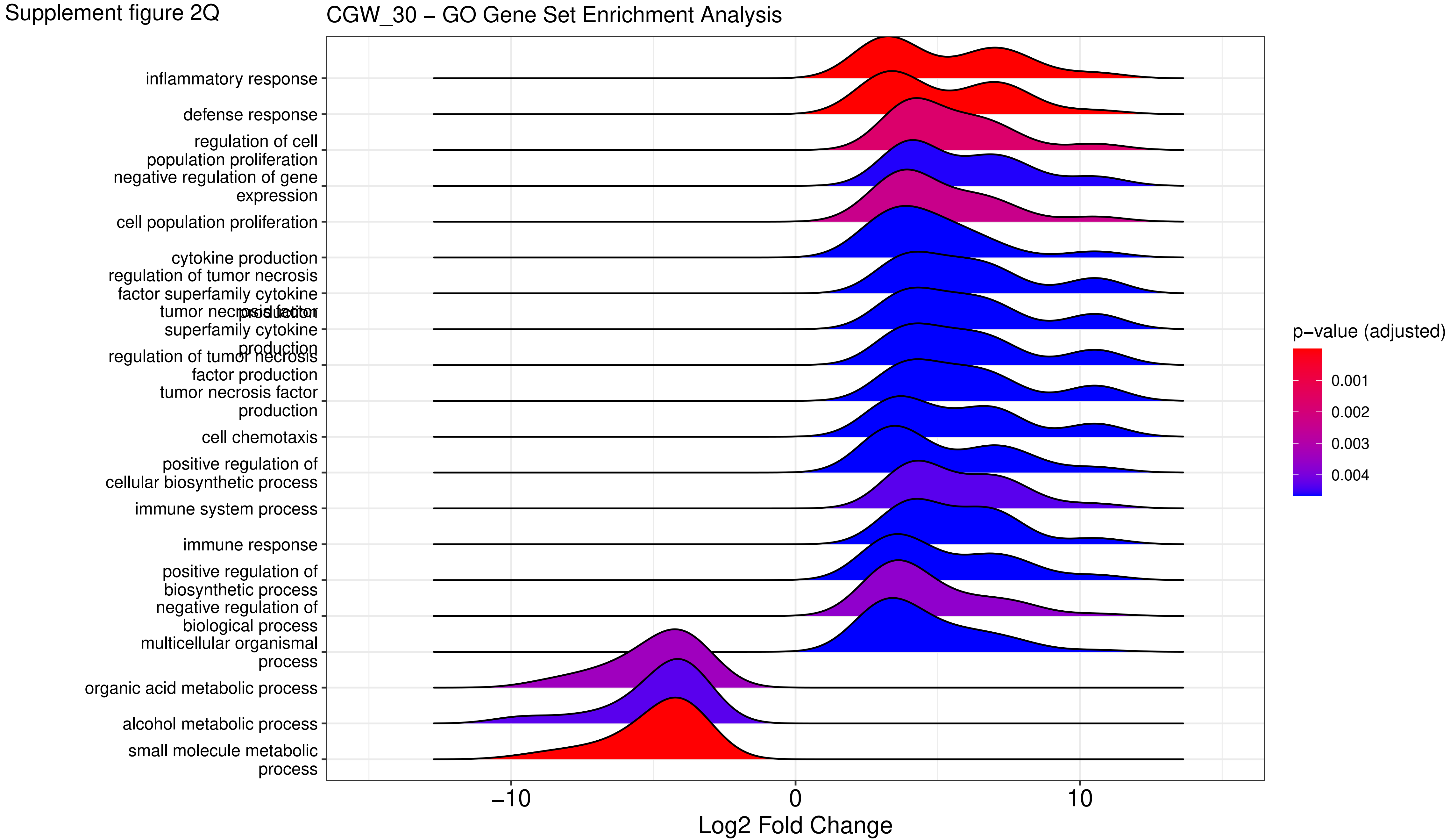

Supplement figure 2R

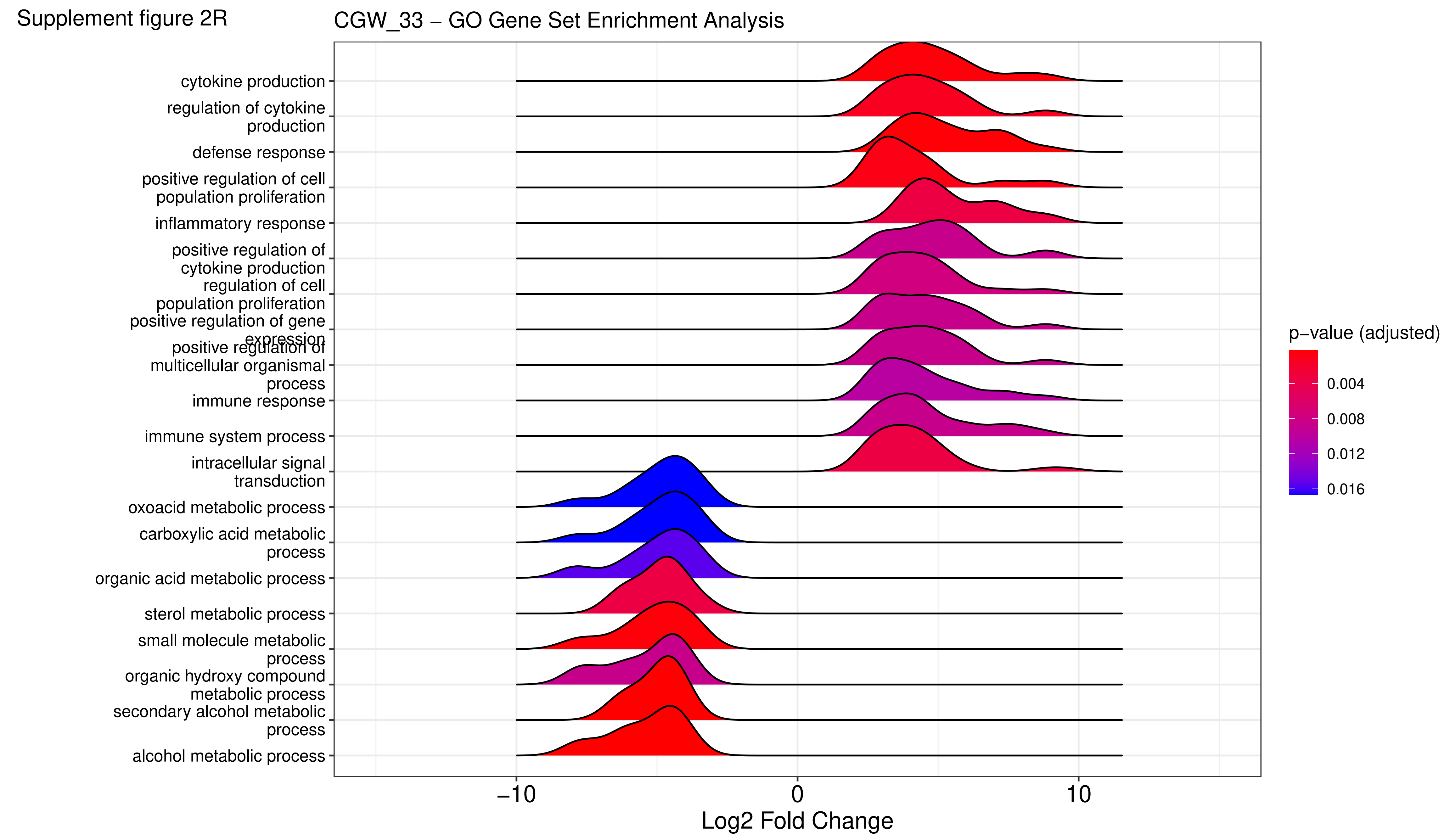

Supplement figure 2S

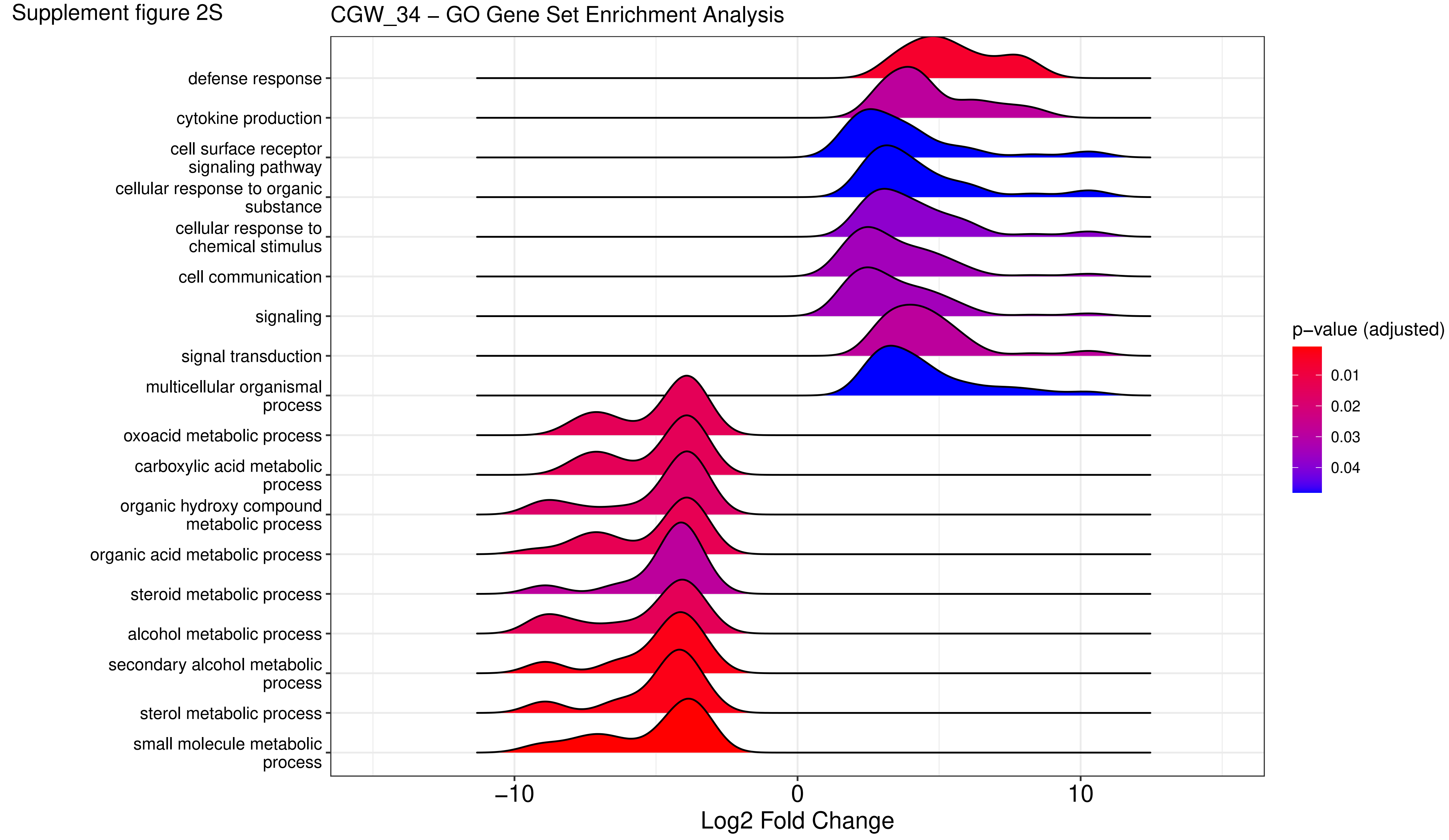

Supplement figure 2T

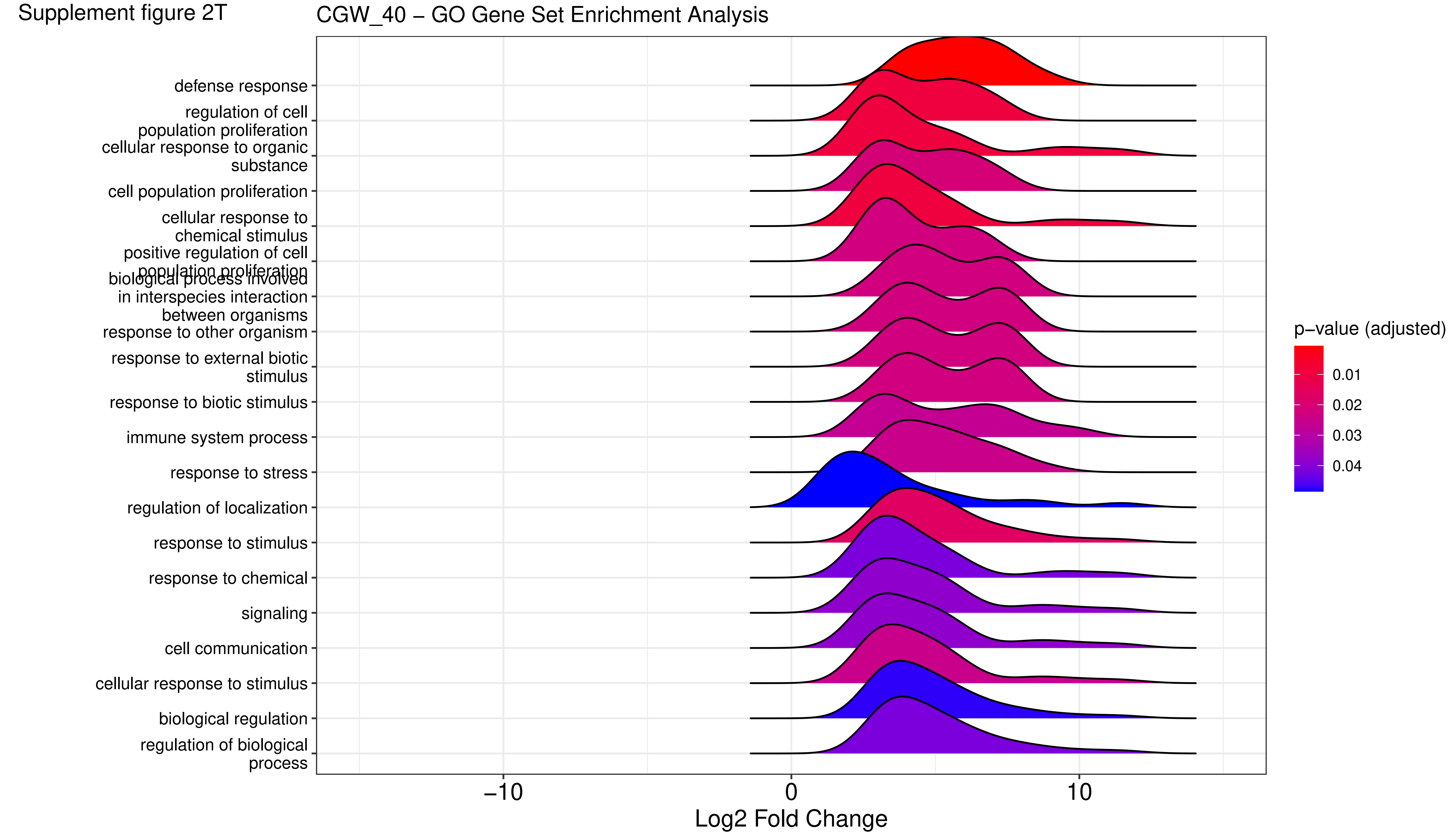

Supplement figure 2U

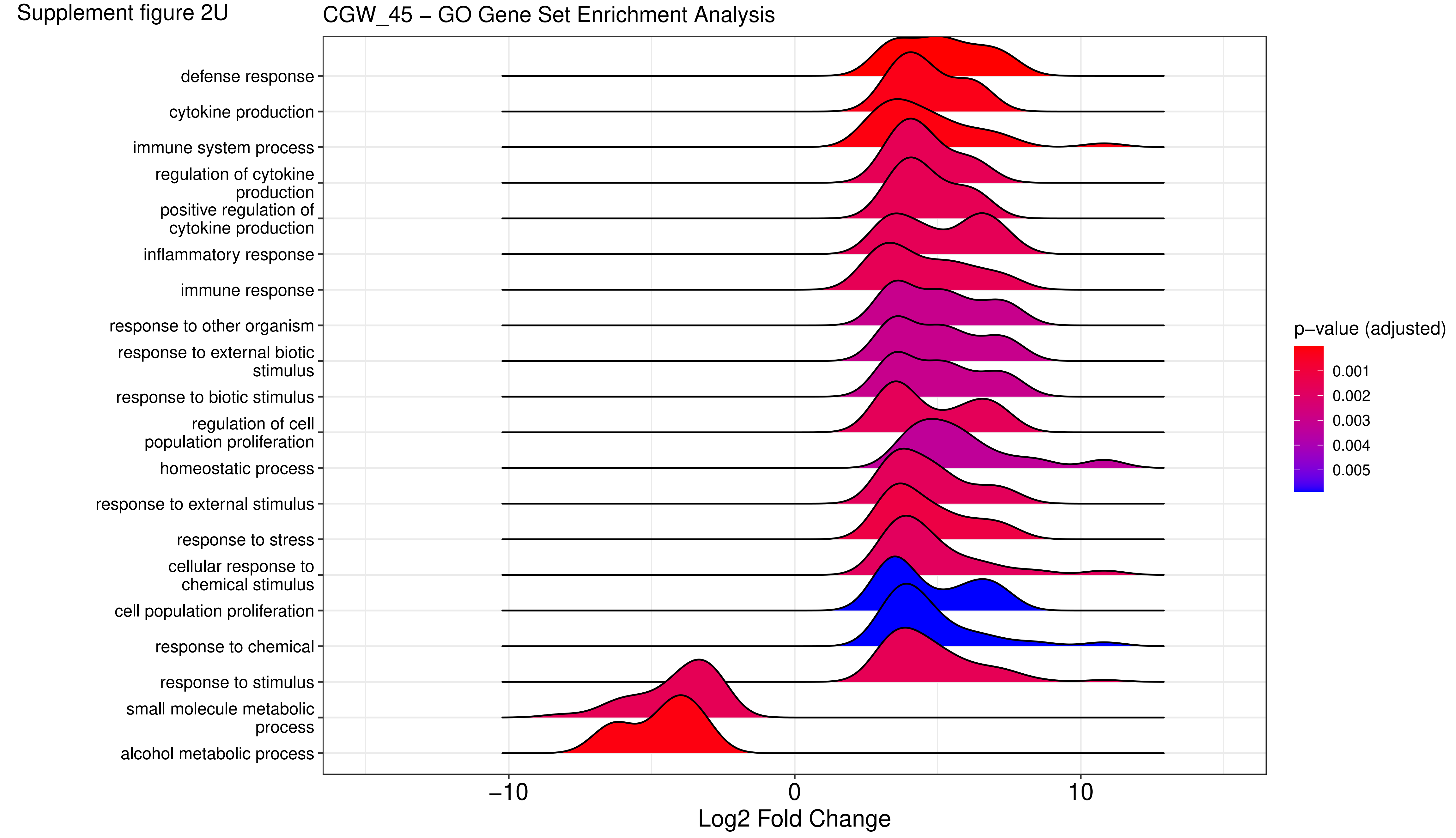

Supplement figure 2V

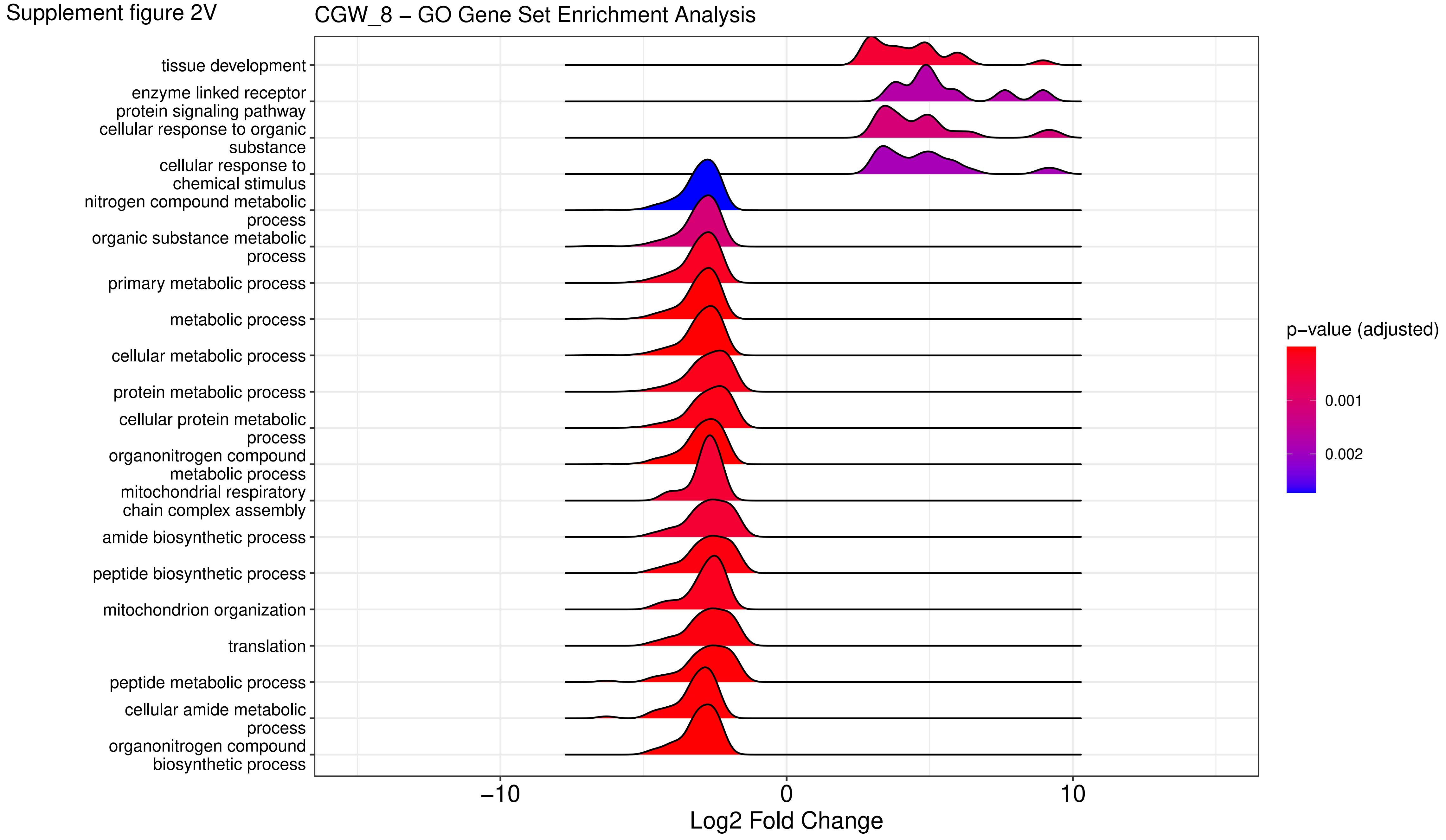

Supplement figure 2W

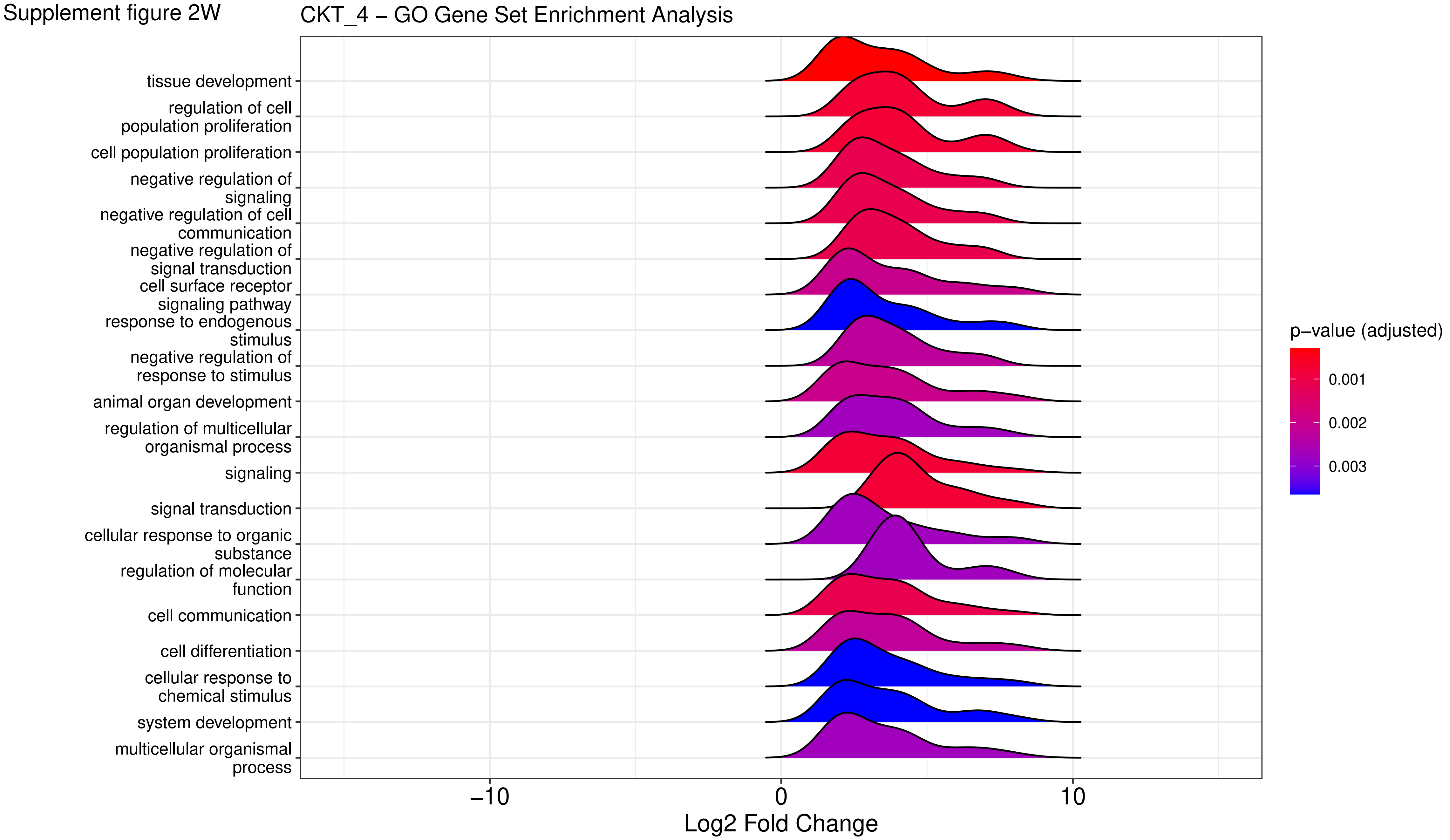

Supplement figure 2X

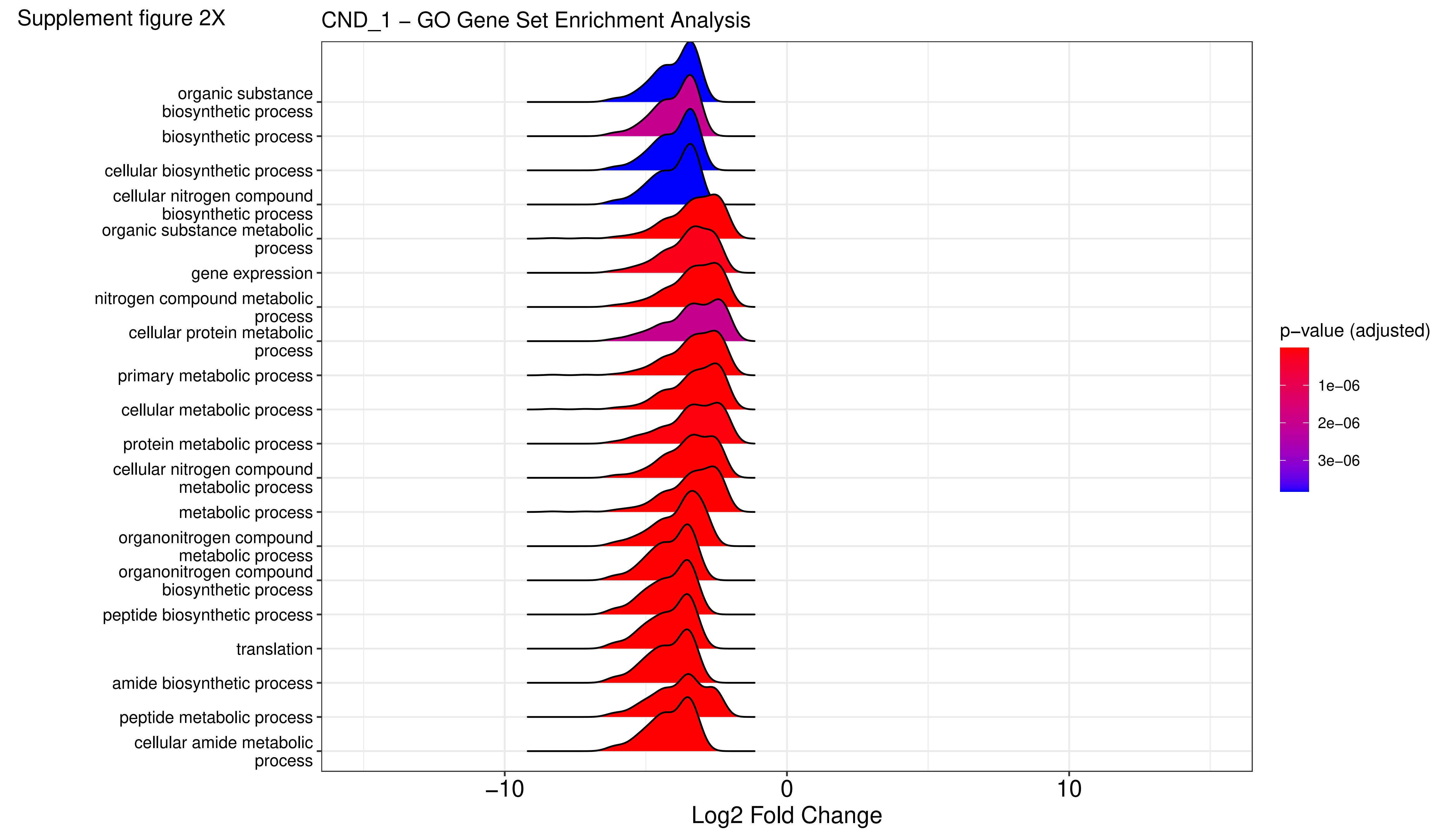

Supplement figure 2Y

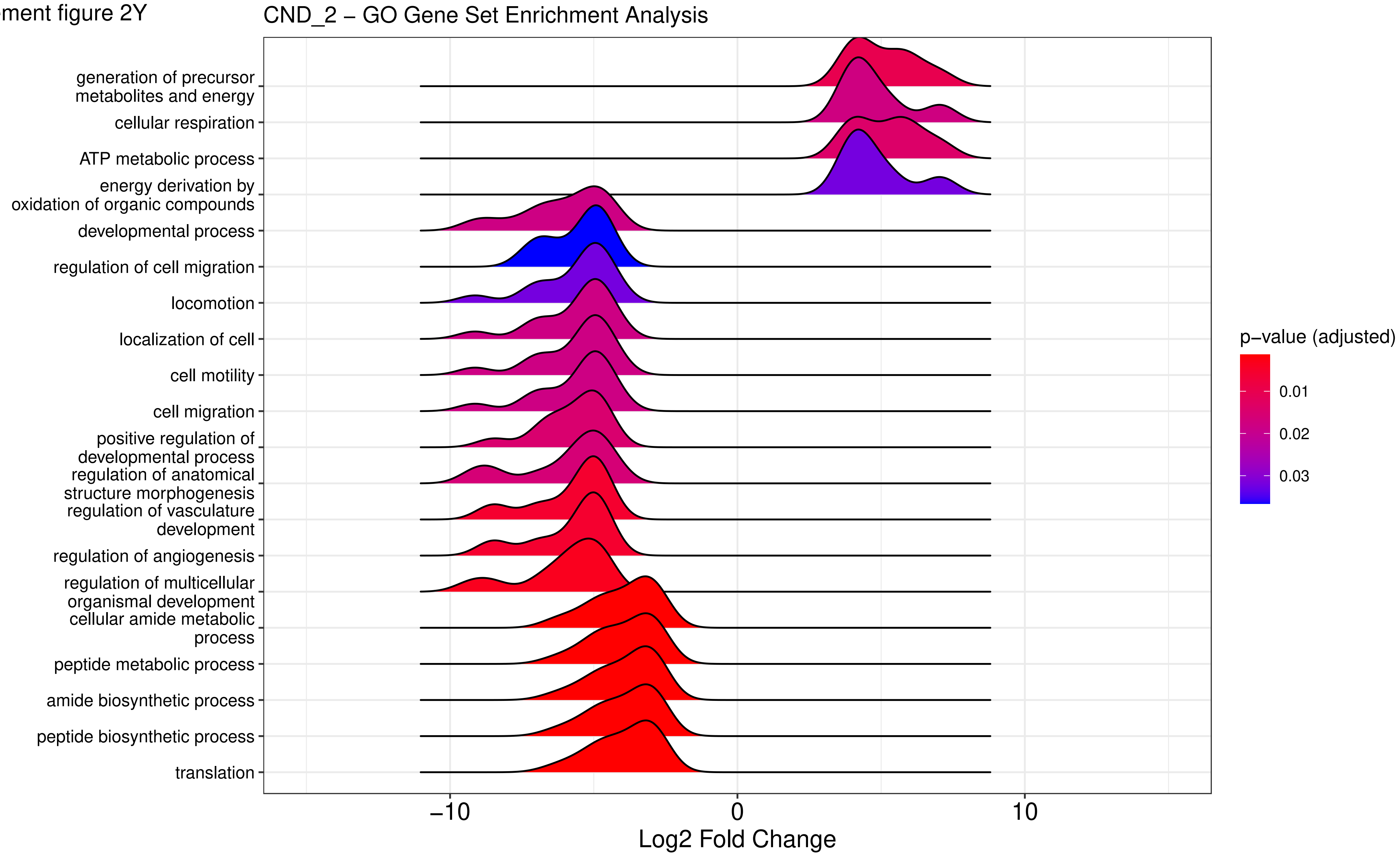

Supplement figure 2Z

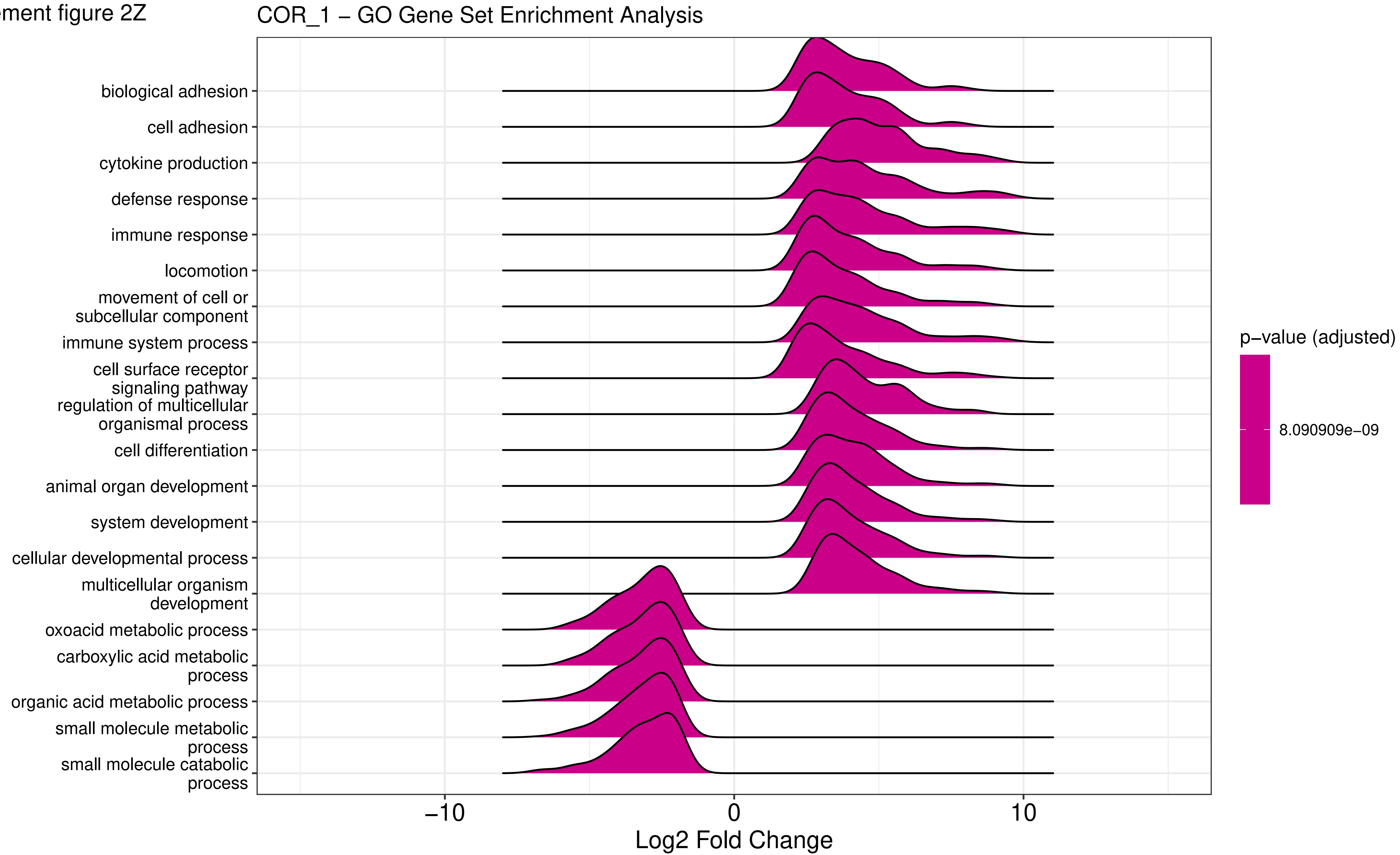

Supplement figure 2AA

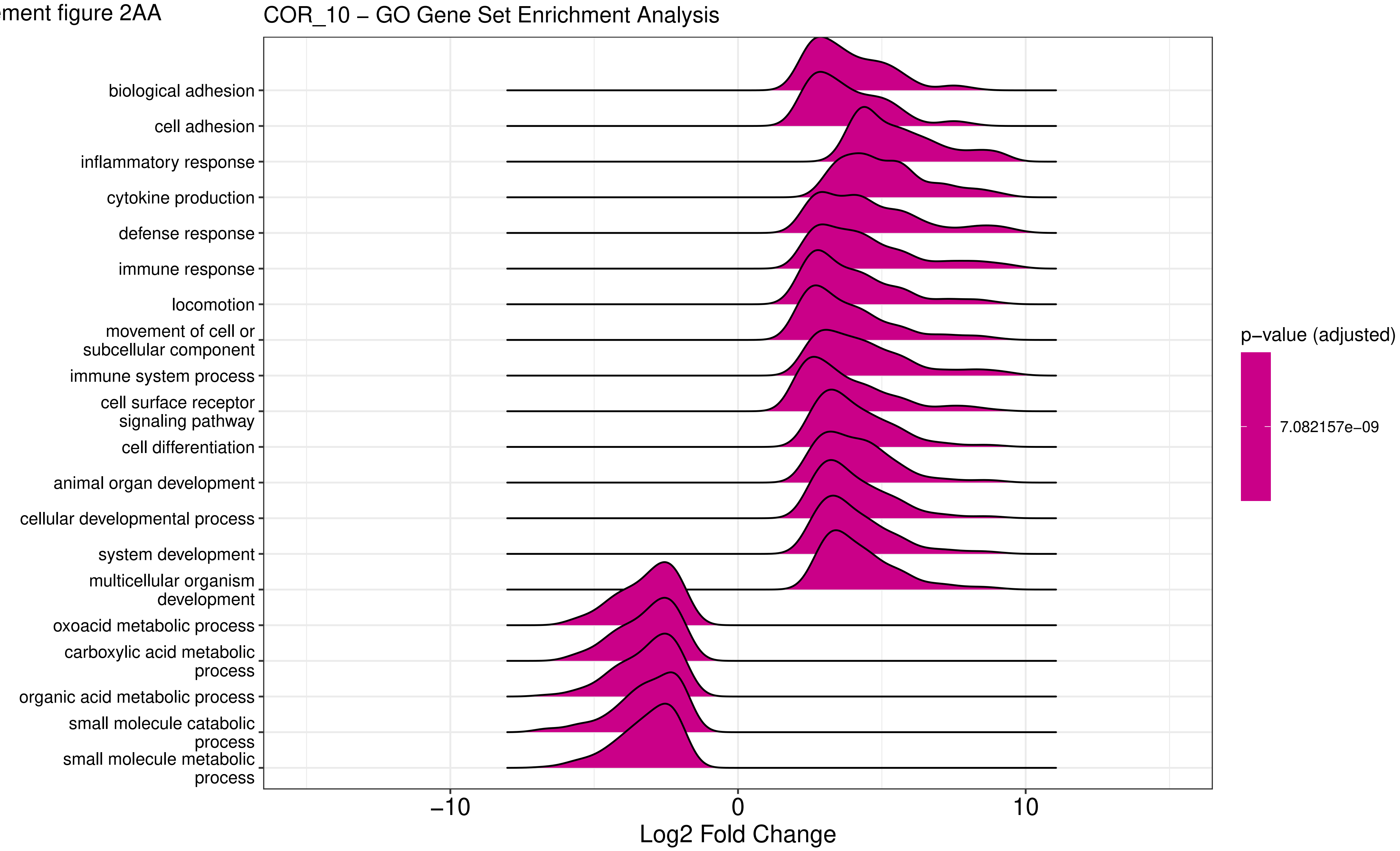

Supplement figure 2AB

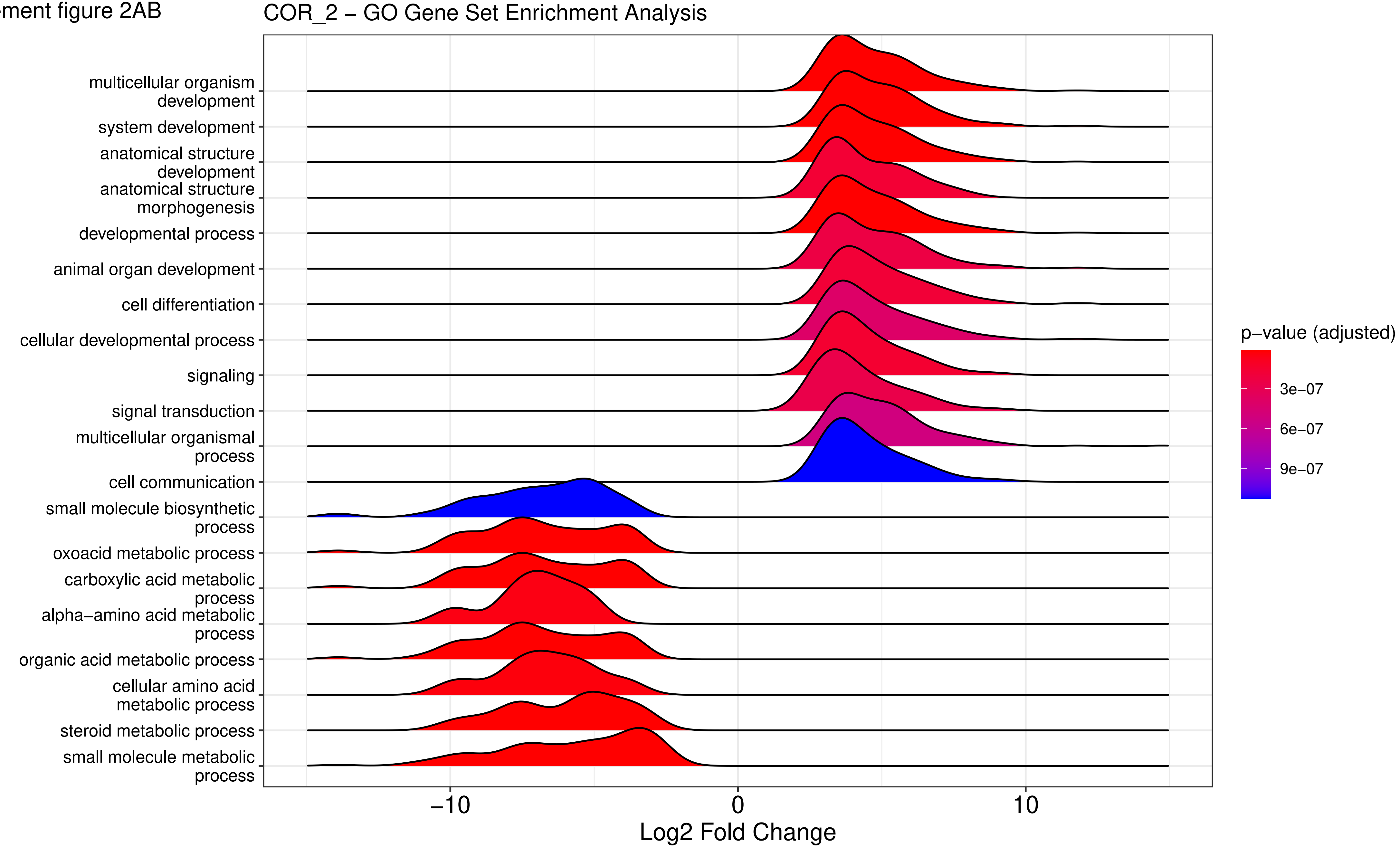

Supplement figure 2AC

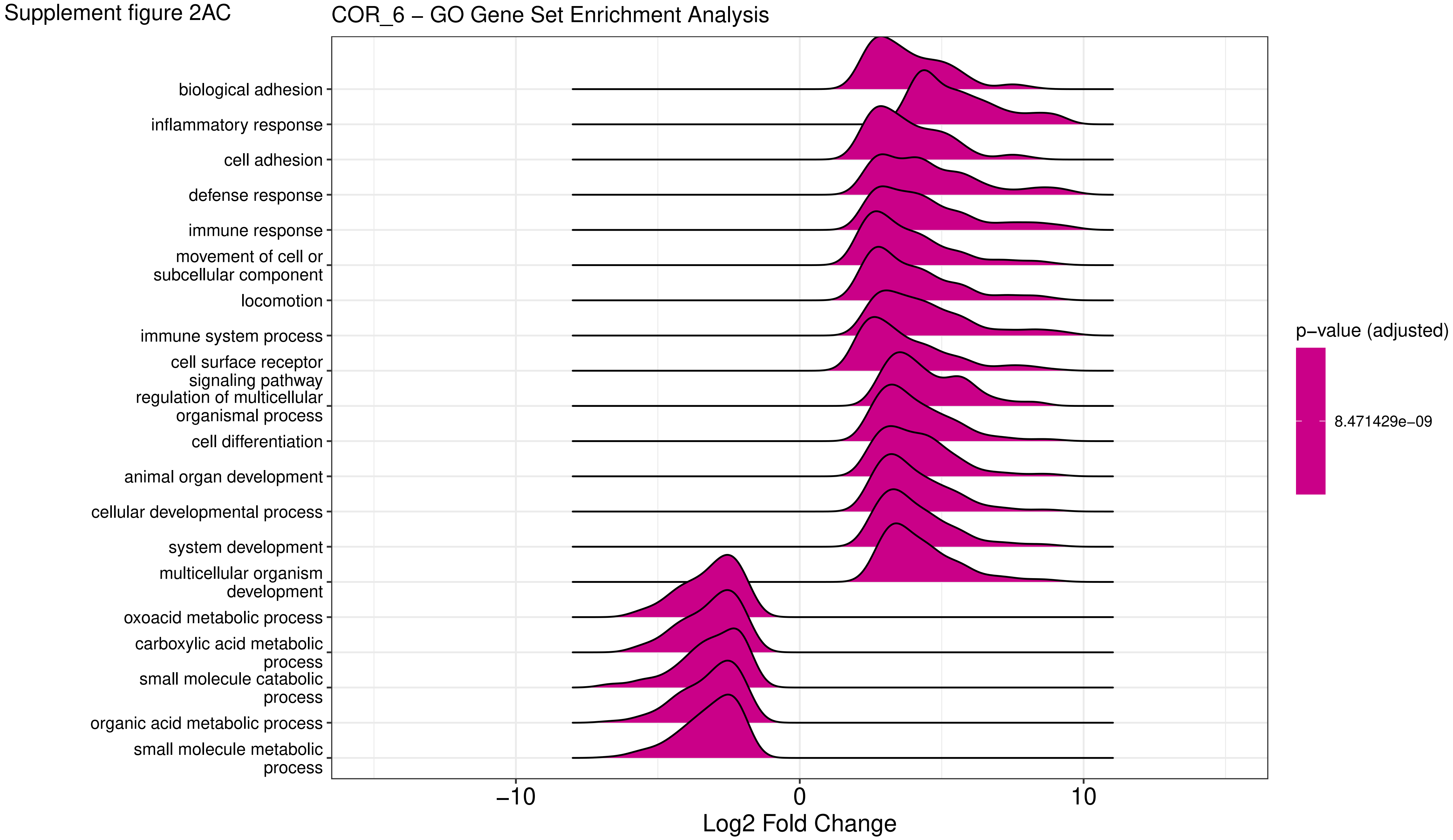

Supplement figure 2AD

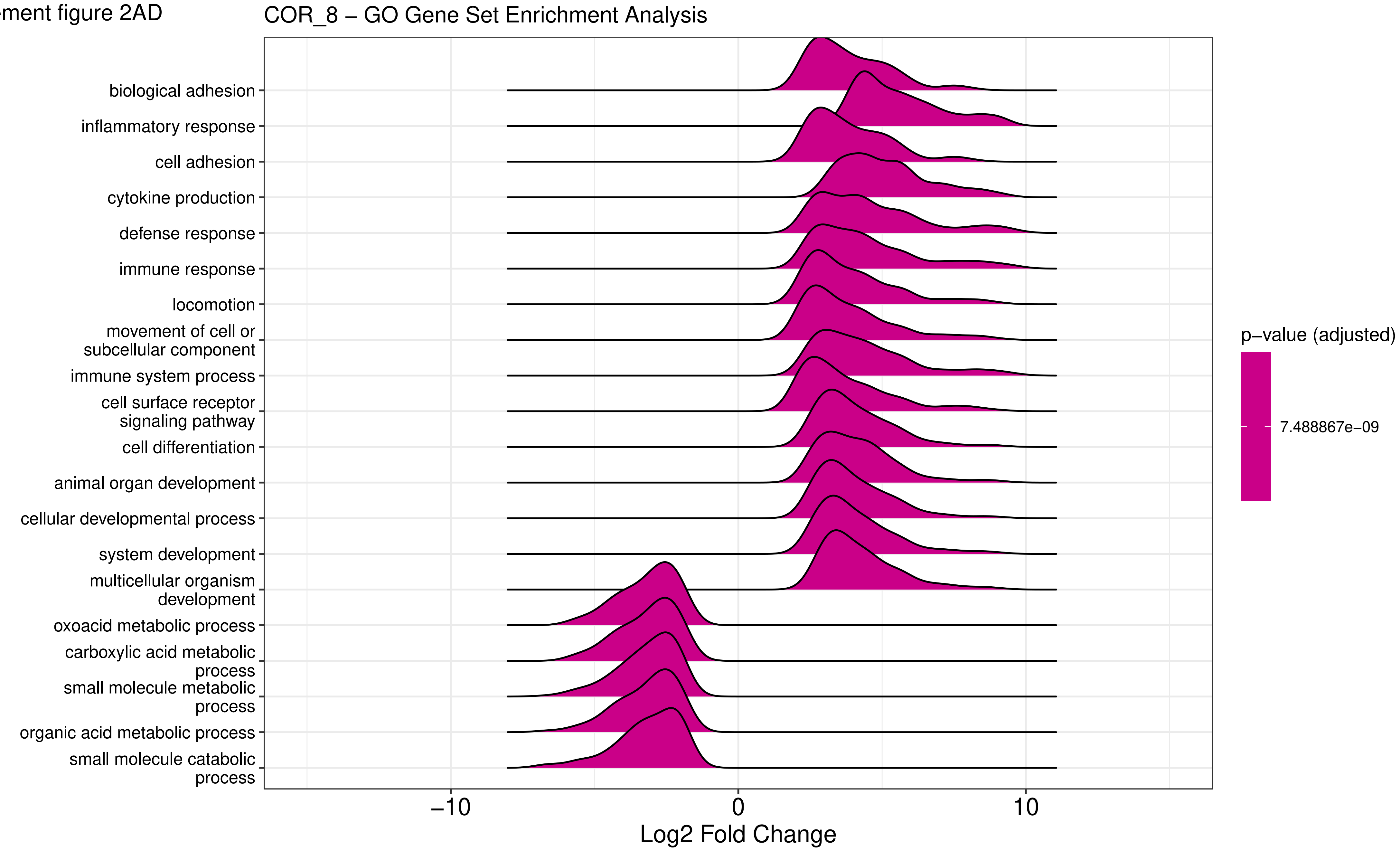

Supplement figure 2AE

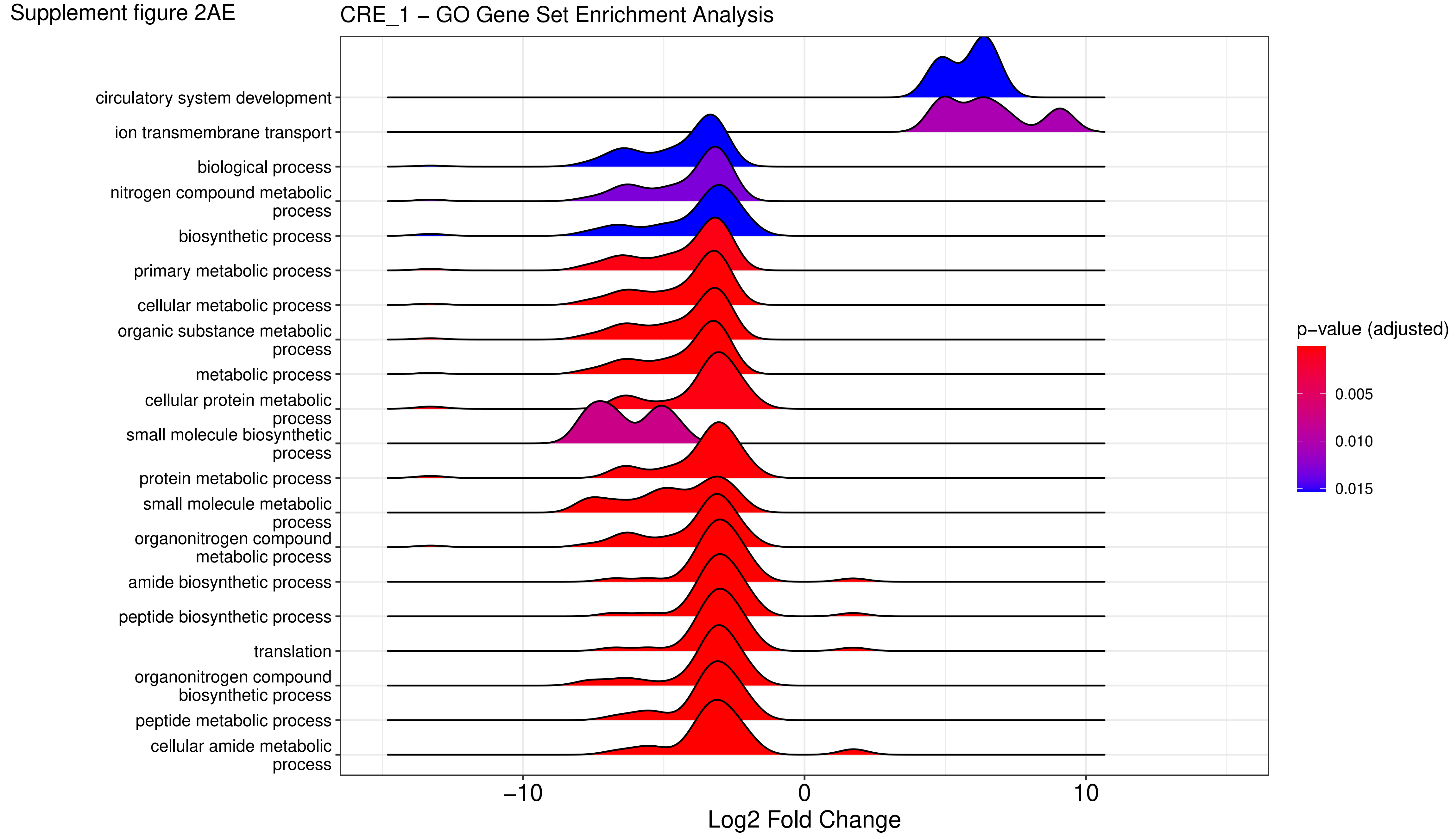

Supplement figure 2AF

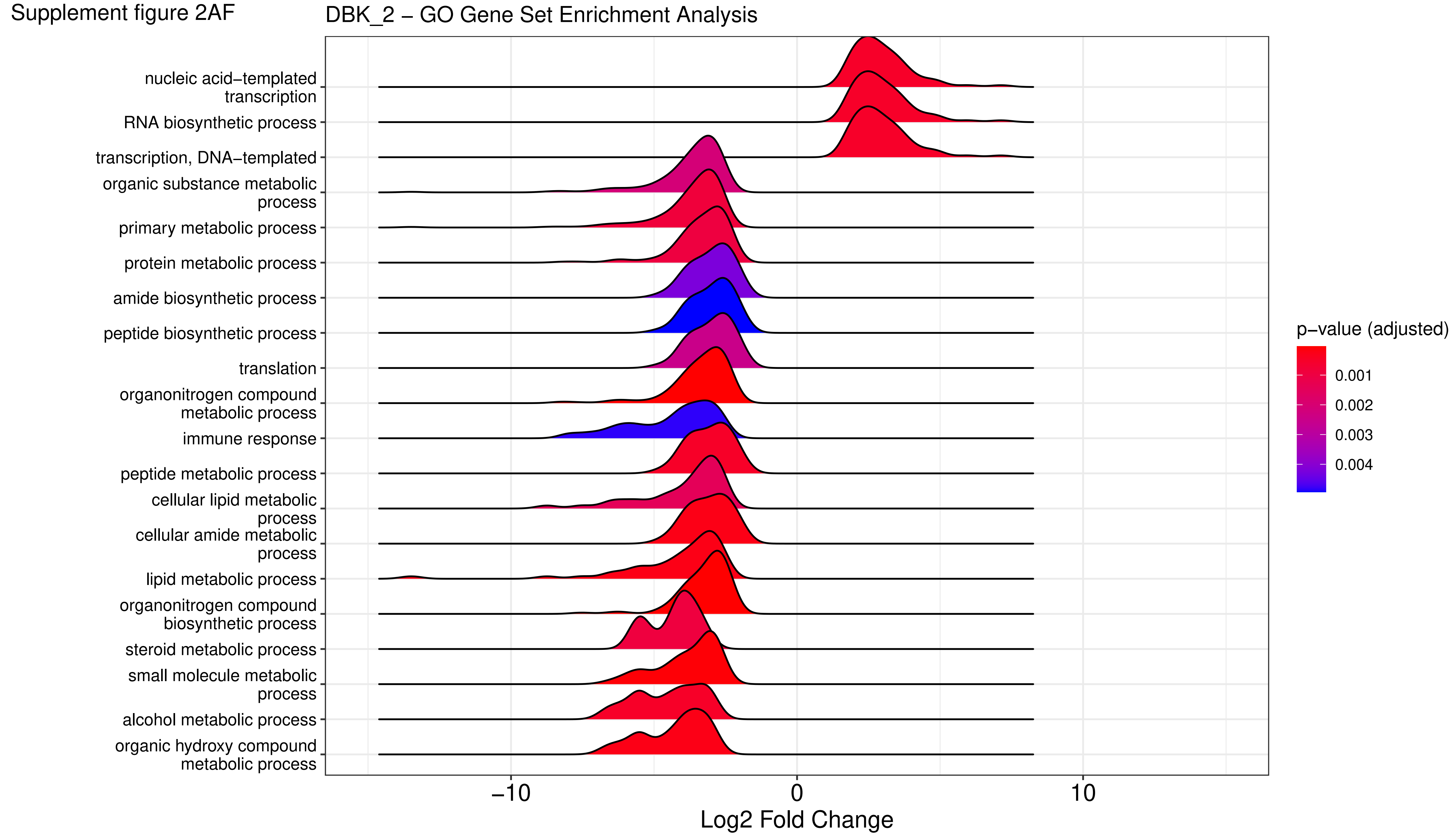

Supplement figure 2AG

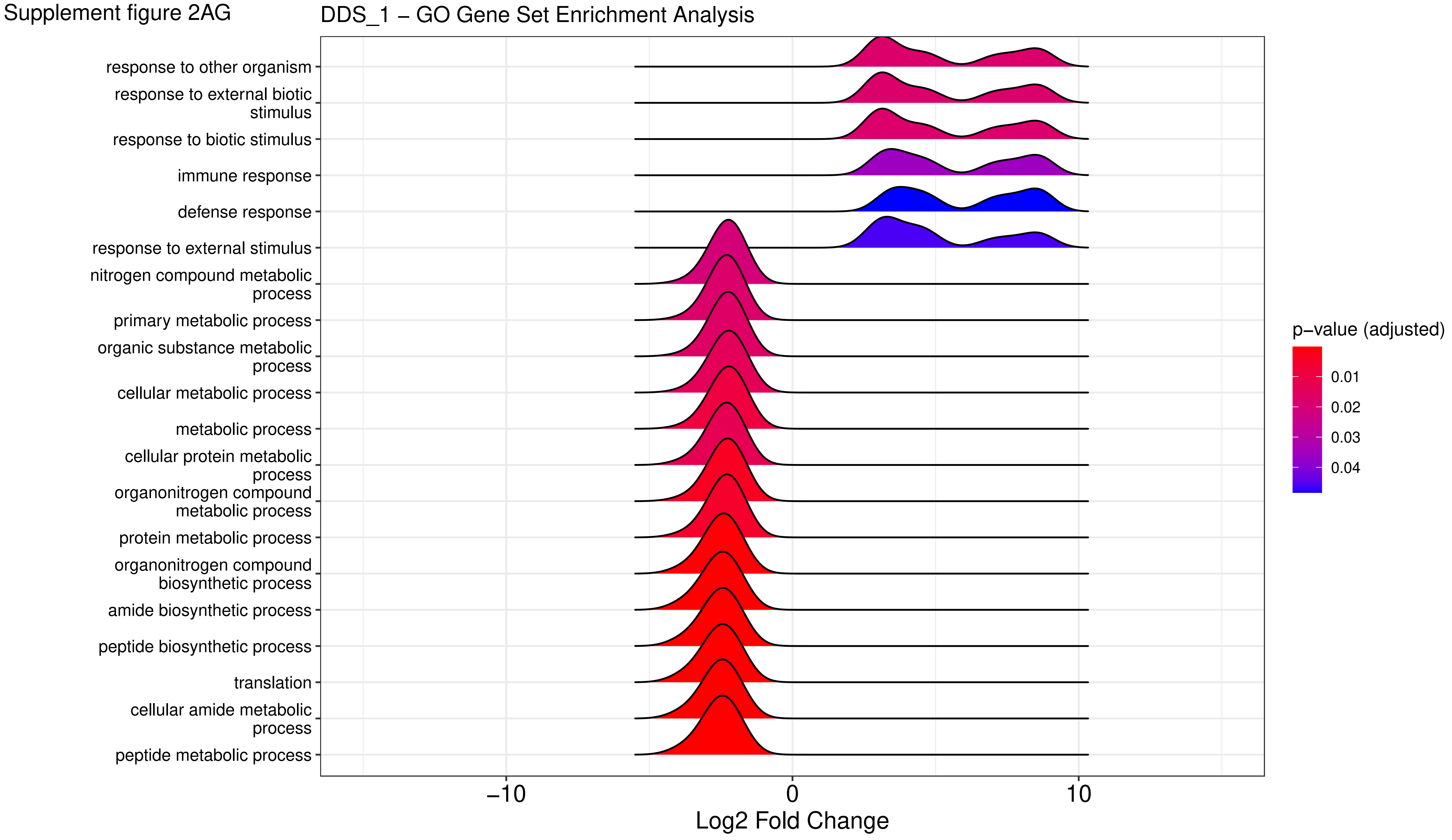

Supplement figure 2AH

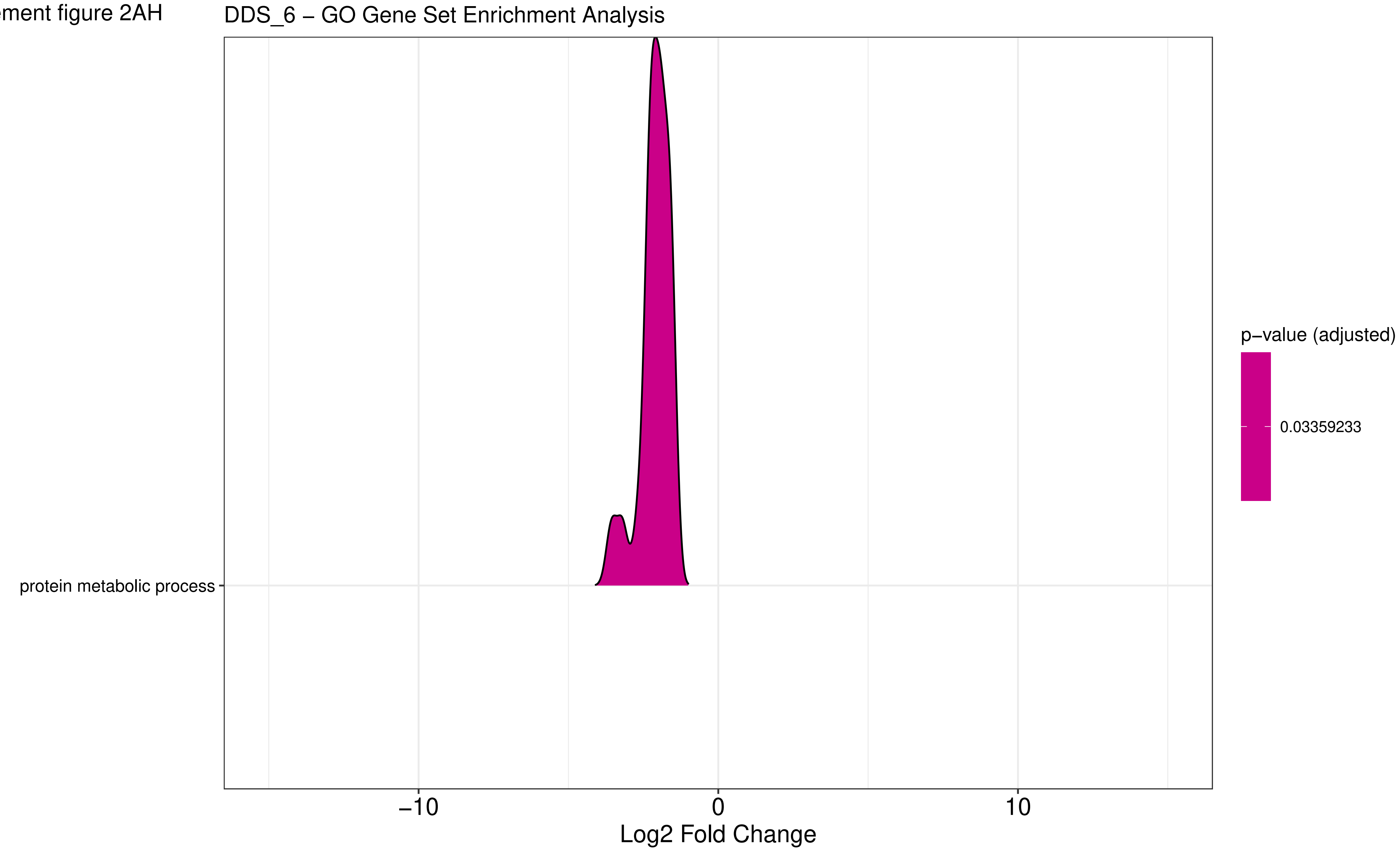

Supplement figure 2AI

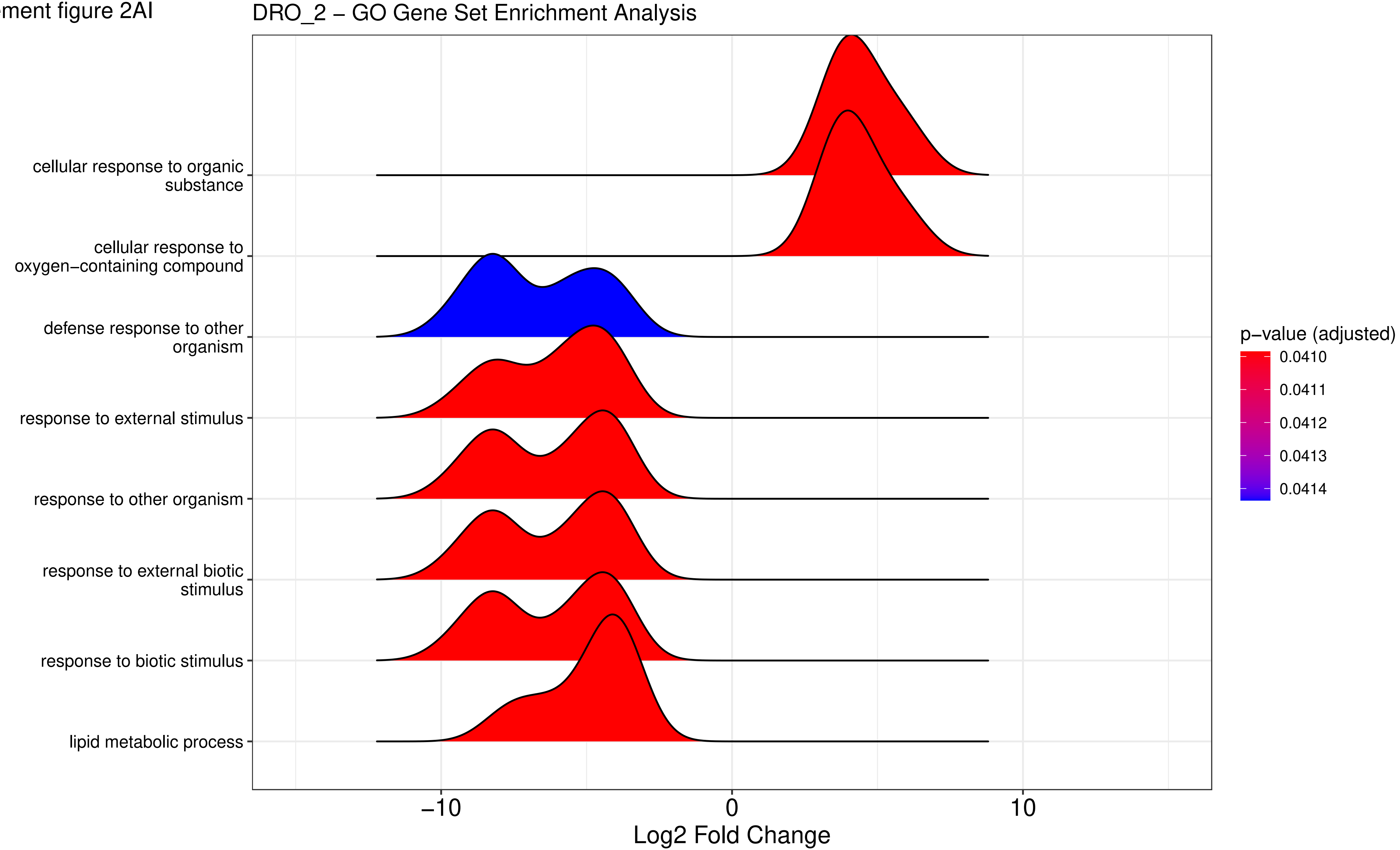

Supplement figure 2AJ

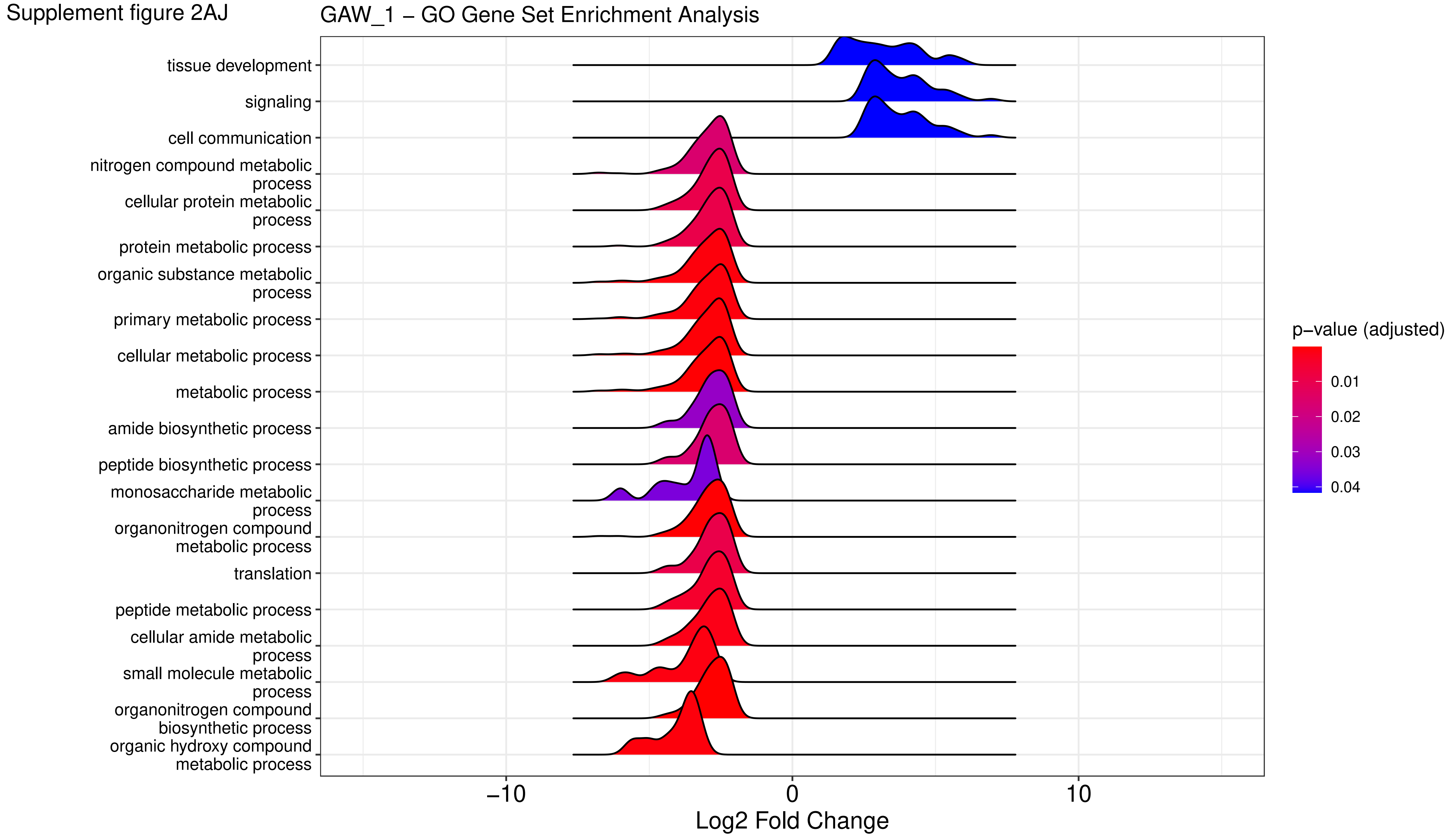

Supplement figure 2AK

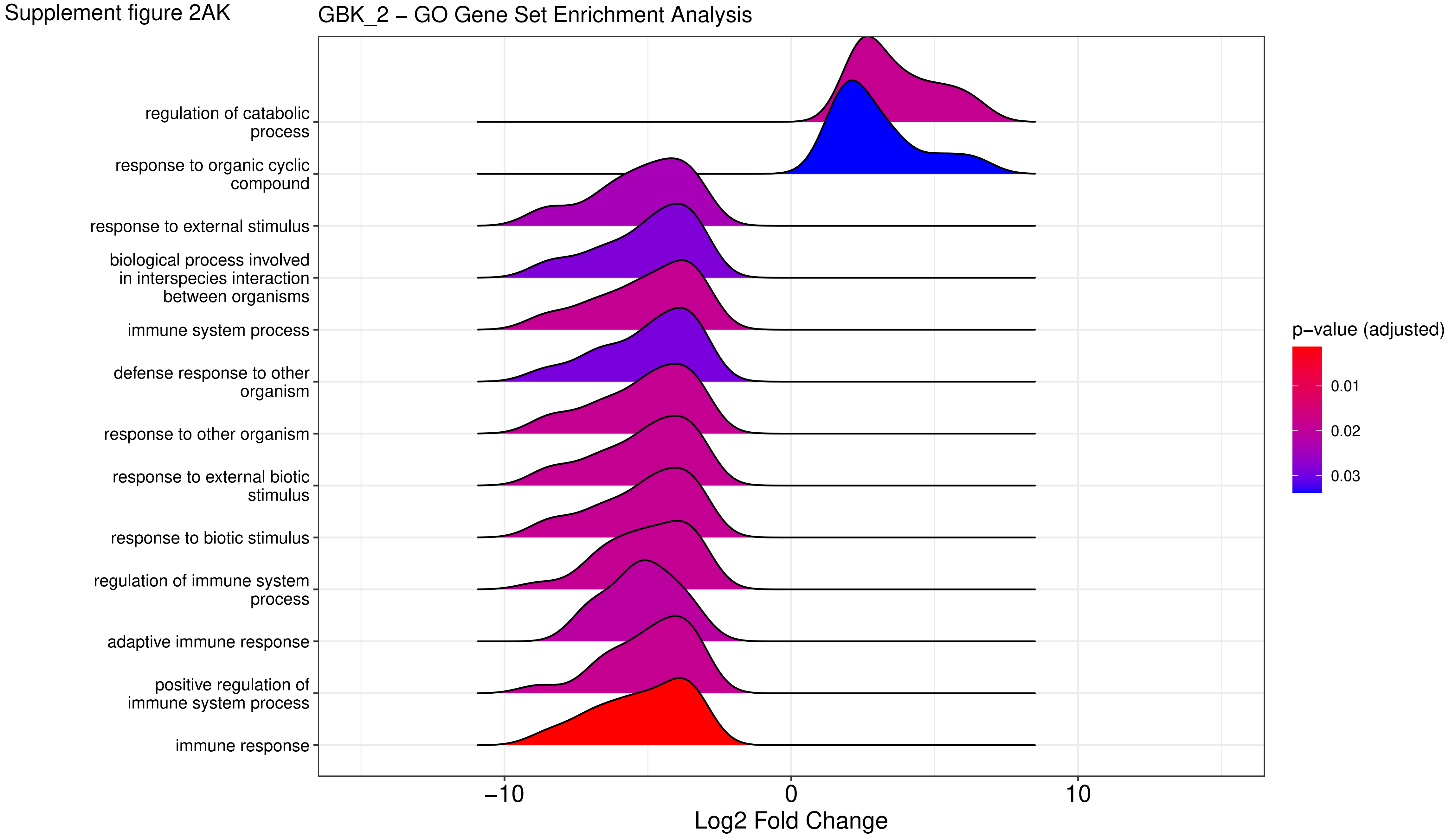

Supplement figure 2AL

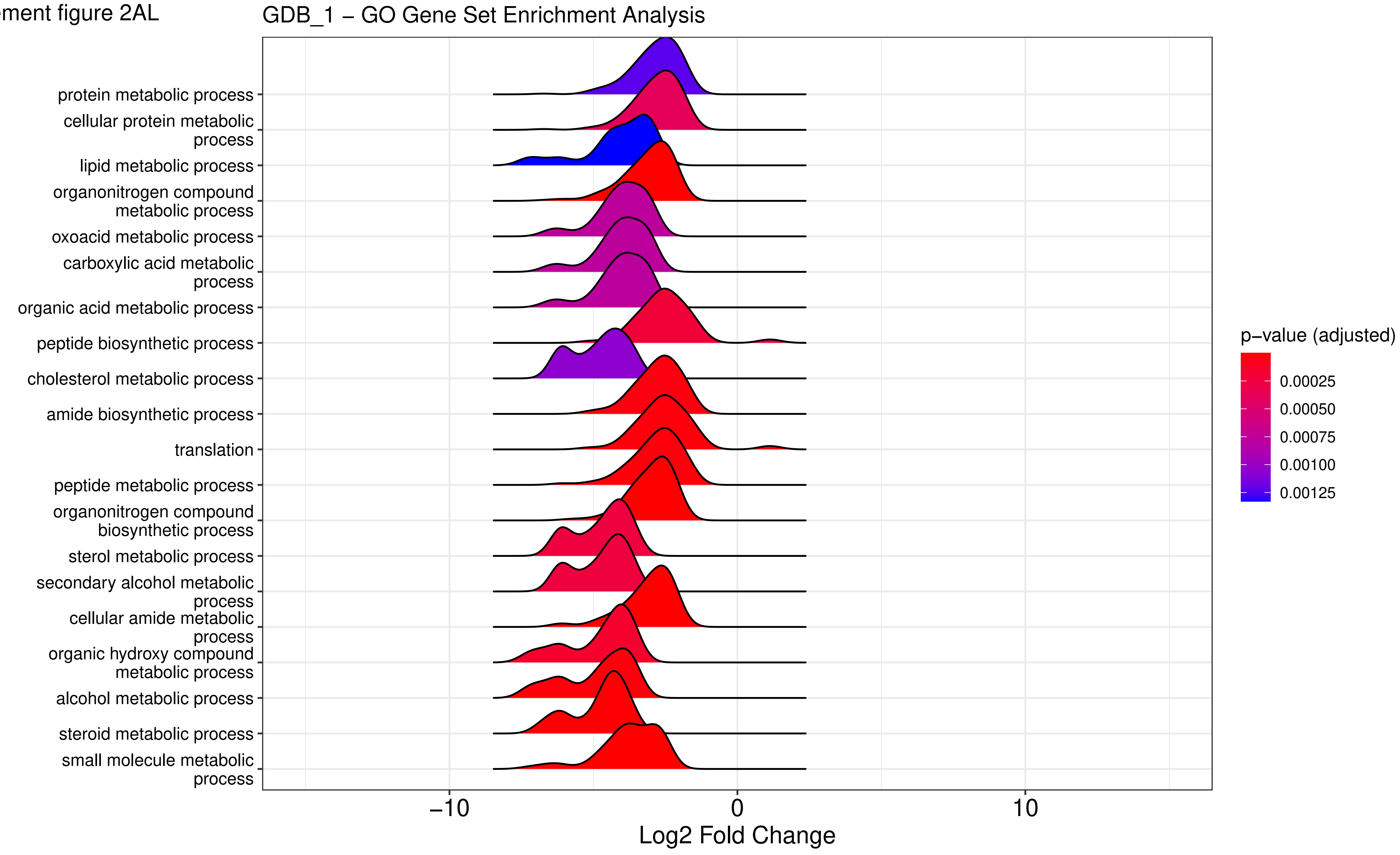

Supplement figure 2AM

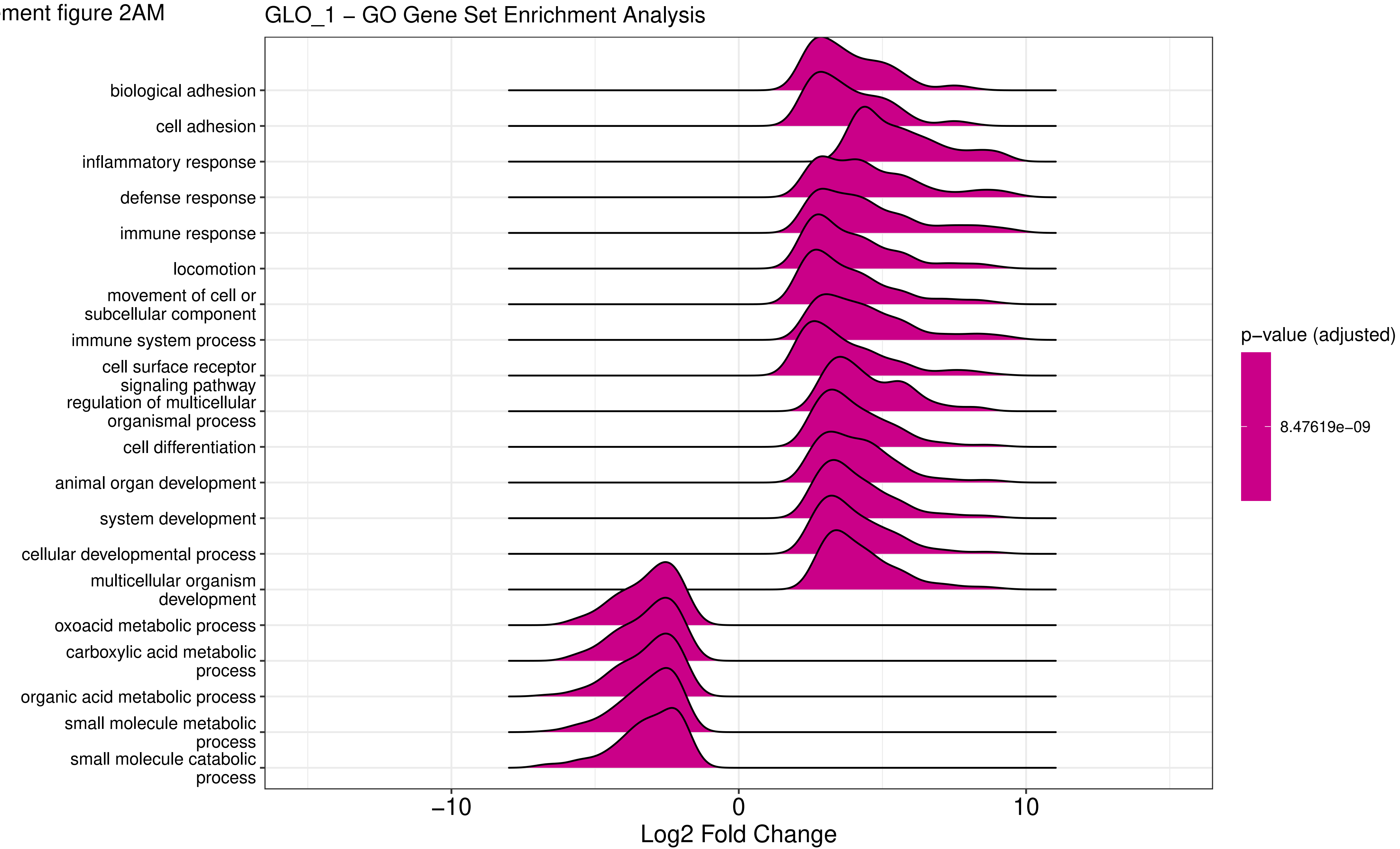

Supplement figure 2AN

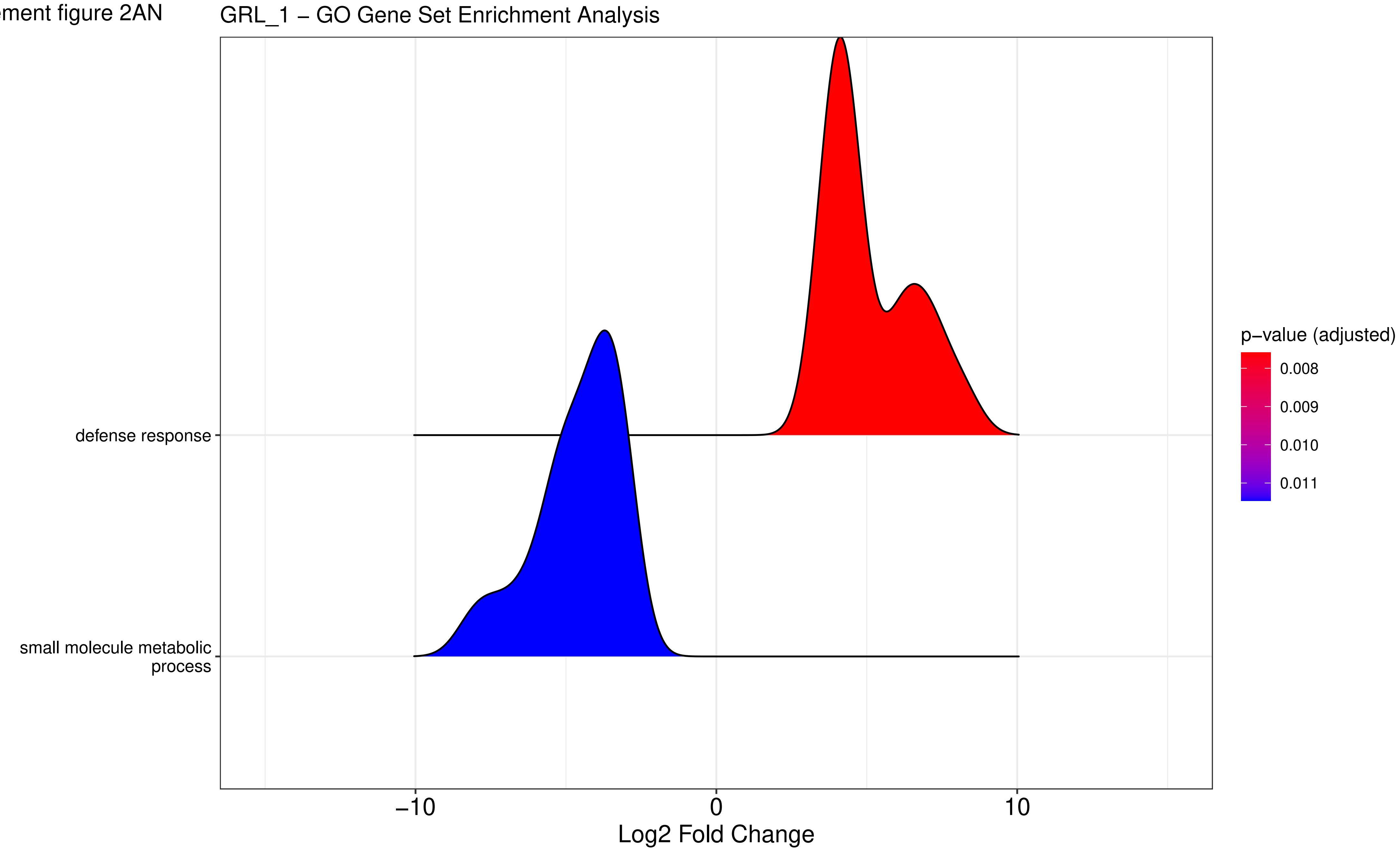

Supplement figure 2AO

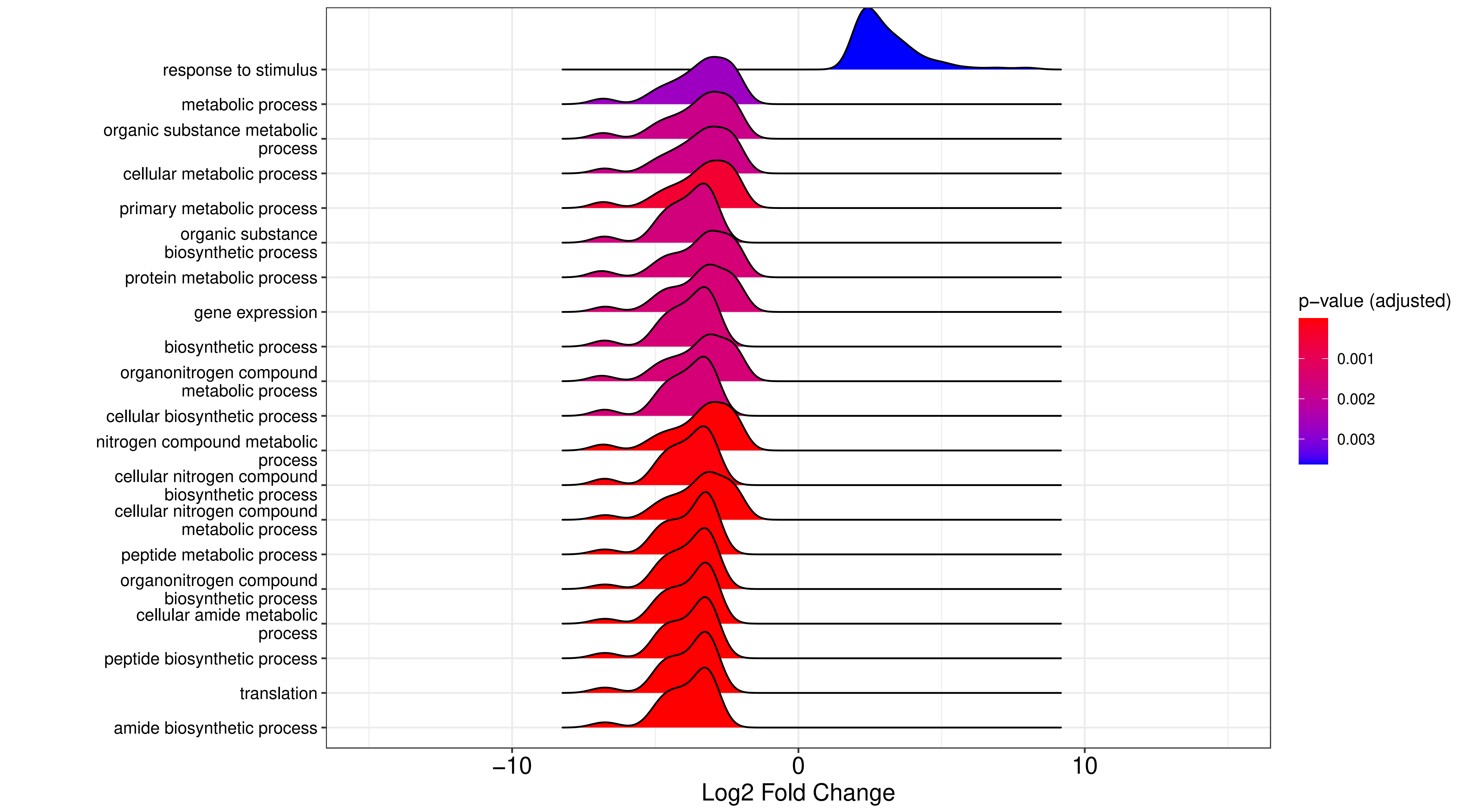

Supplement figure 2AP

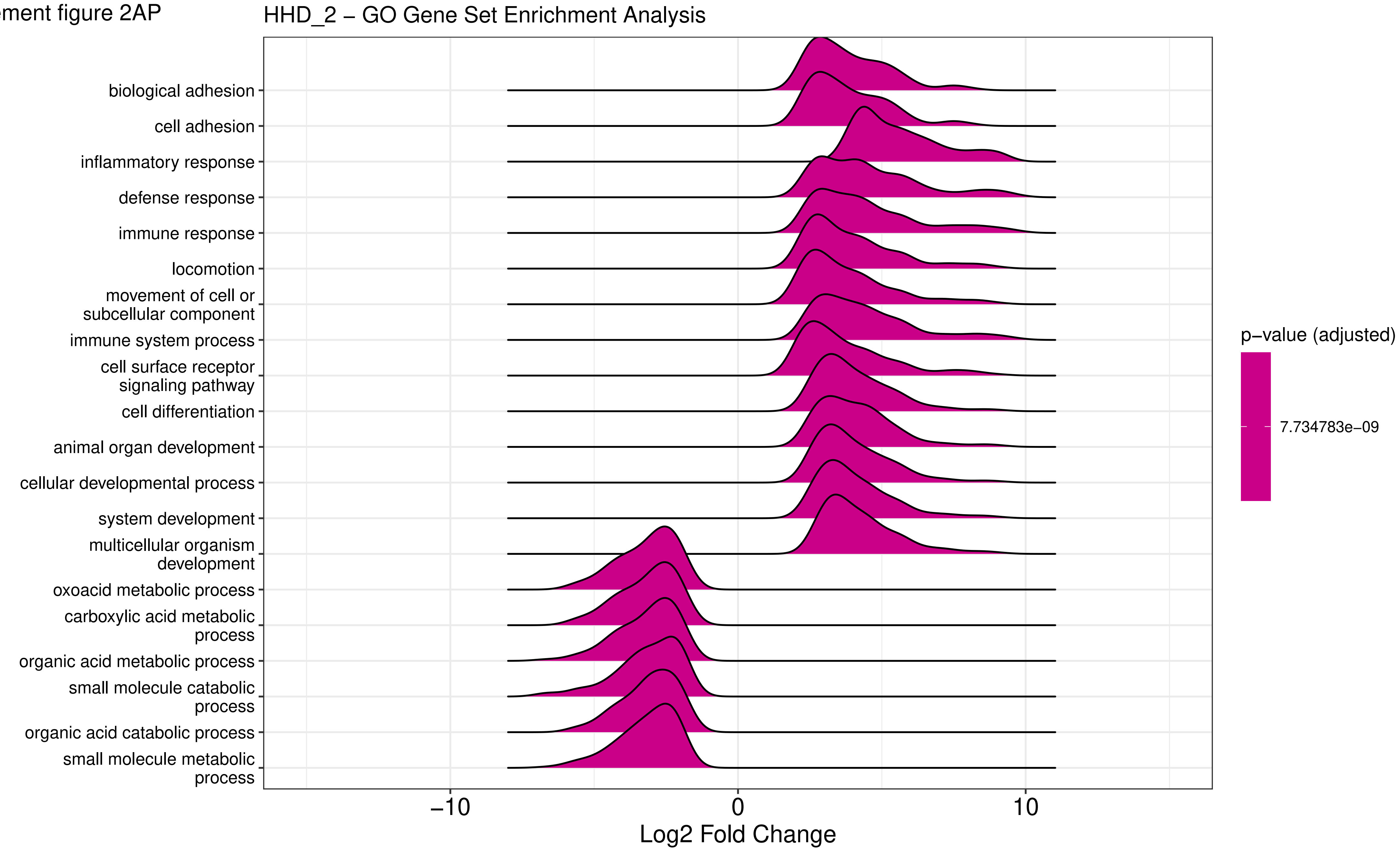

Supplement figure 2AQ

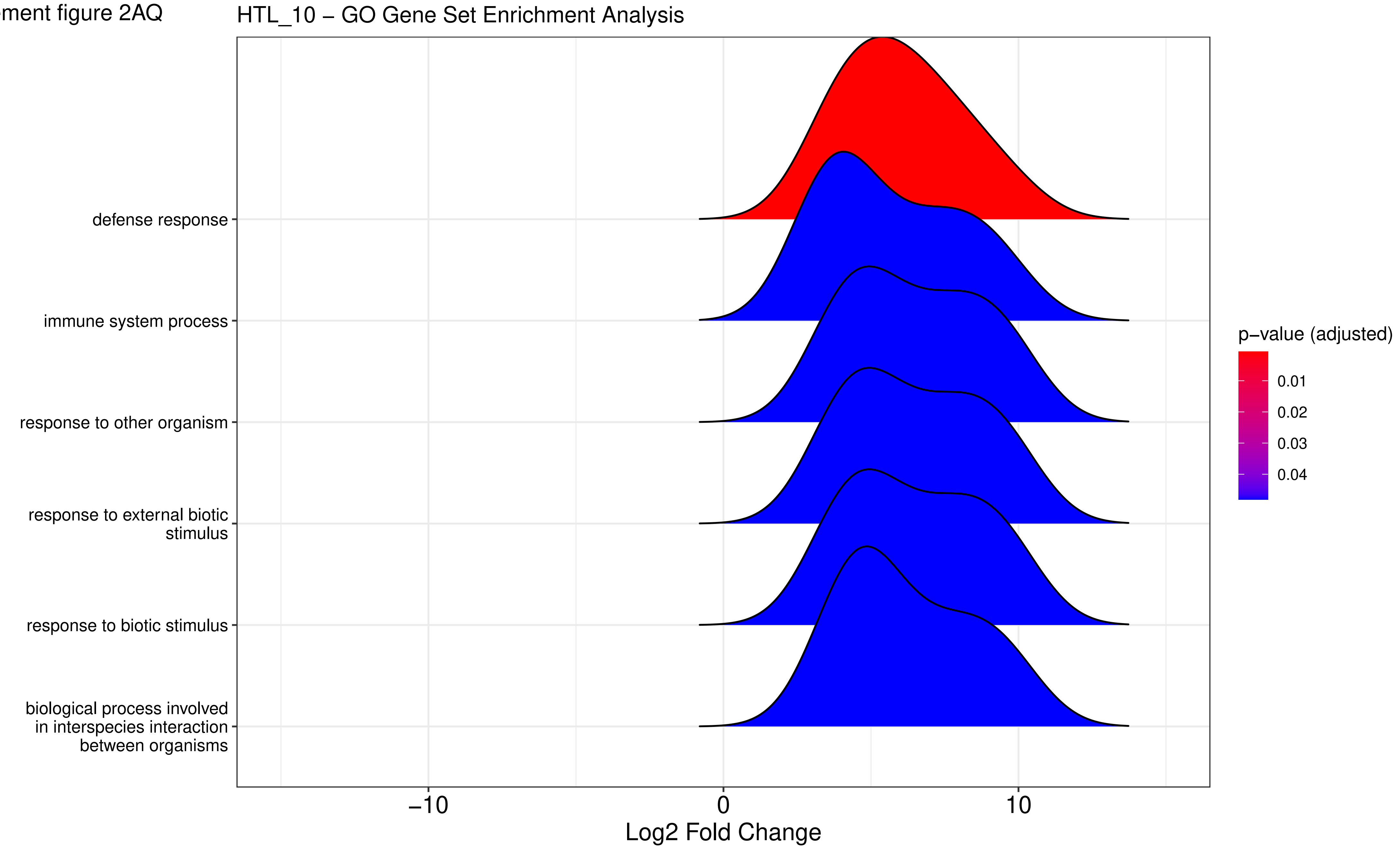

Supplement figure 2AR

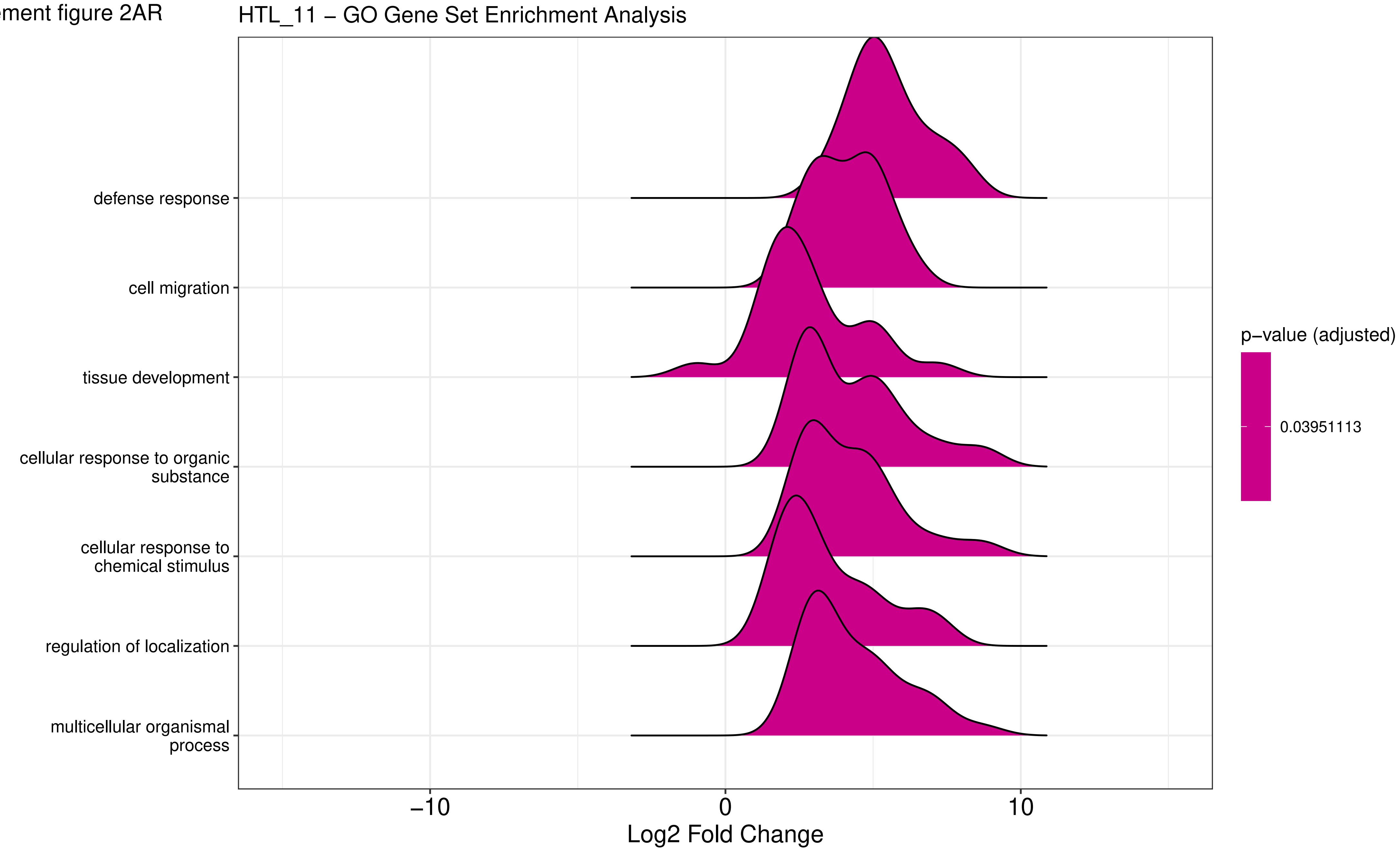

Supplement figure 2AS

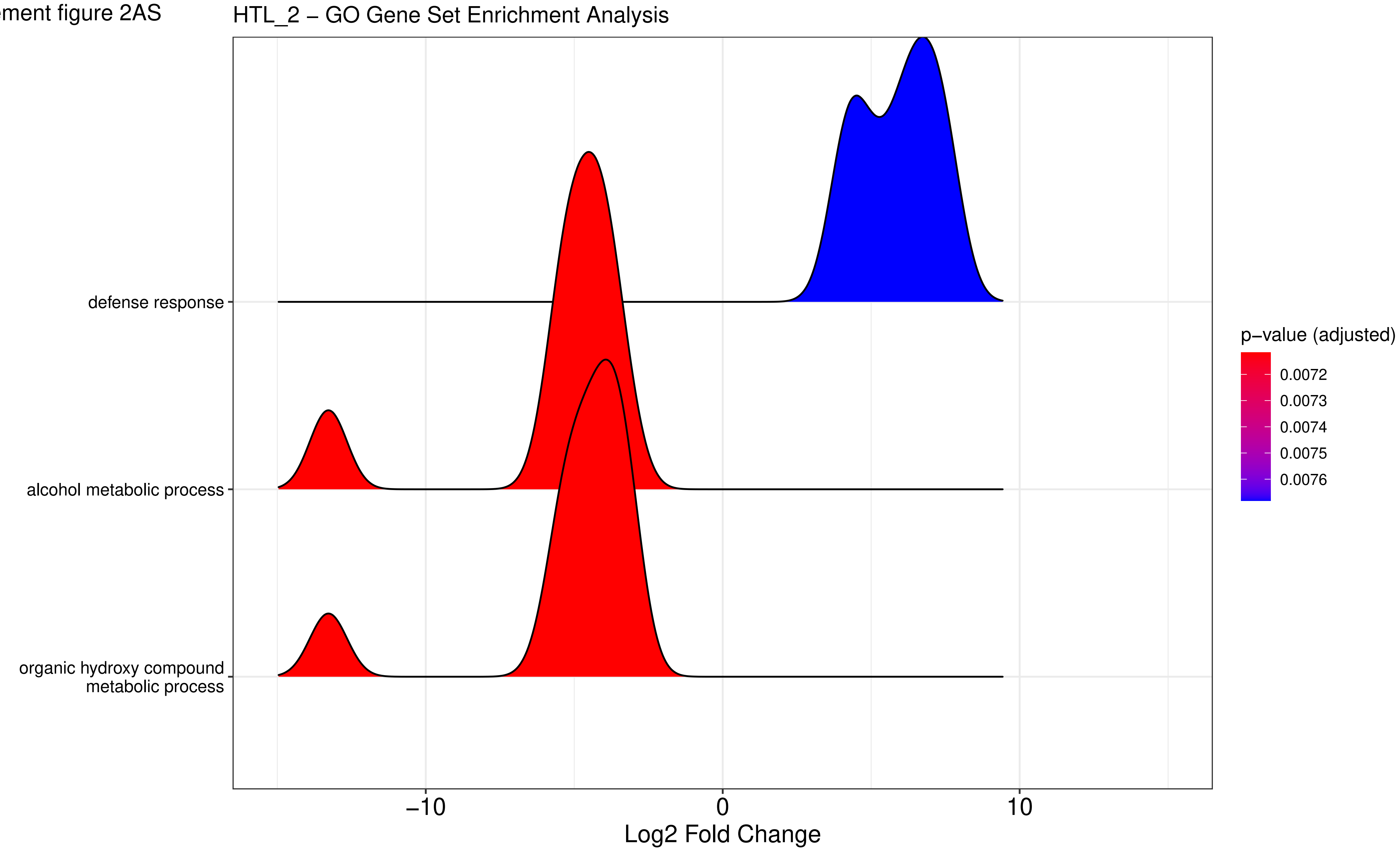

Supplement figure 2AT

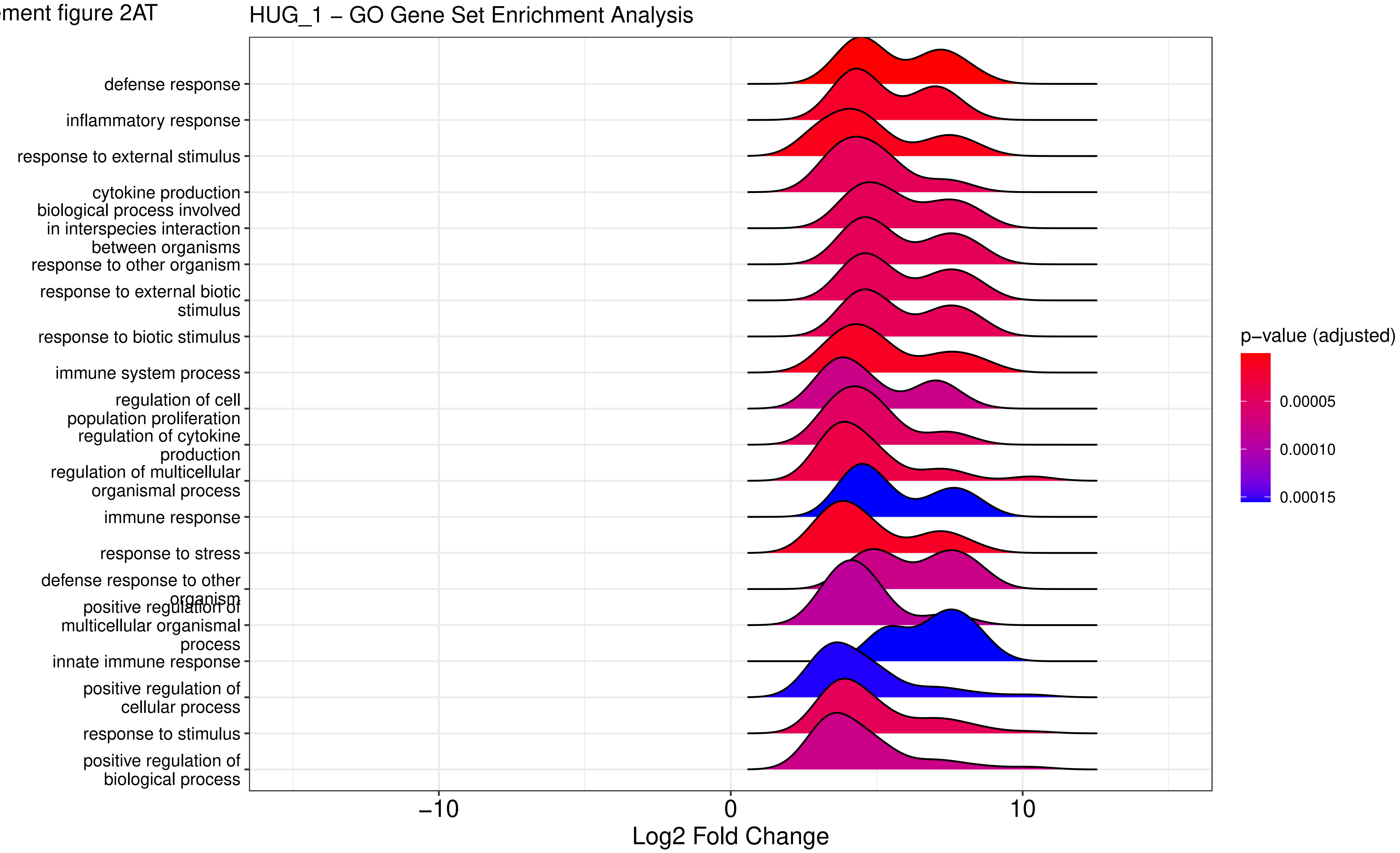

Supplement figure 2AU

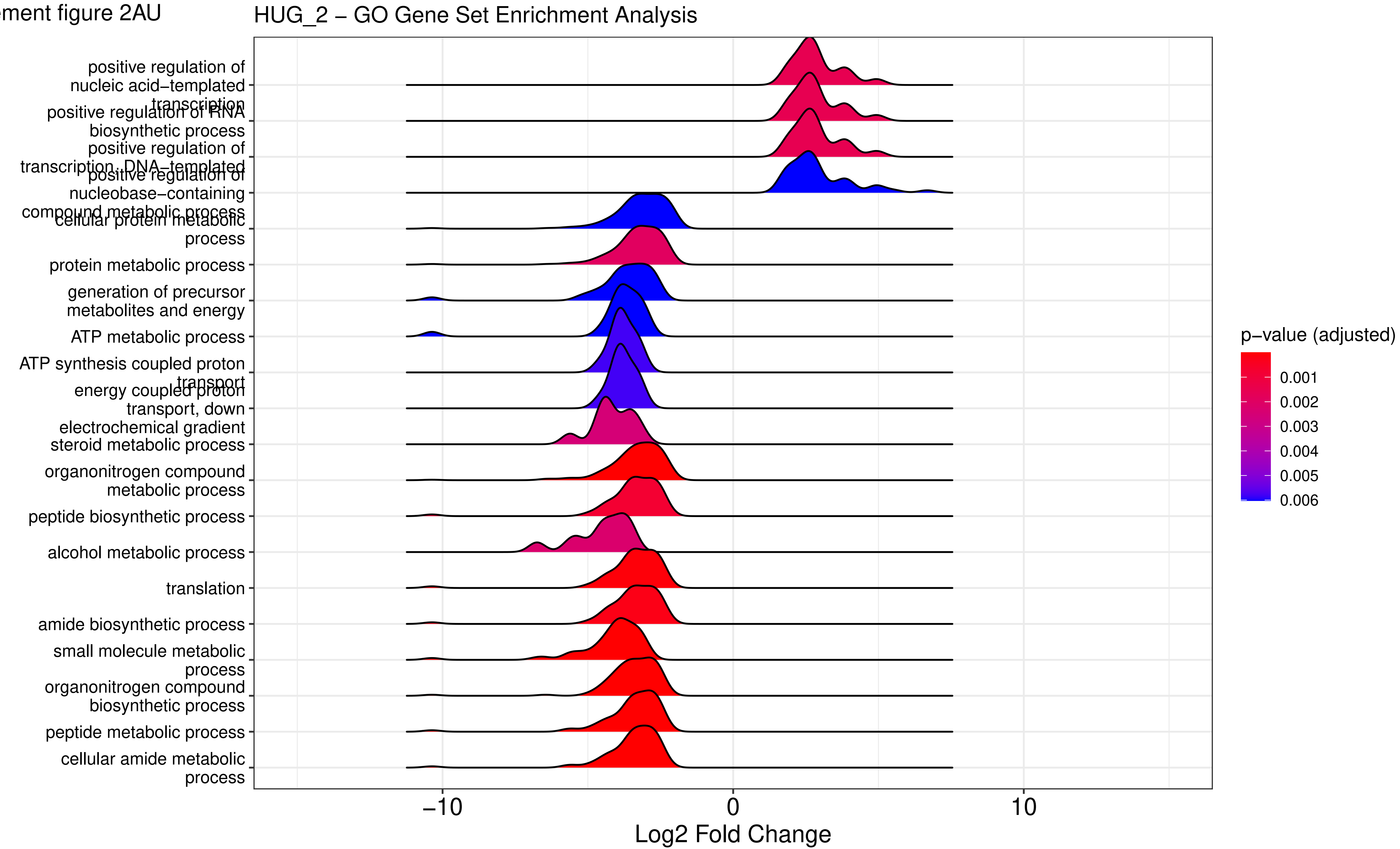

Supplement figure 2AV

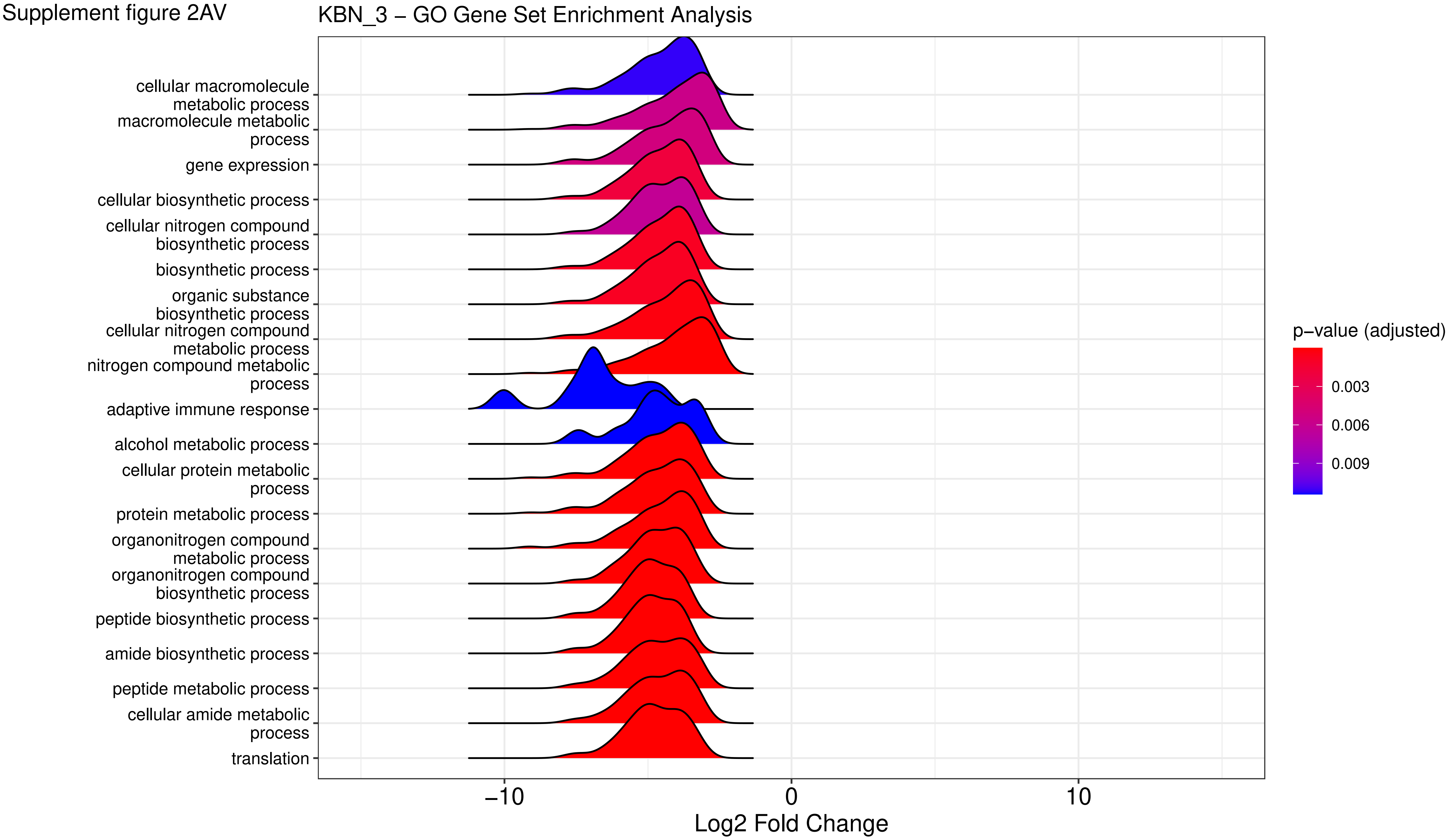

Supplement figure 2AW

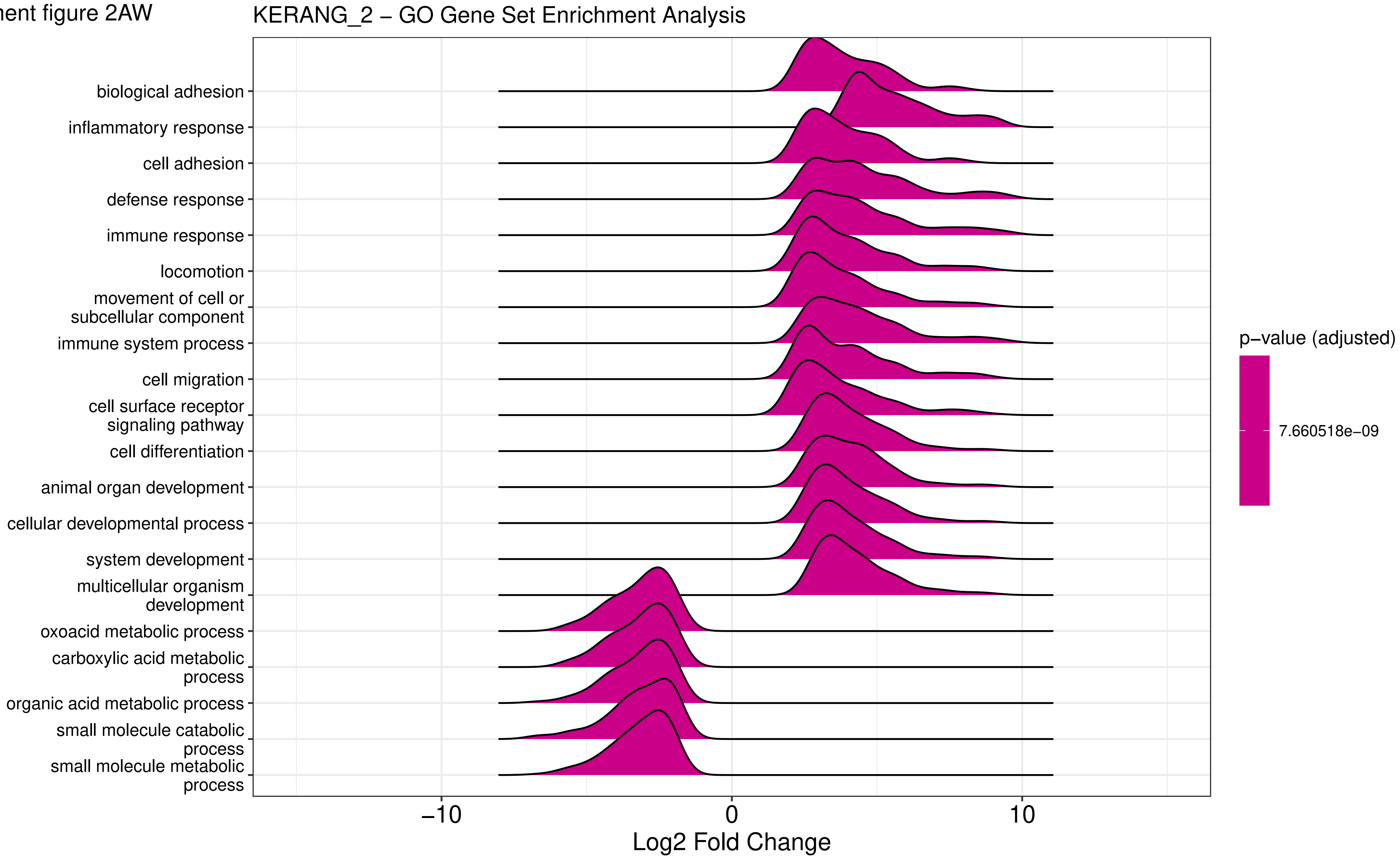

Supplement figure 2AX

KGS_5 – GO Gene Set Enrichment Analysis

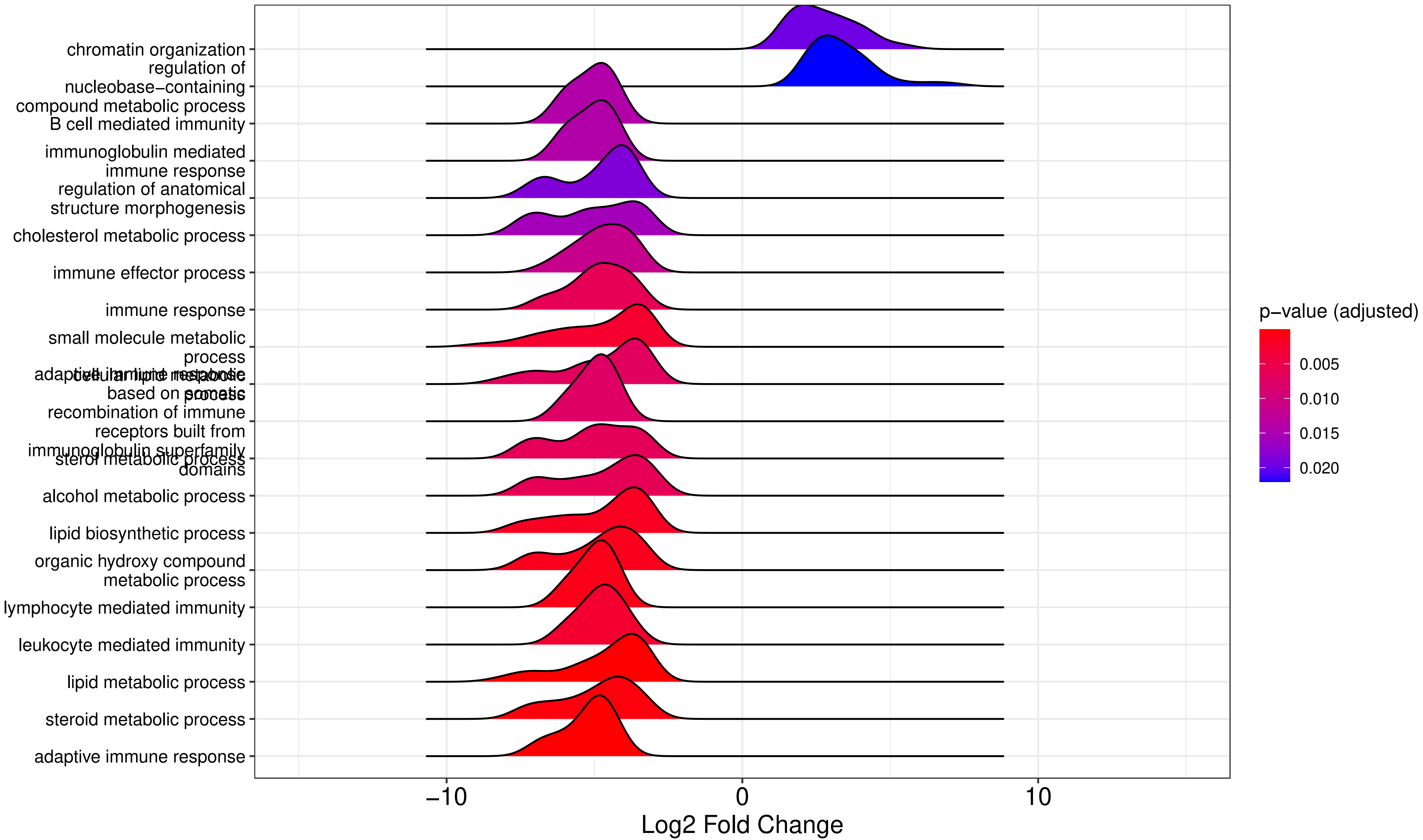

Supplement figure 2AY

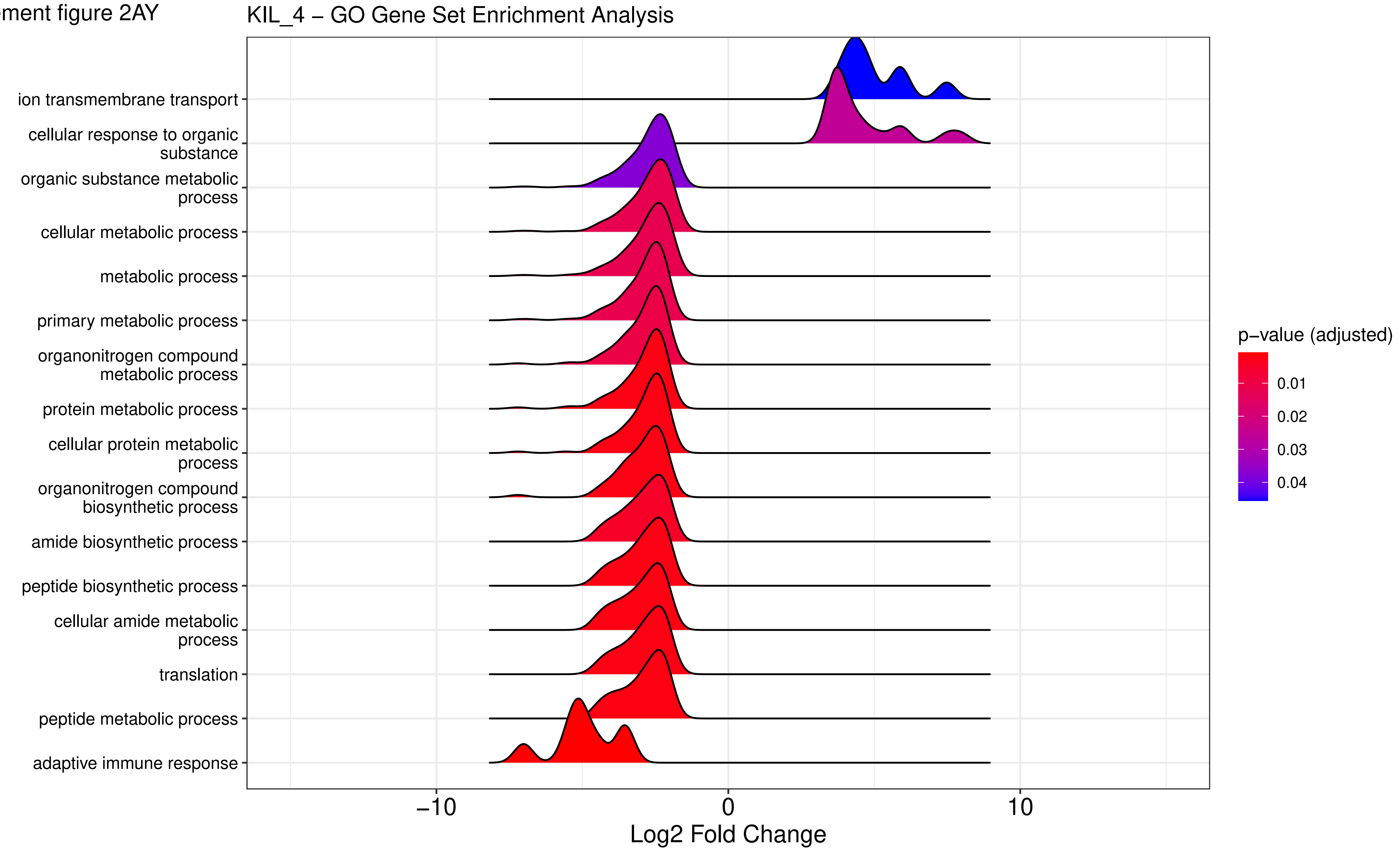

Supplement figure 2AZ

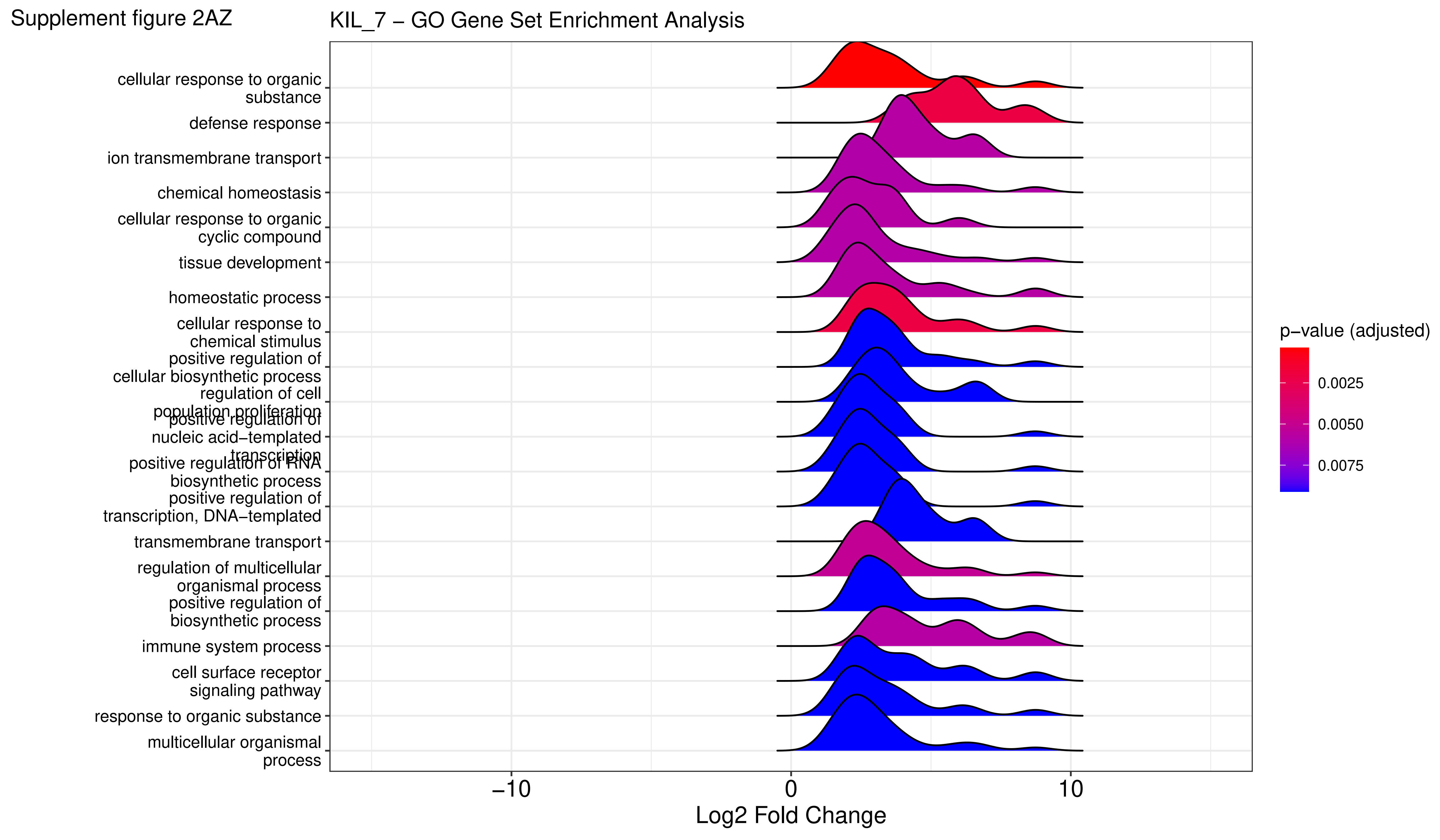

Supplement figure 2BA

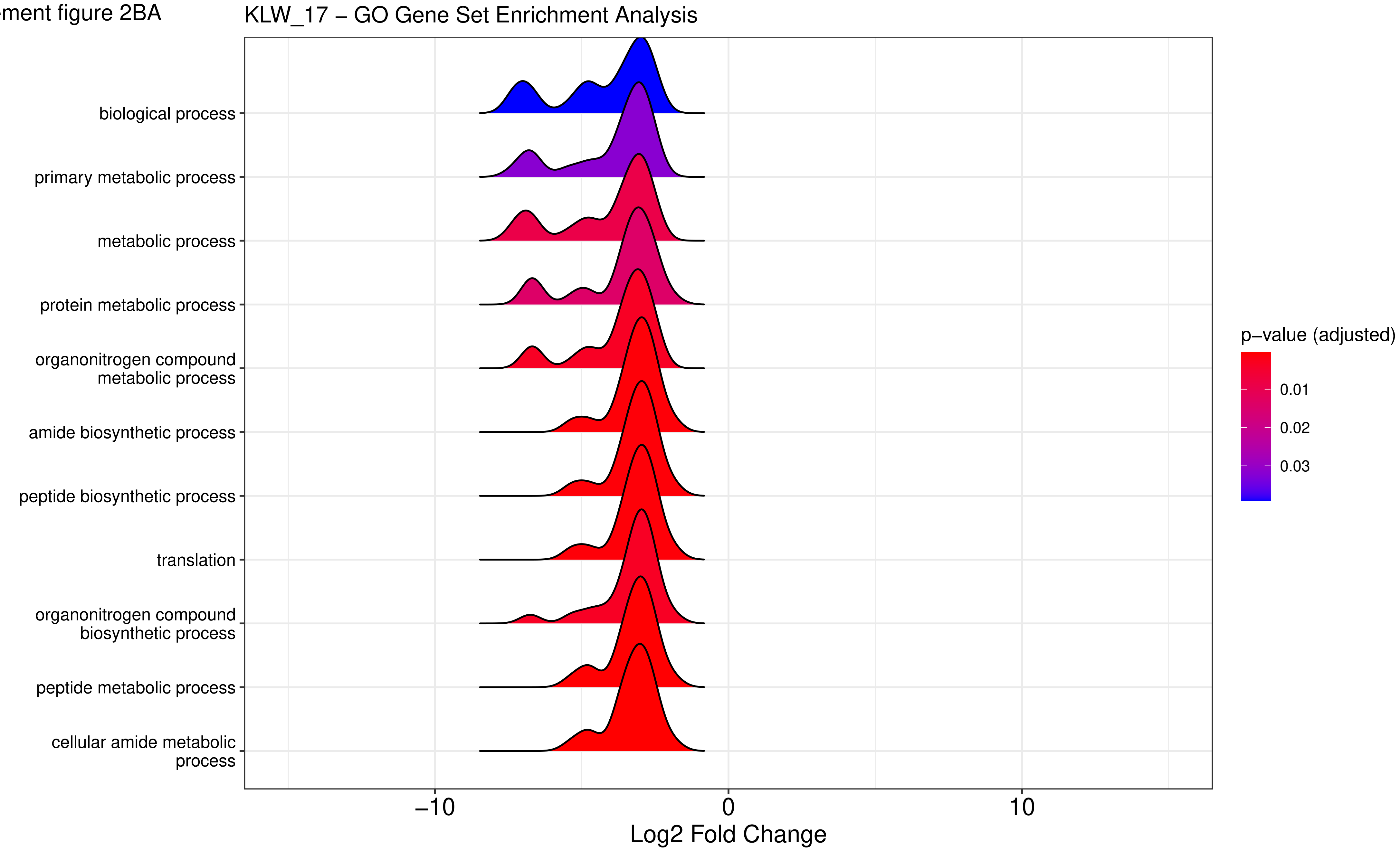

Supplement figure 2BB

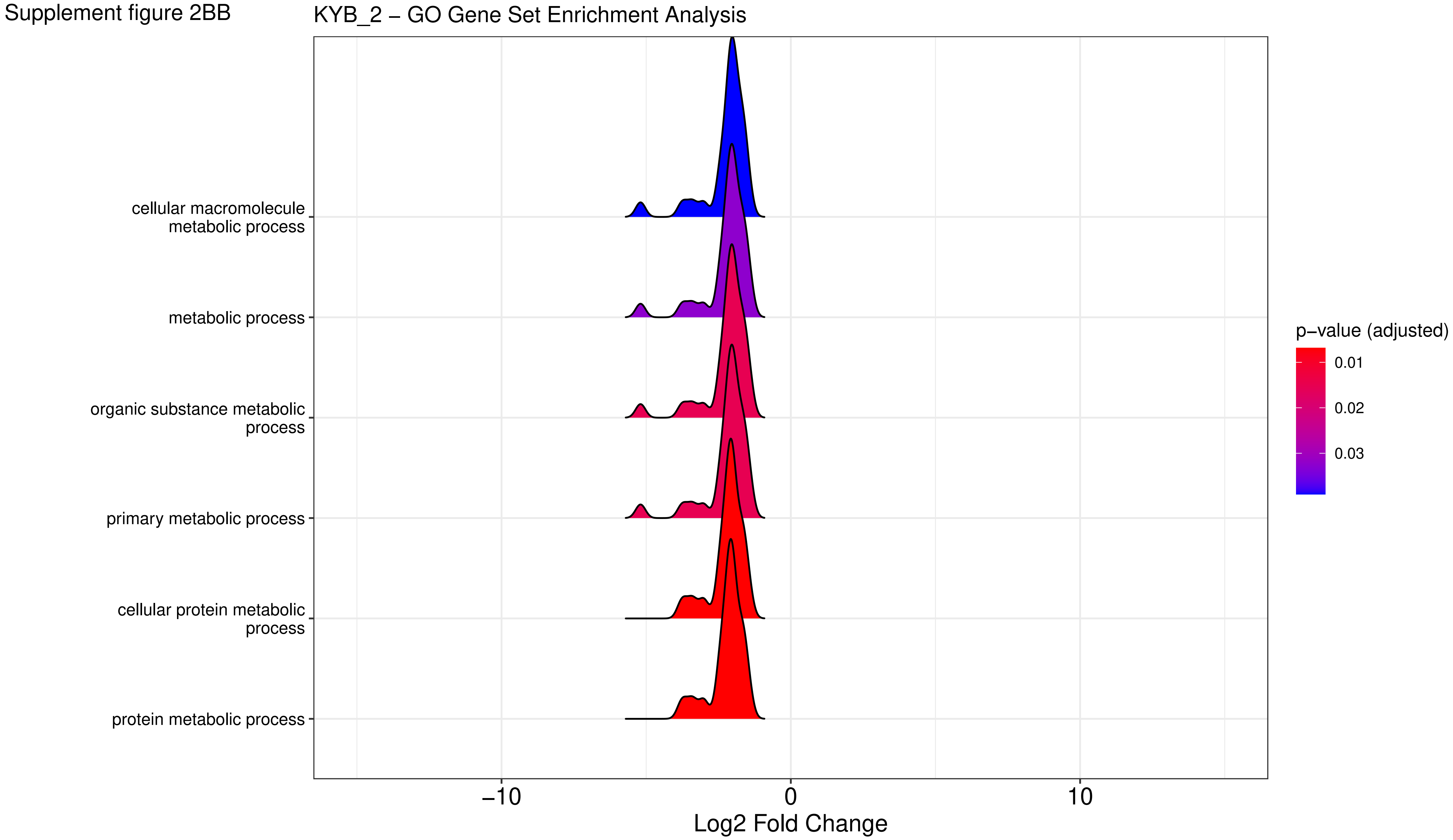

Supplement figure 2BC

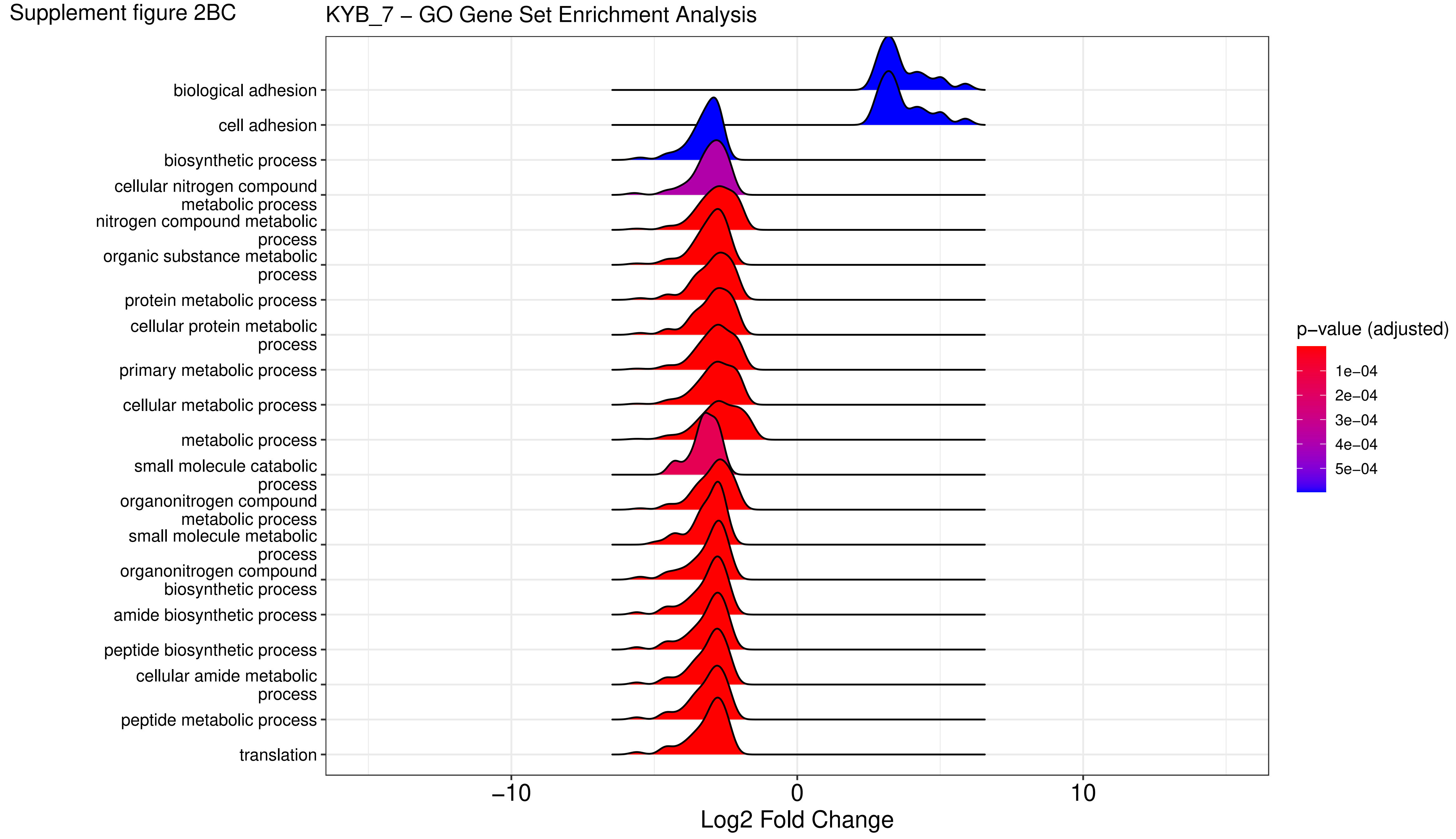

Supplement figure 2BD

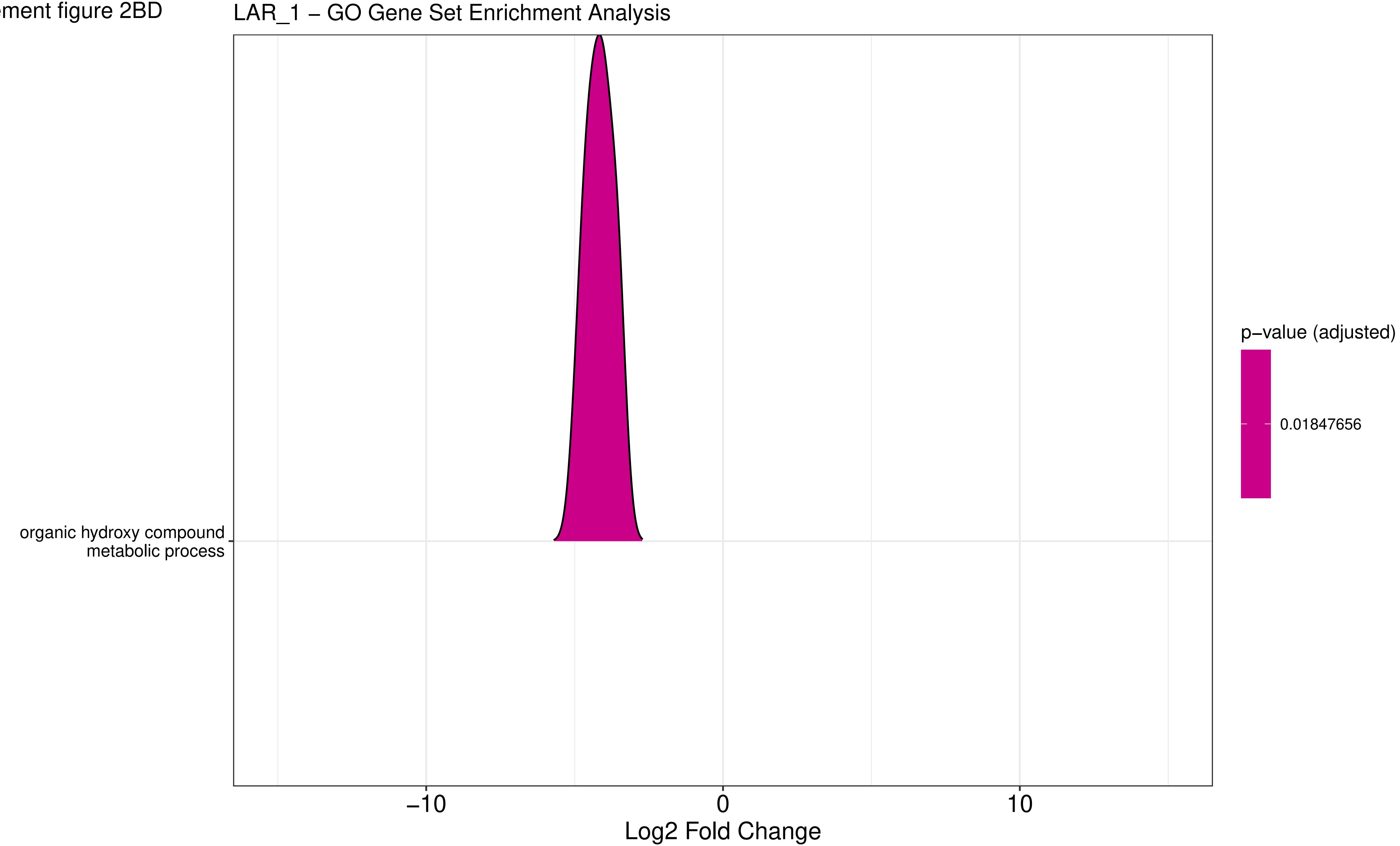

Supplement figure 2BE

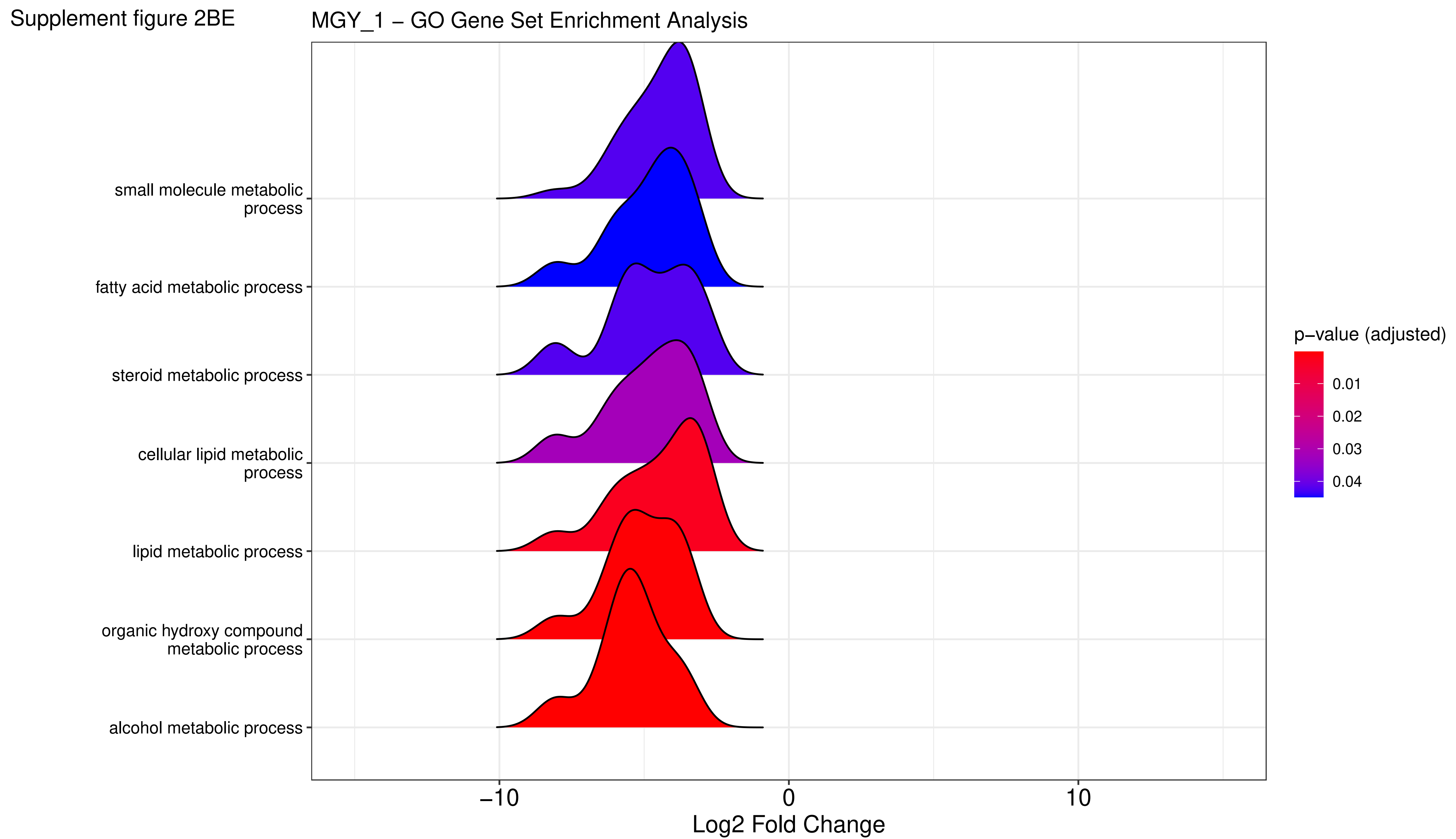

Supplement figure 2BF

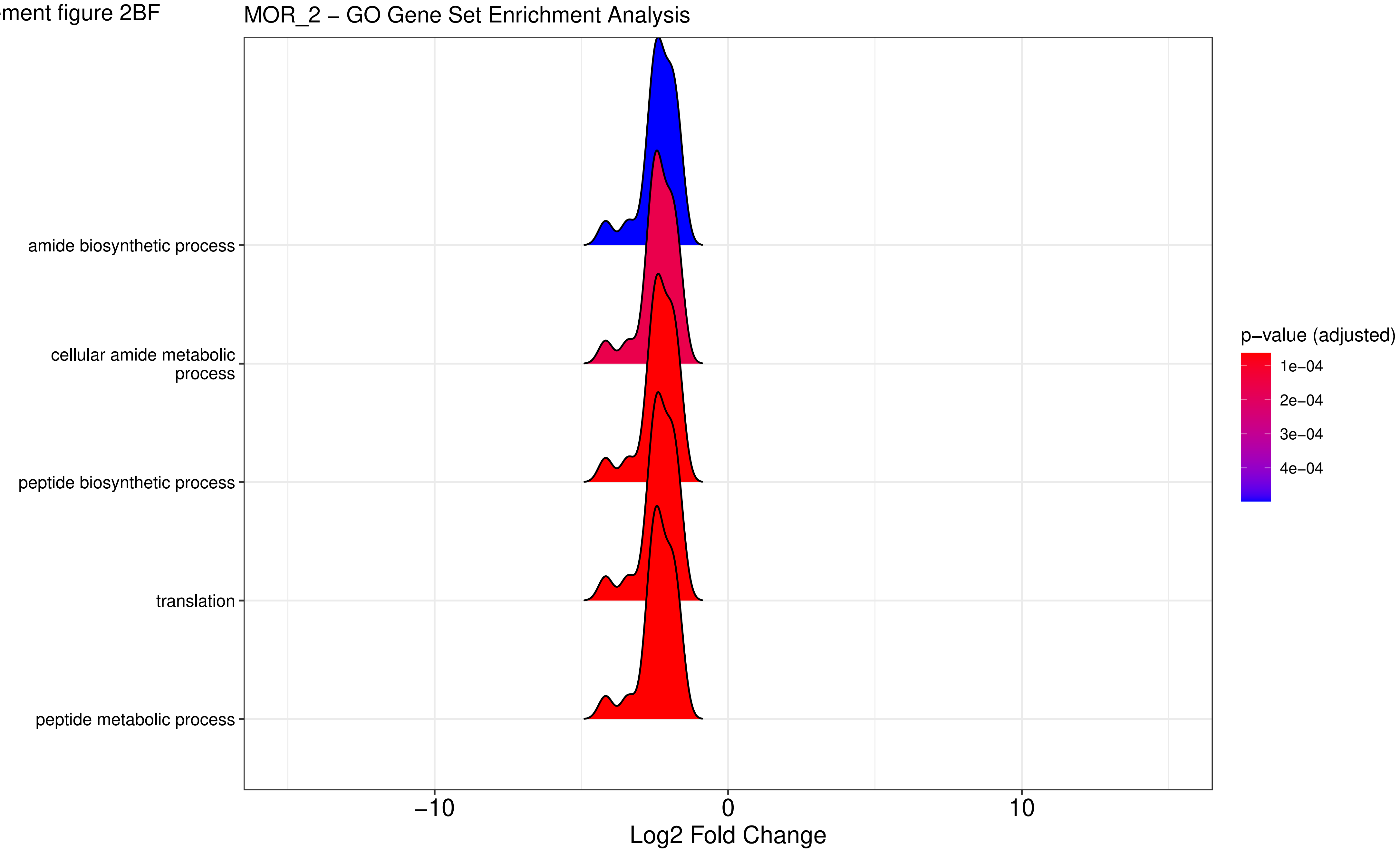

Supplement figure 2BG

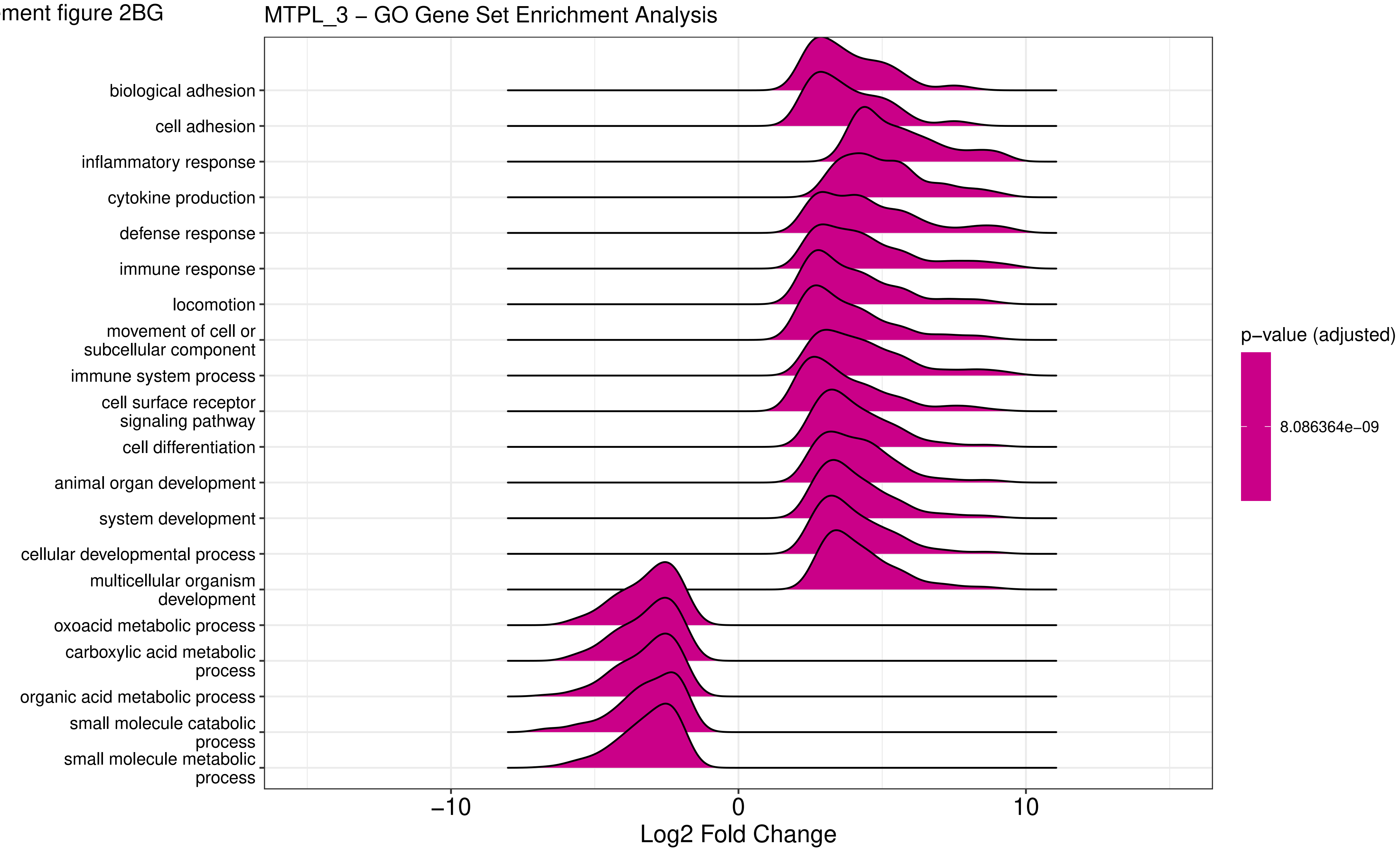

Supplement figure 2BH

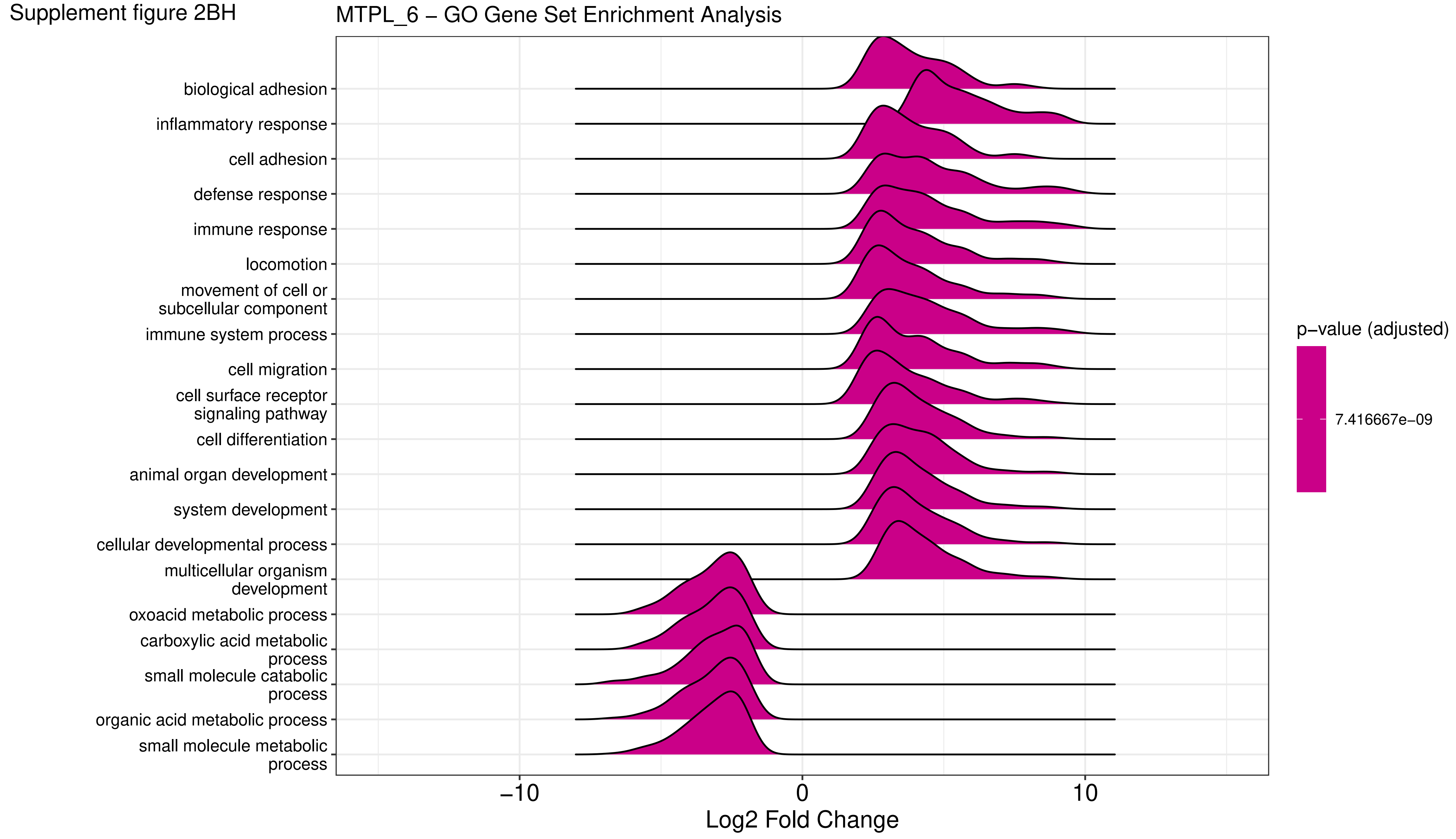

Supplement figure 2BI

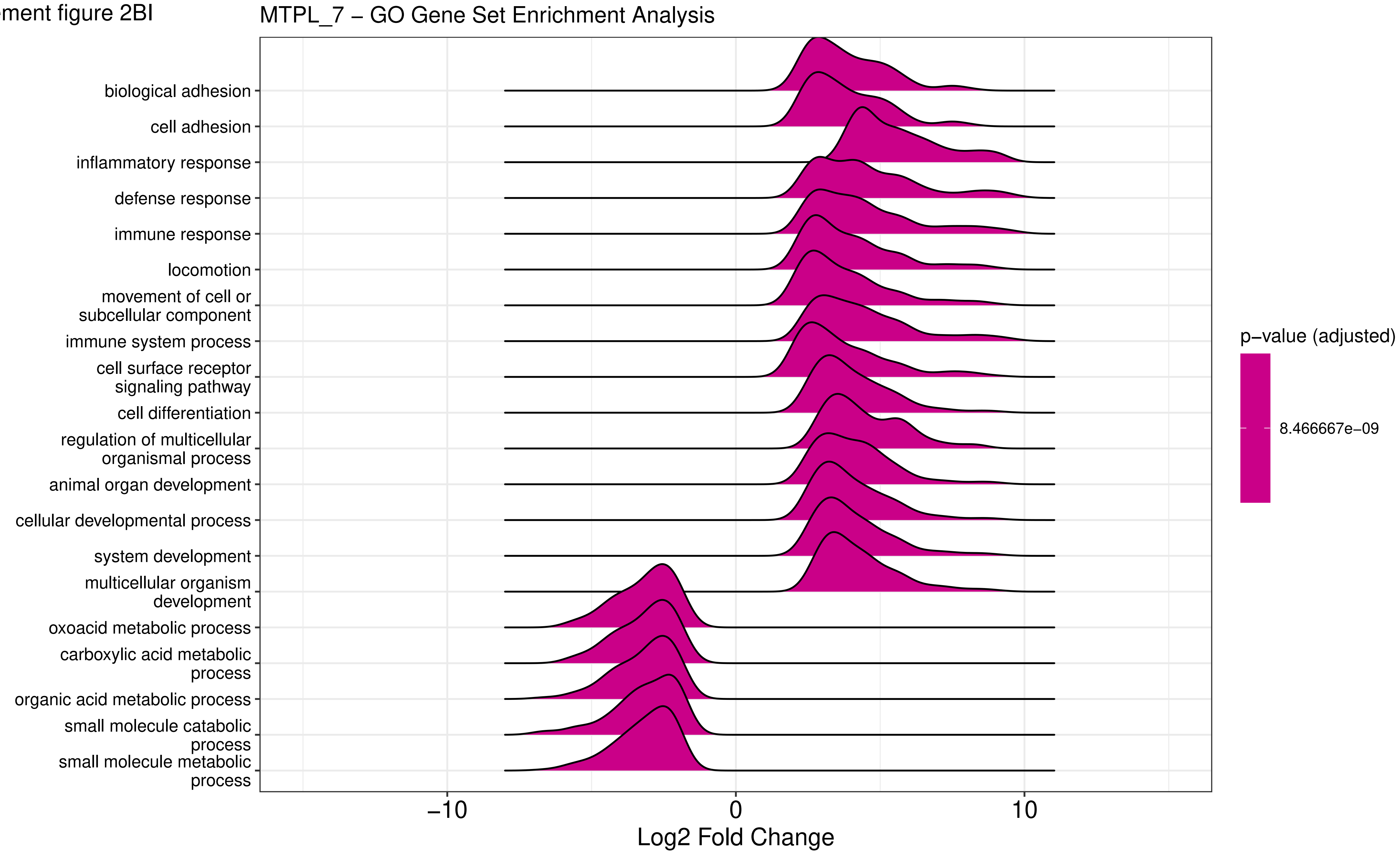

Supplement figure 2BJ

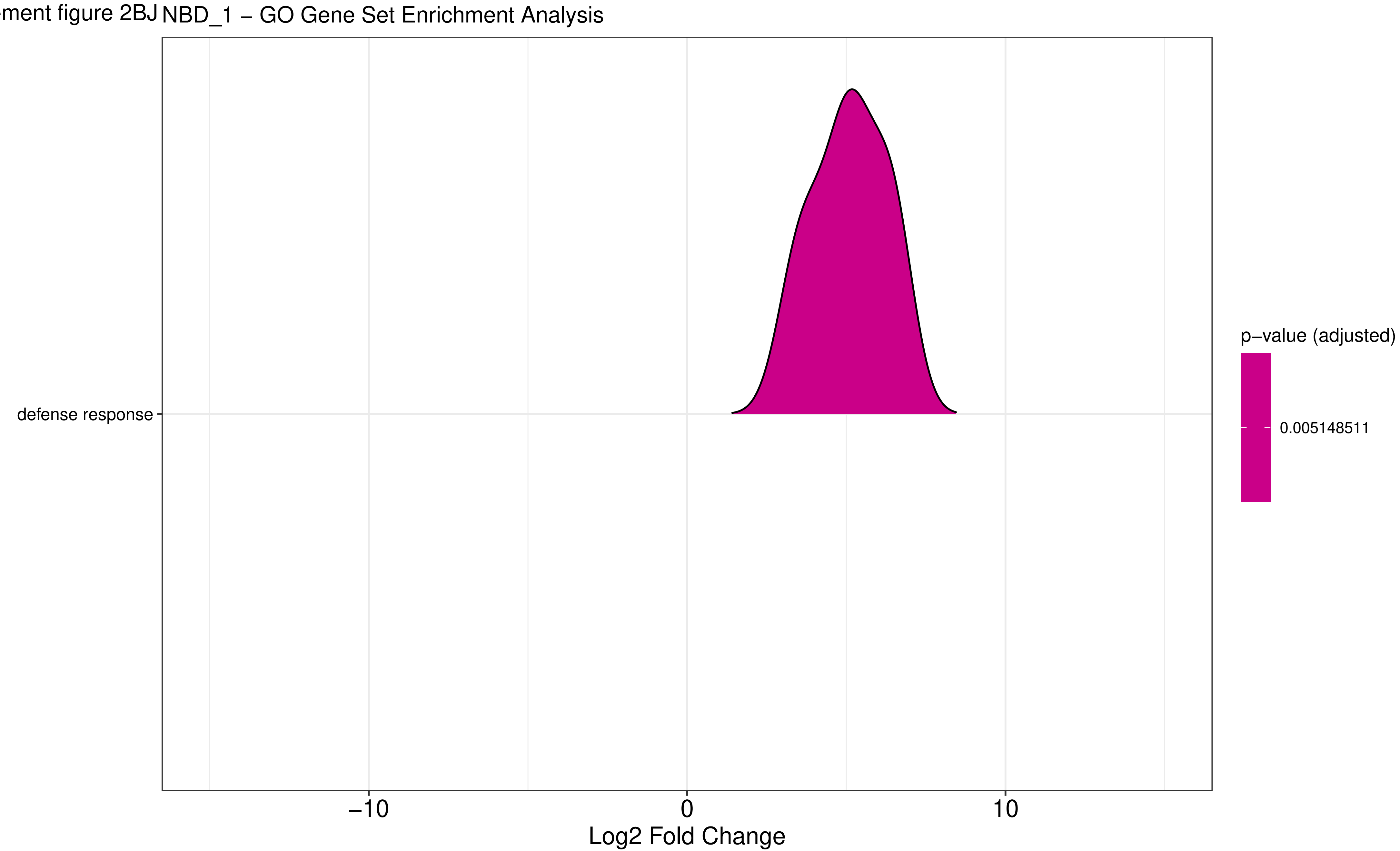

Supplement figure 2BK

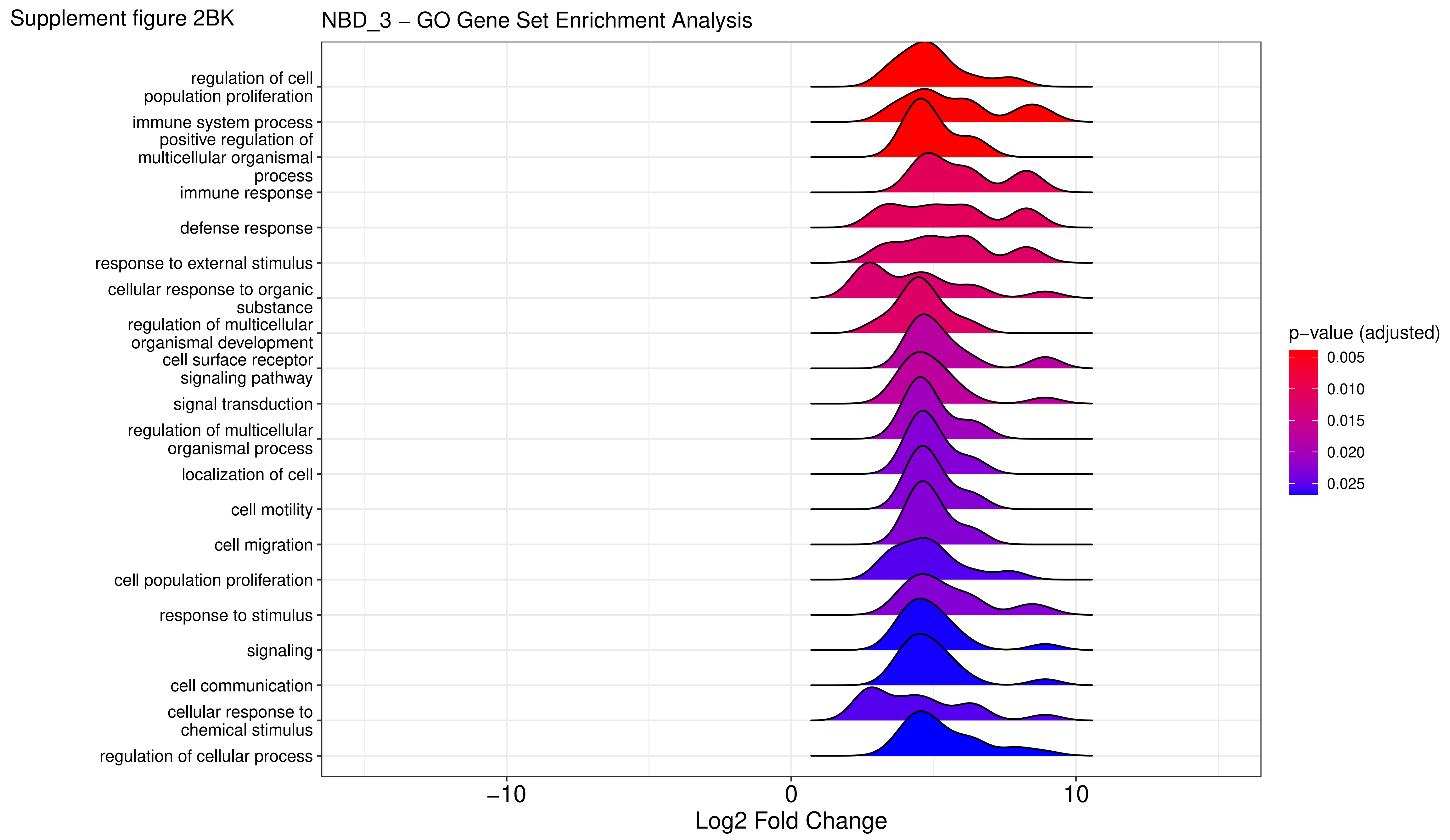

Supplement figure 2BL

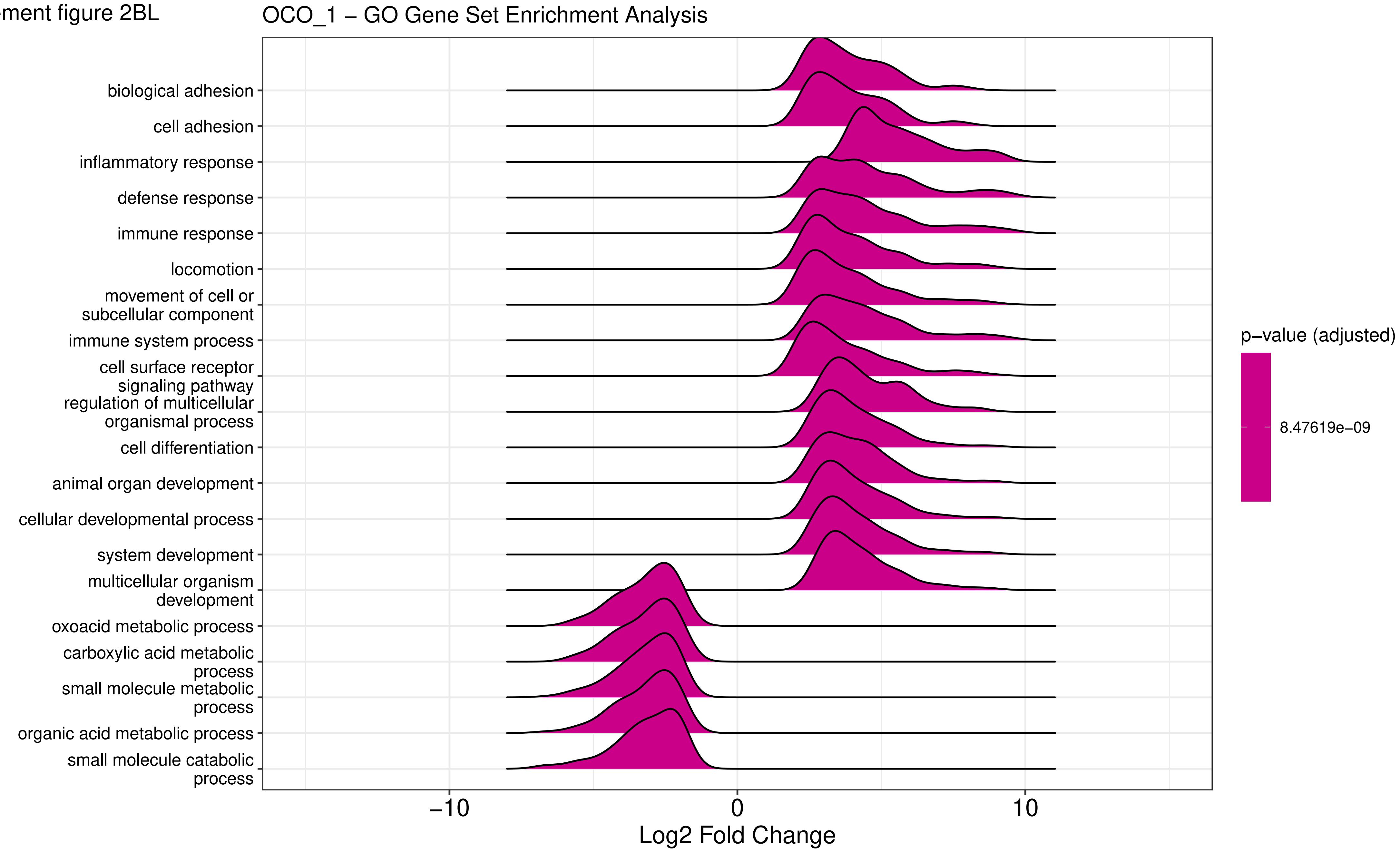

Supplement figure 2BM

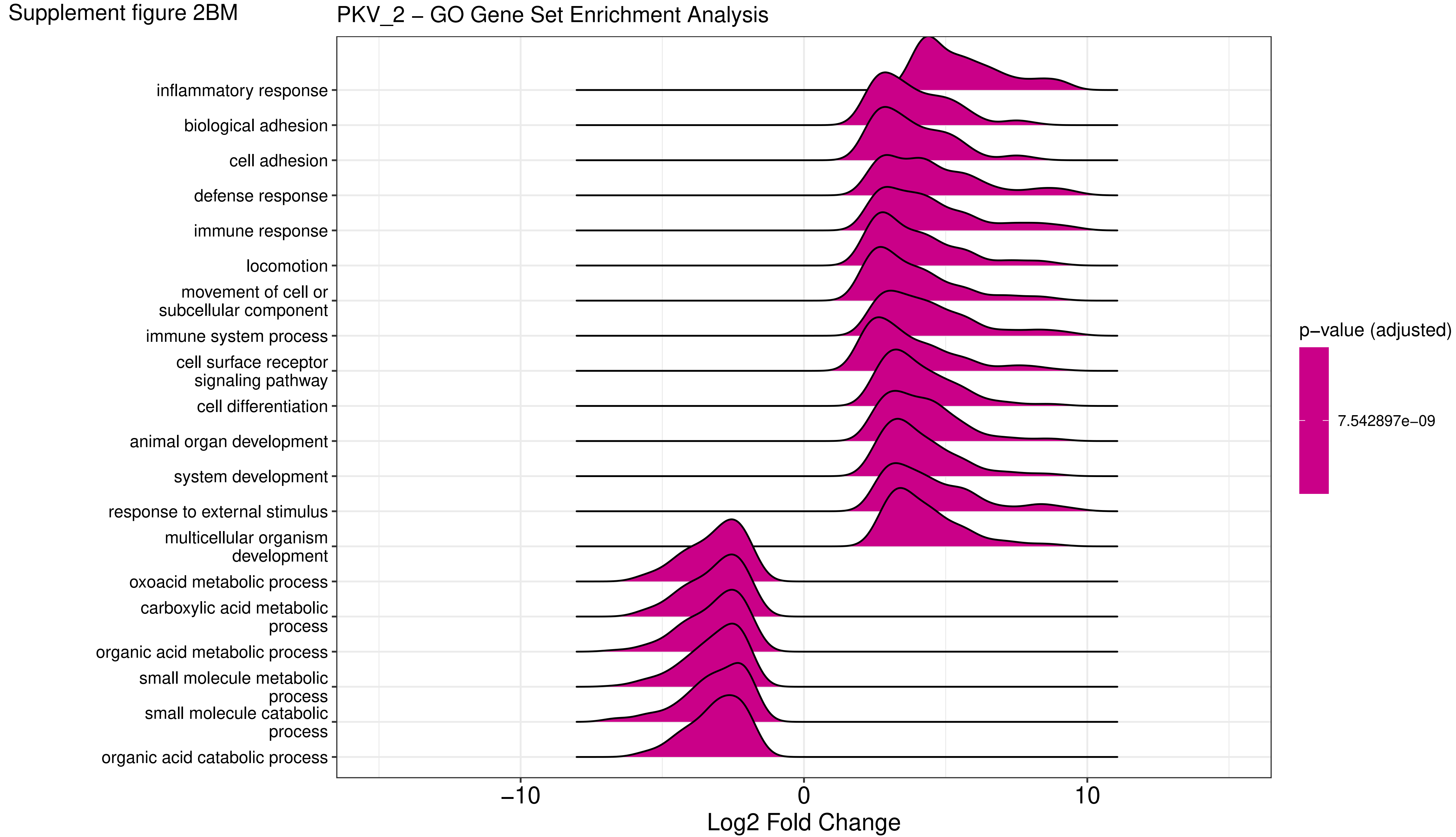

Supplement figure 2BN

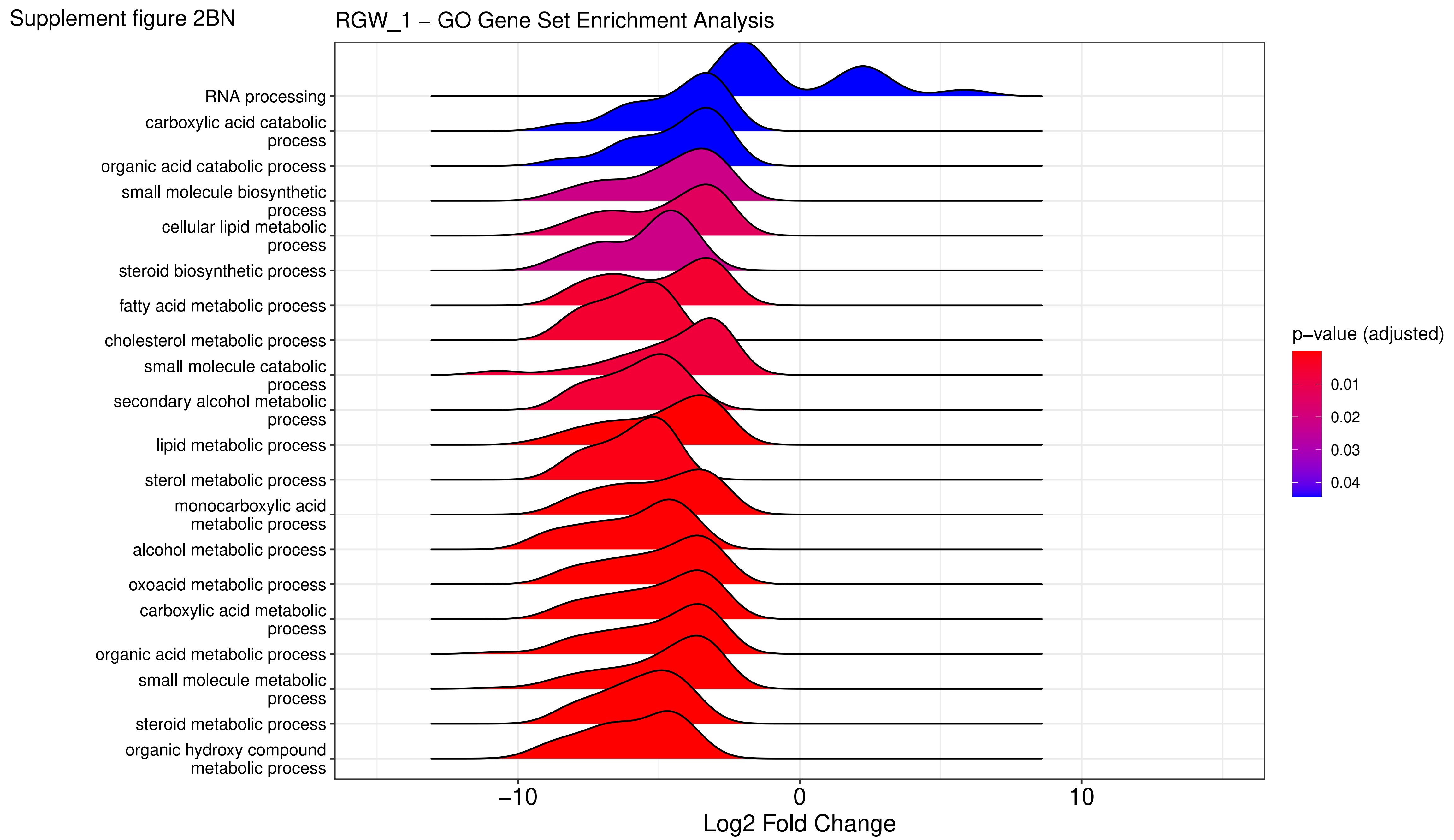

Supplement figure 2BO

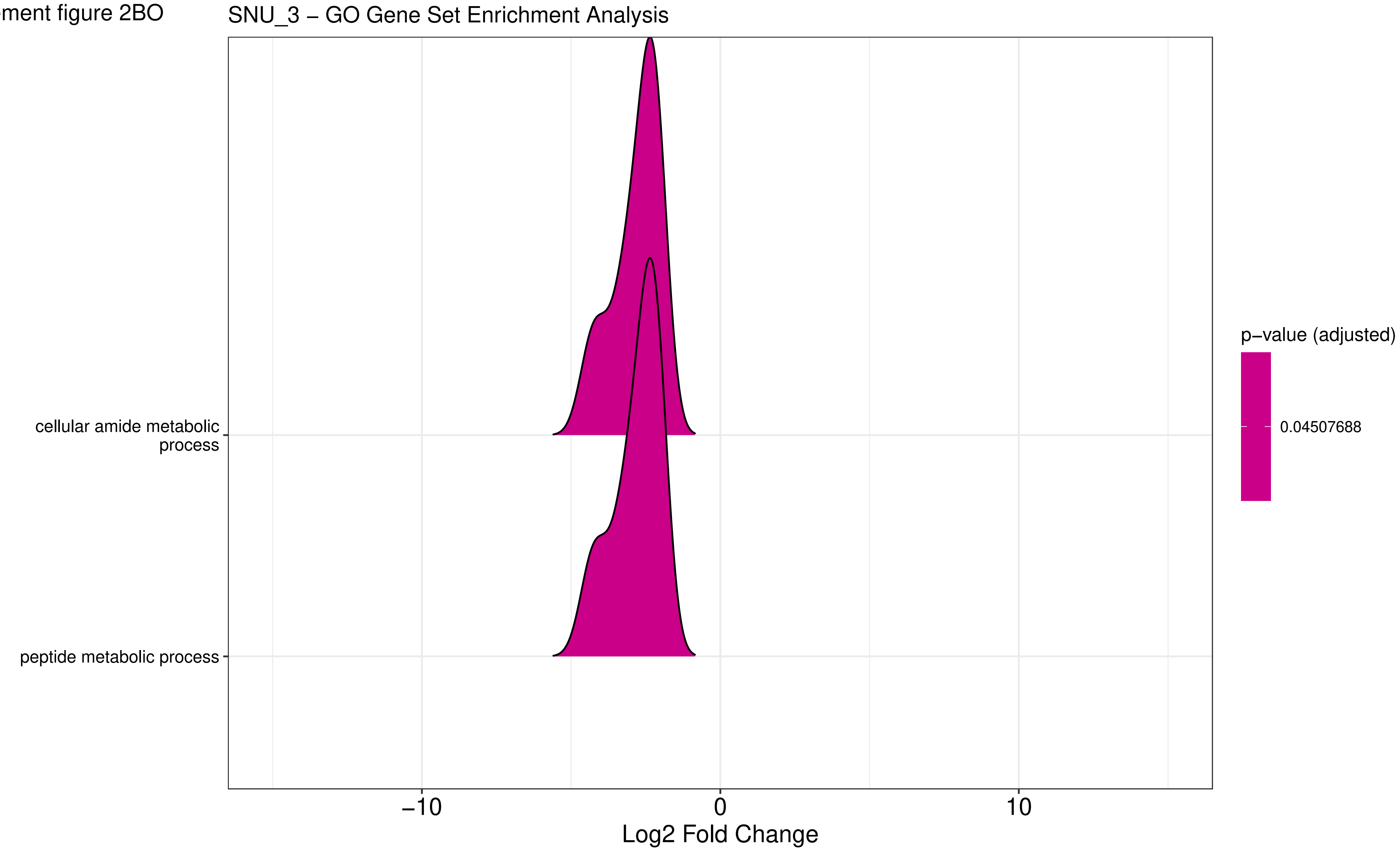

Supplement figure 2BP

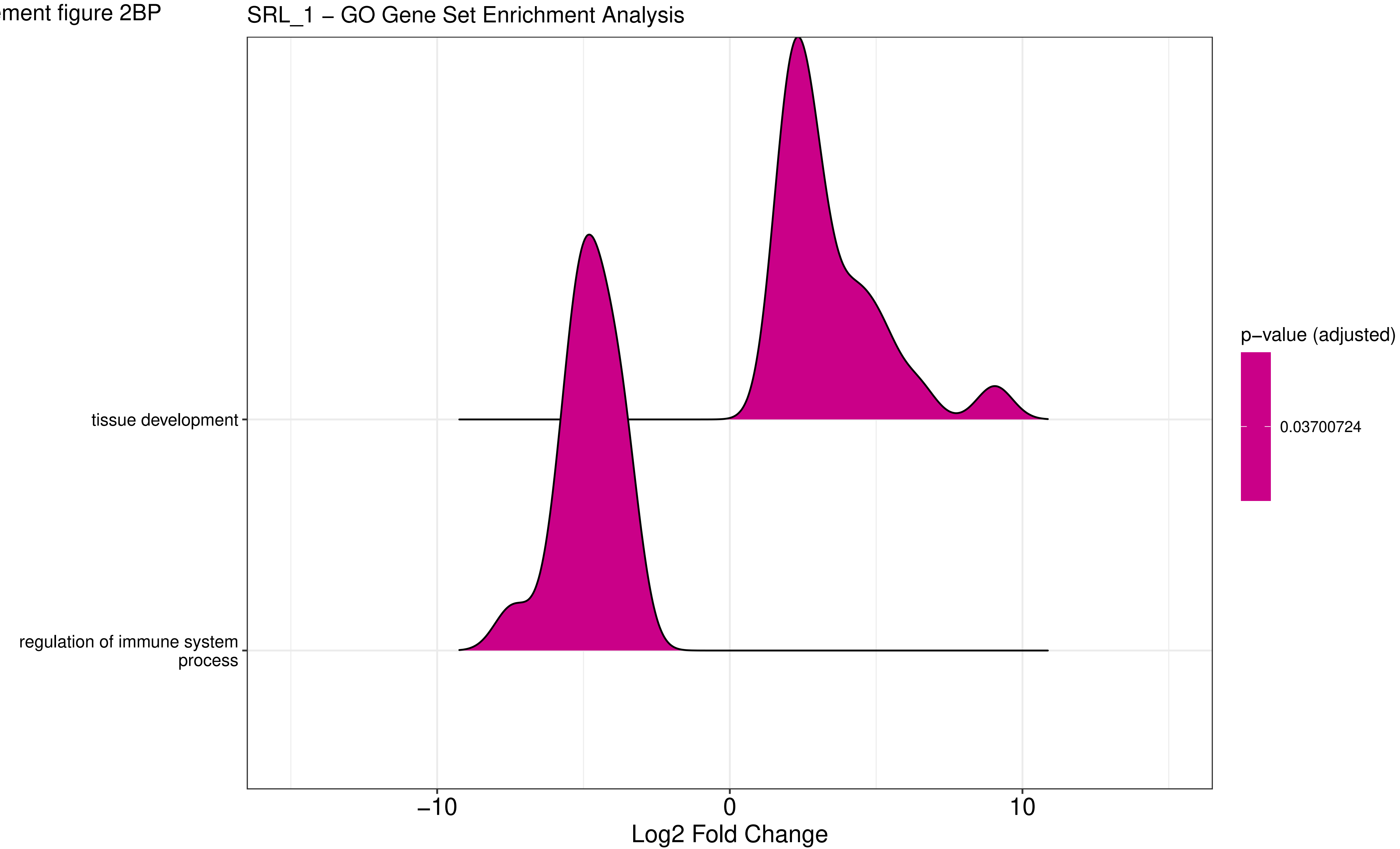

Supplement figure 2BQ

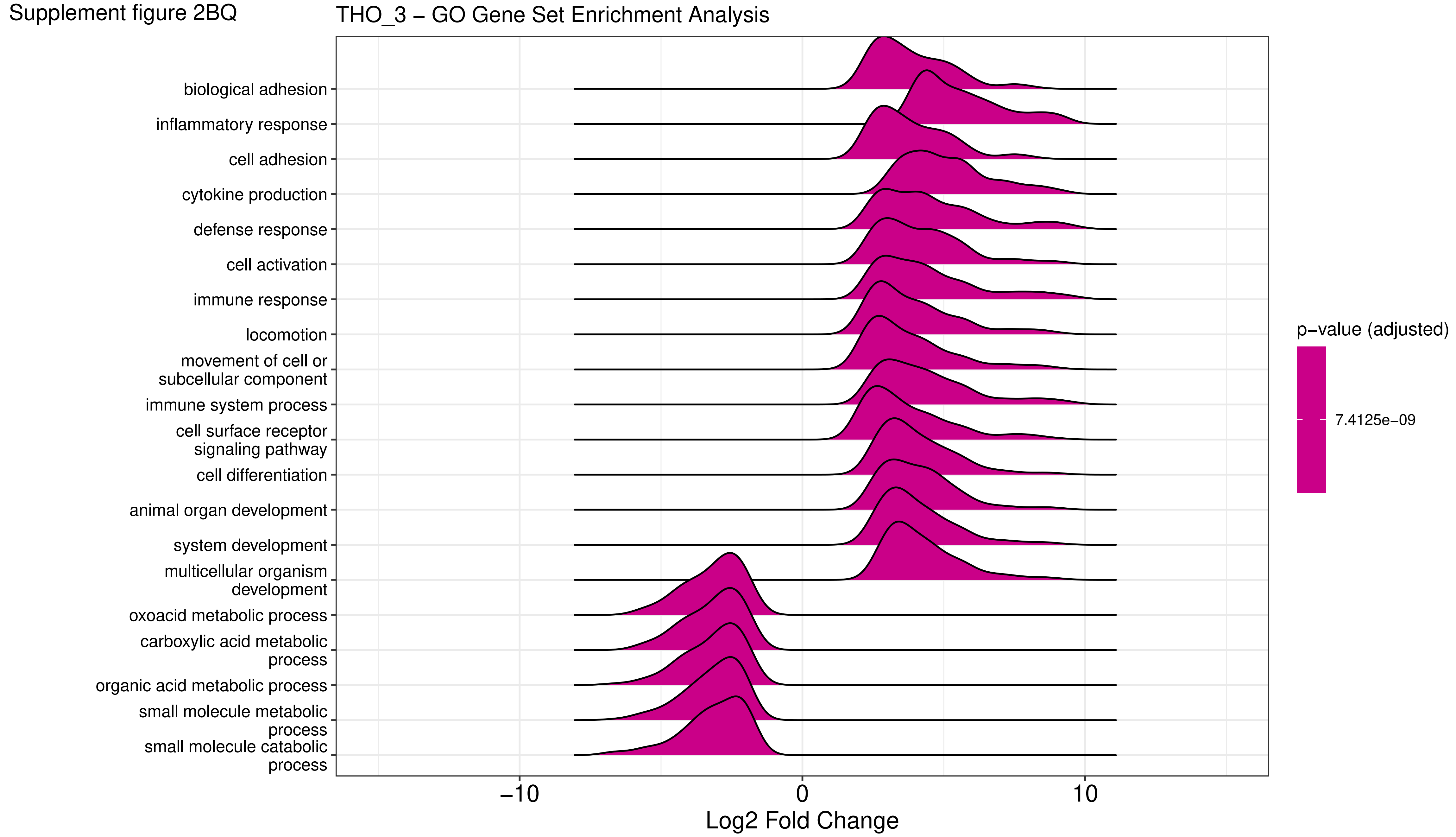

Supplement figure 2BR

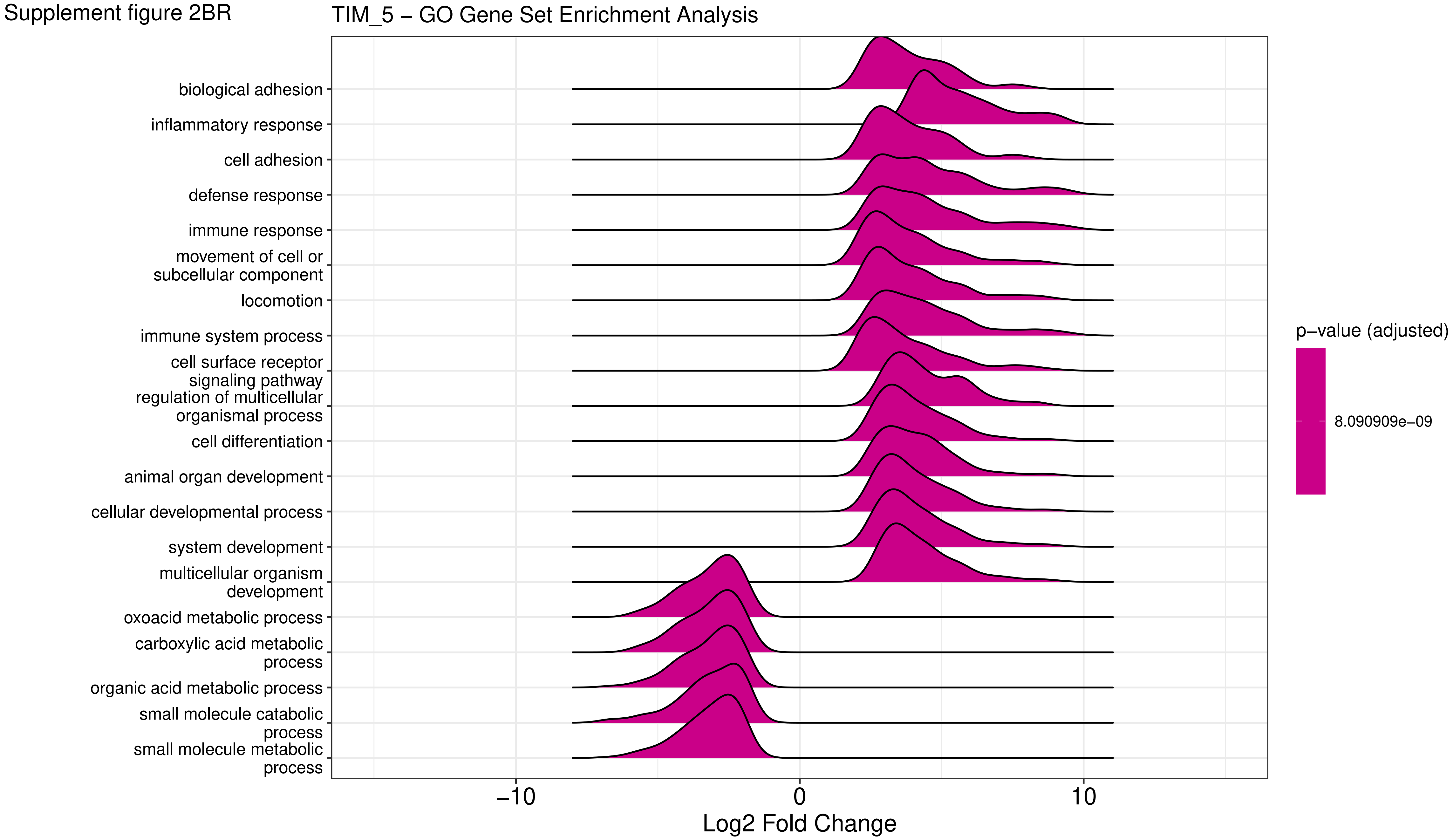

Supplement figure 2BS

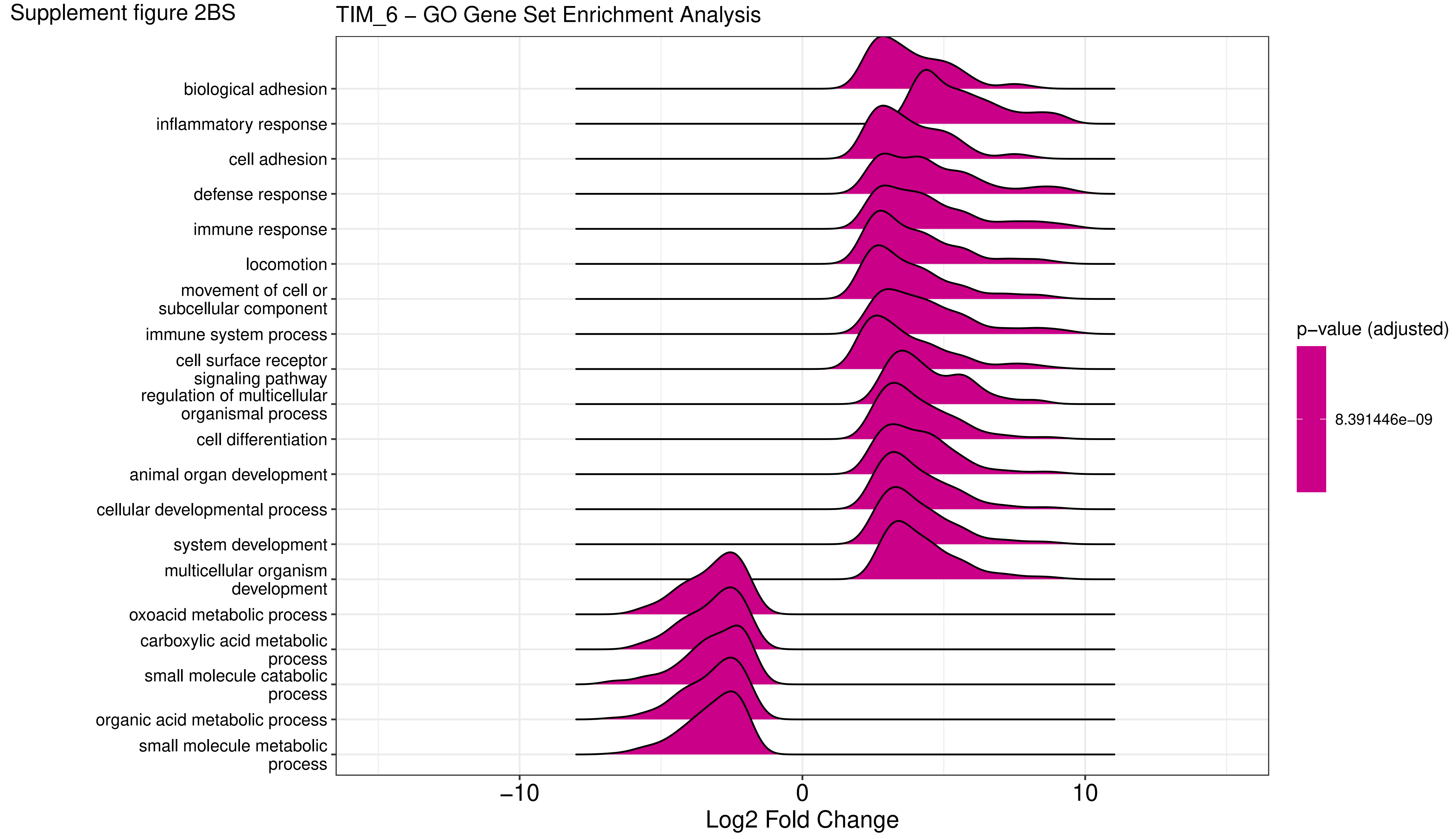

Supplement figure 2BT

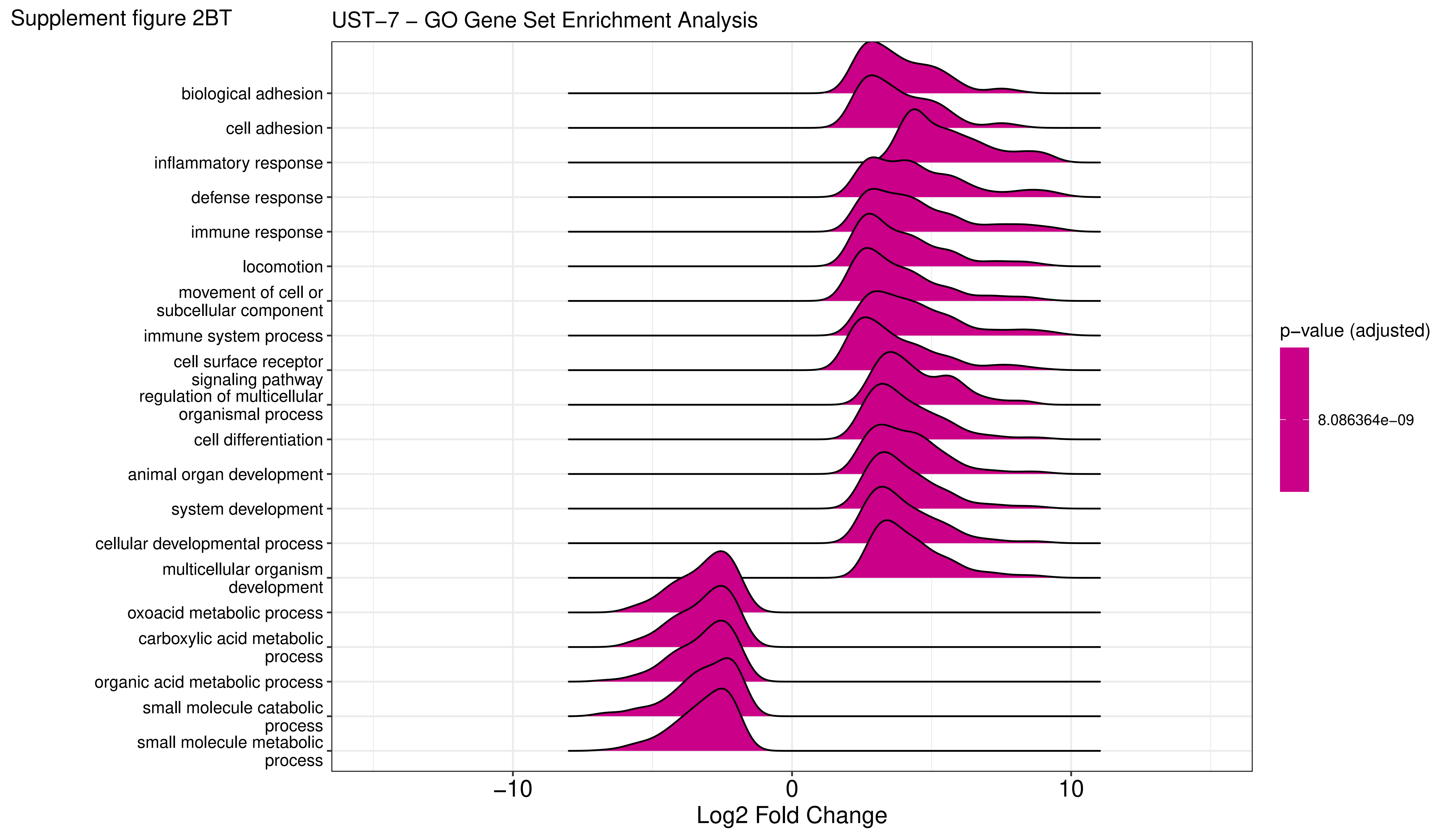

Supplement figure 2BU

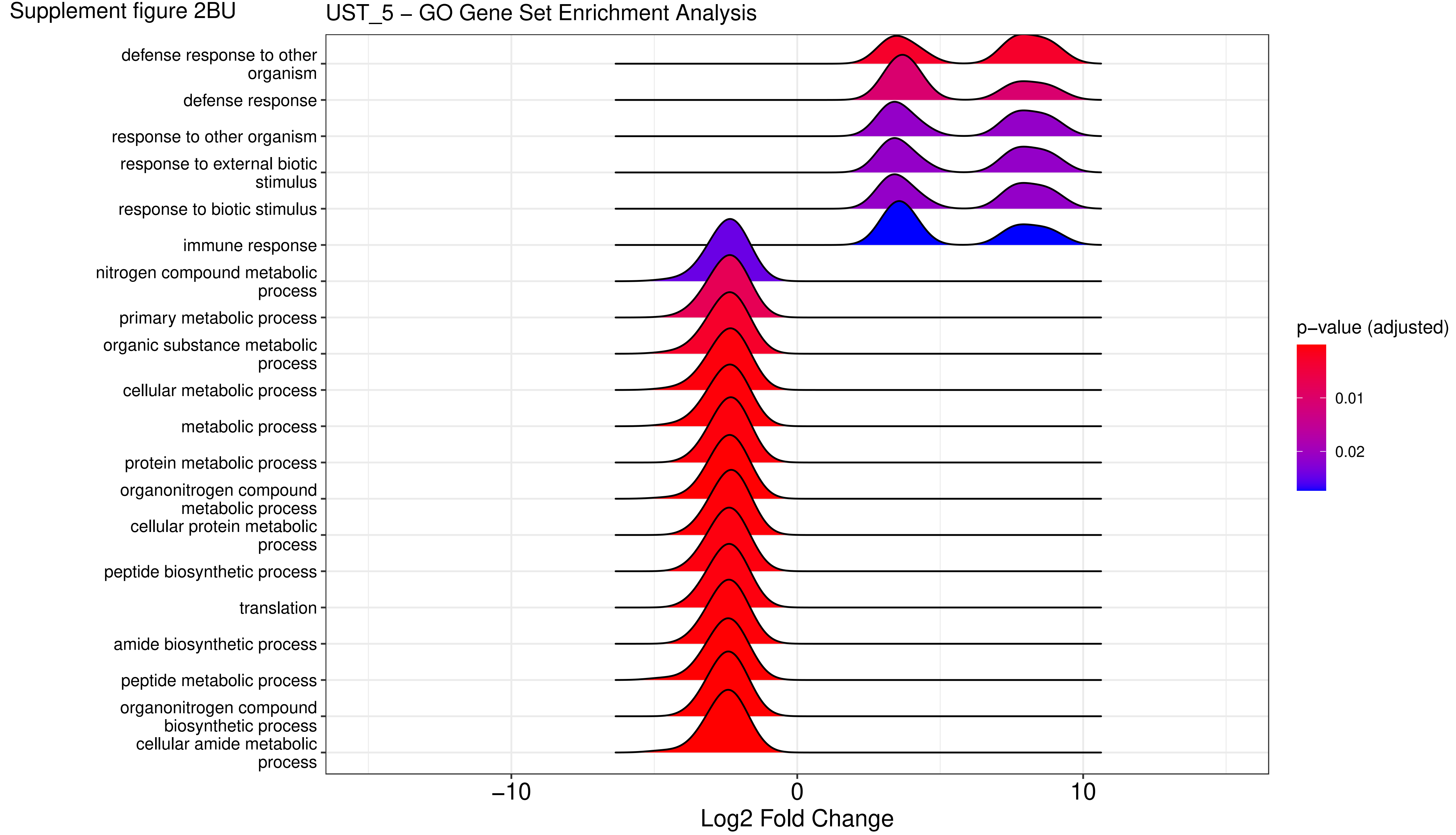

Supplement figure 2BV

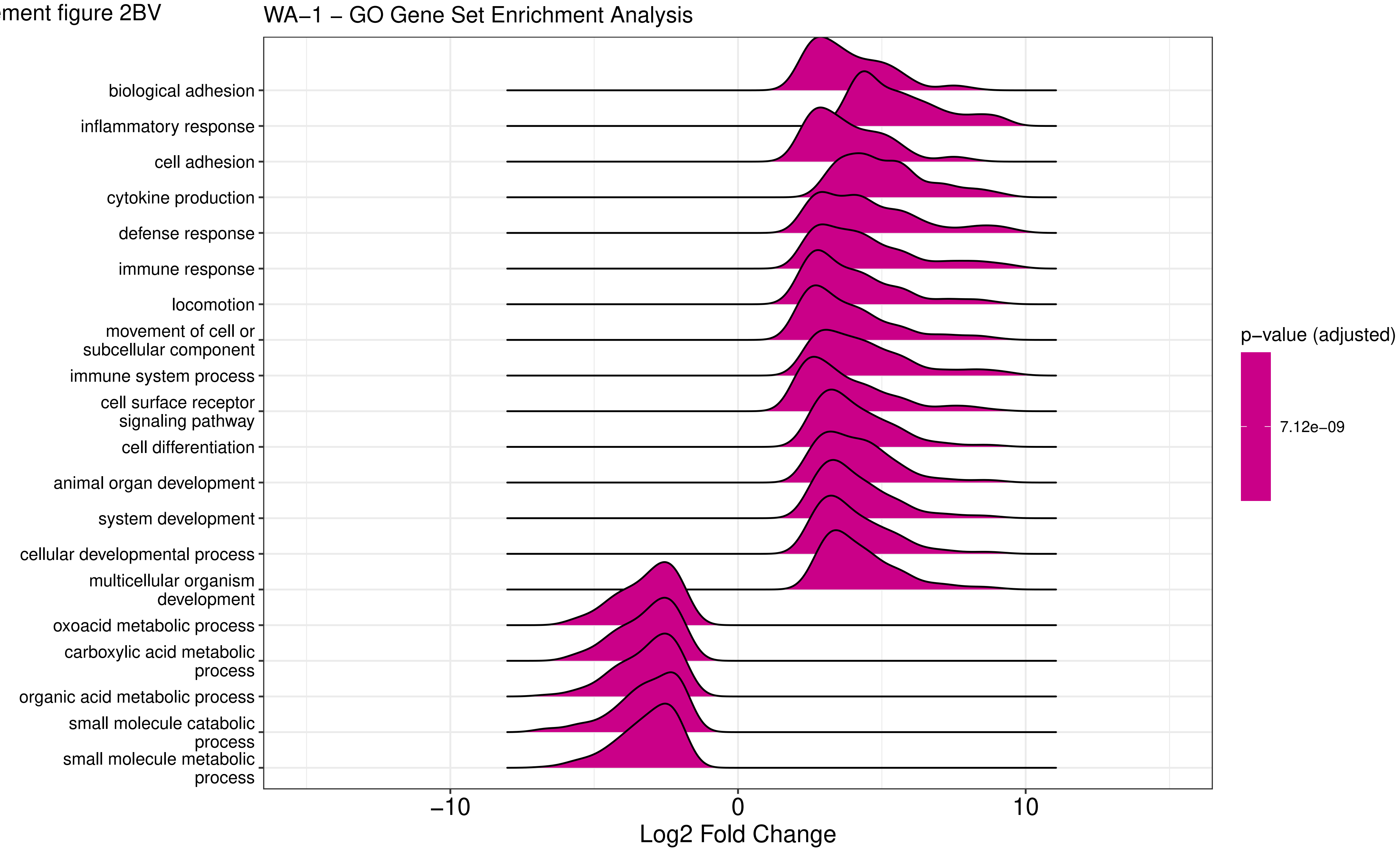

Supplement figure 2BW

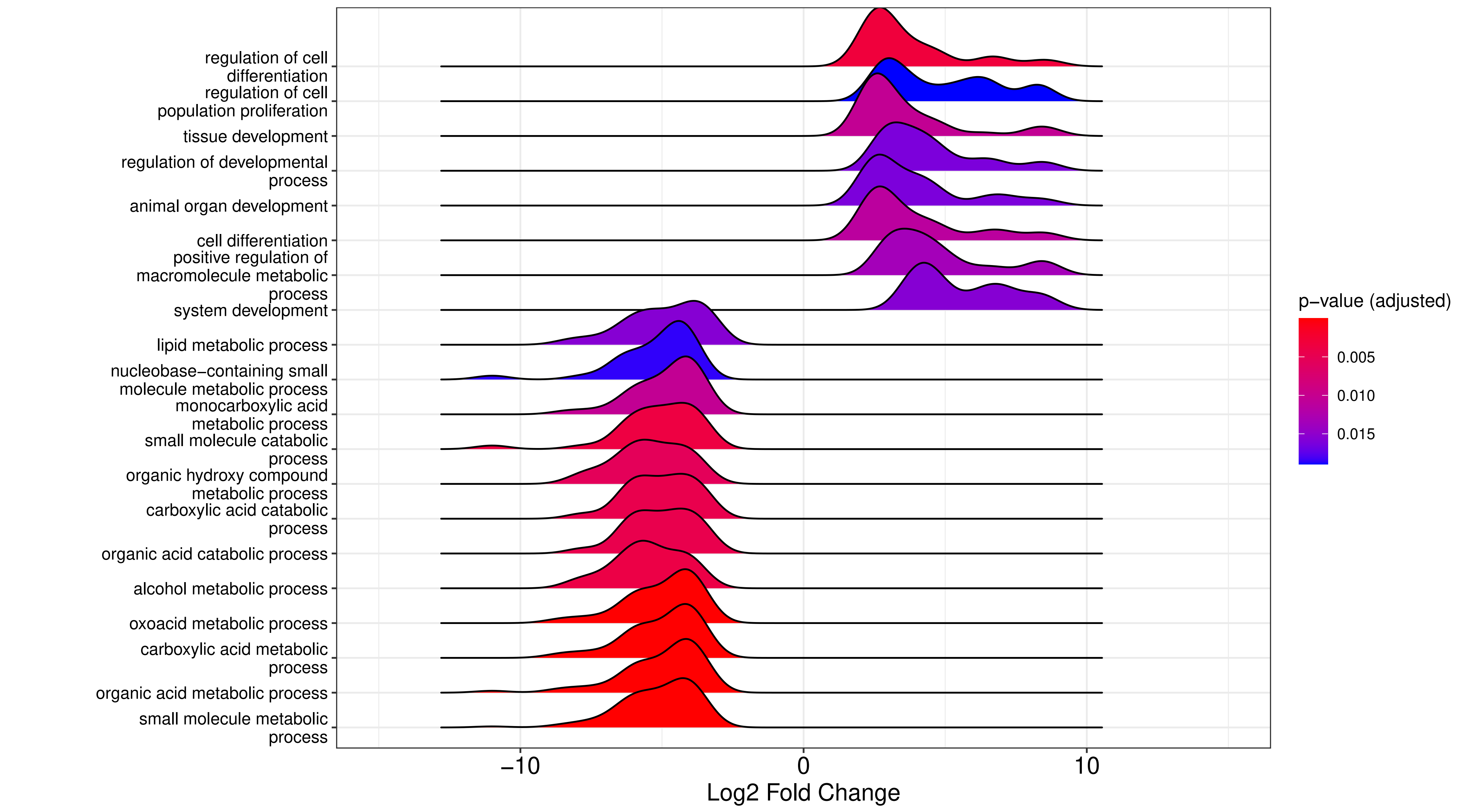

Supplement figure 2BX

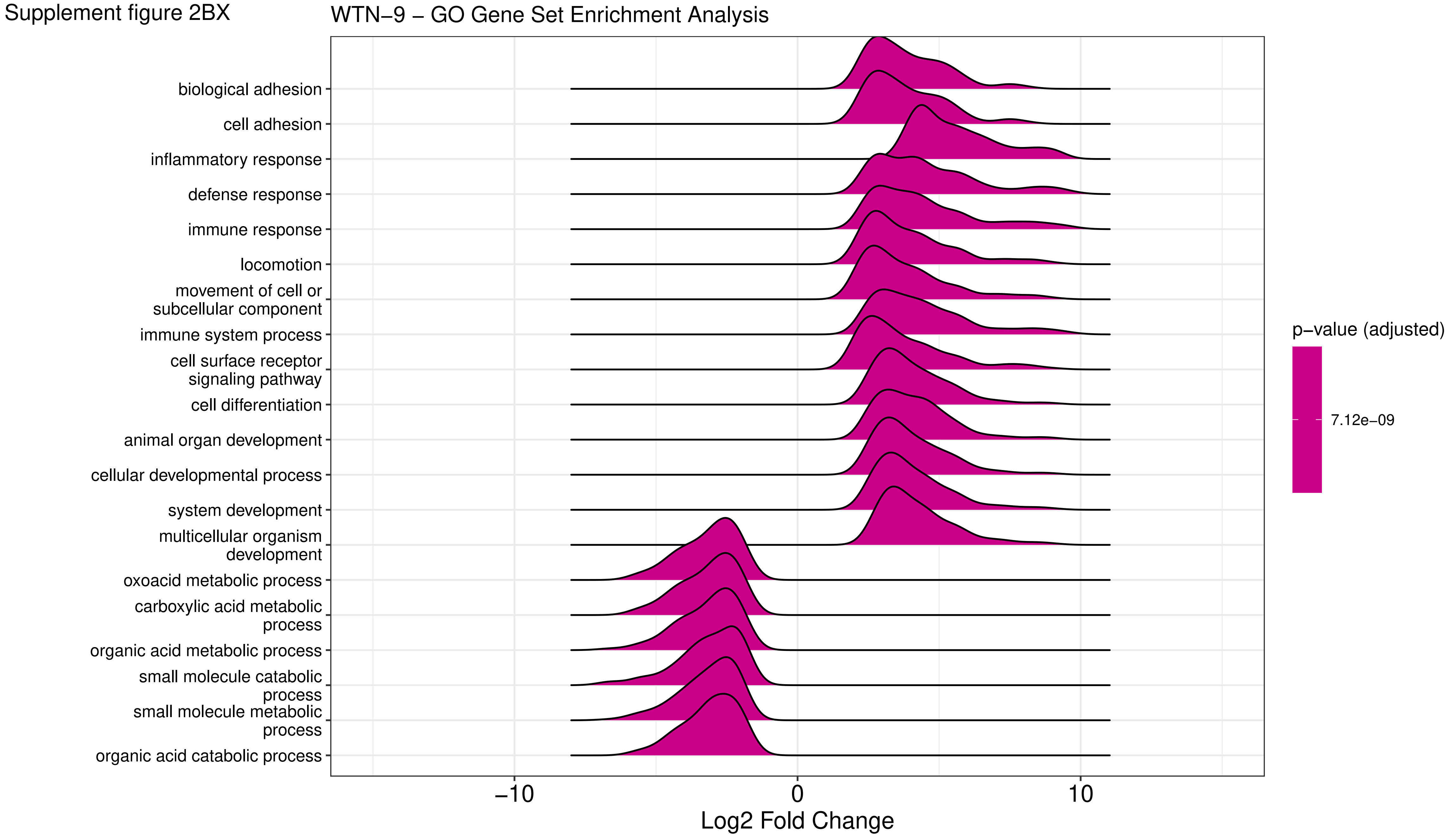

Supplement figure 2BY

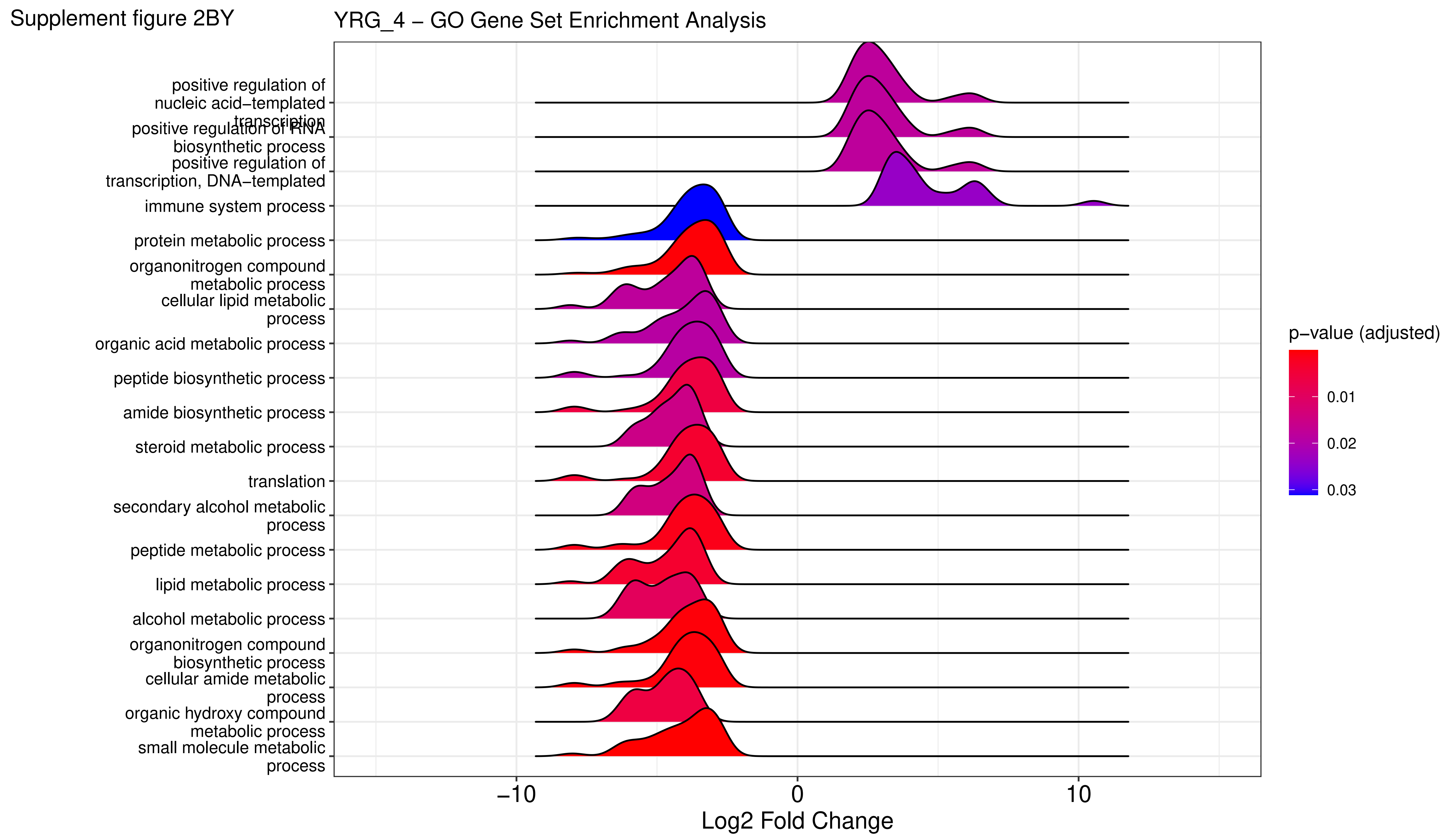

Supplement figure 2BZ

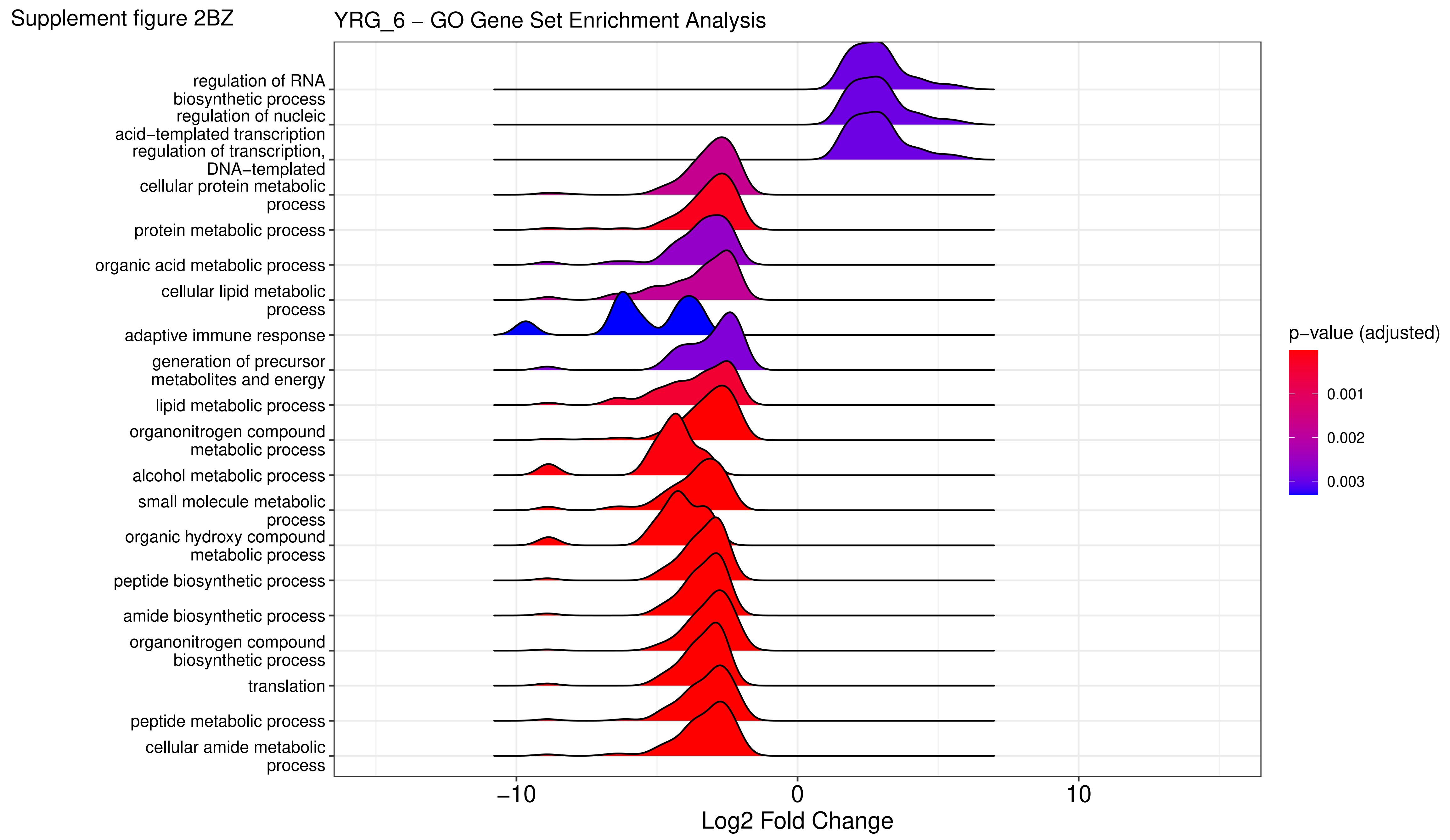
