## Supplementary figure 1 for "Pathogen profiling of Australian rabbits by metatranscriptomic sequencing"

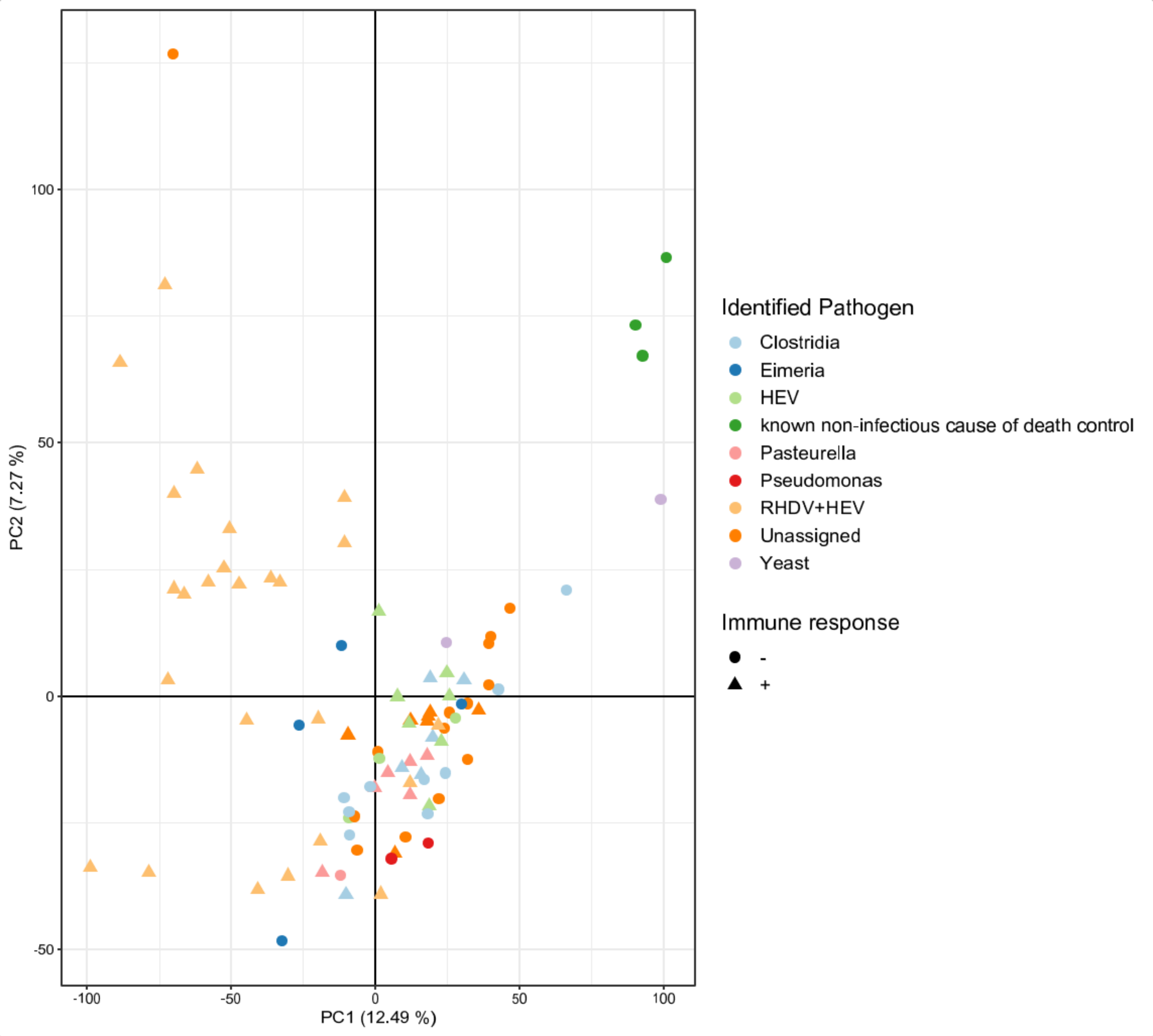


**Supplementary Figure 1:** Principal component analysis (PCA) on normalised counts of the host transcriptome. Colors refer to the identified pathogen. Shapes indicate whether an immune response could be detected via Gene set enrichment analysis. Gene sets were defined by Gene Ontology (GO)-terms. GO-terms considered to indicate evidence of an 'immune response' are listed in Supplementary table 2.
